# High precision in microRNA prediction: a novel genome-wide approach based on convolutional deep residual networks

**DOI:** 10.1101/2020.10.23.352179

**Authors:** C. Yones, J. Raad, L.A. Bugnon, D.H. Milone, G. Stegmayer

## Abstract

**Motivation:** MicroRNAs (miRNAs) are small non-coding RNAs that have a key role in the regulation of gene expression. The importance of miRNAs is widely acknowledged by the community nowadays, and the precise prediction of novel candidates with computational methods is still very needed. This could be done by searching homologous with sequence alignment tools, but this will be restricted only to sequences very similar to the known miRNA precursors (pre-miRNAs). Furthermore, other important properties of pre-miRNAs, such as the secondary structure, are not taken into account by these methods. Many machine learning approaches were proposed in the last years to fill this gap, but these methods were tested in very controlled conditions, which are not fulfilled, for example, when predicting in newly sequenced genomes, where no miRNAs are known. If these methods are used under real conditions, the precision achieved is far from the one published.

**Results:** This work provides a novel approach for dealing with the computational prediction of pre-miRNAs: a convolutional deep residual neural network. The proposed model has been tested on several complete genomes of animals and plants, achieving a precision up to 5 times higher than other approaches at the same recall rates. Also, a novel validation methodology is used to ensure that the performance reported can be achieved when using the method on new unknown species.

**Availability:** To provide fast an easy access to mirDNN, a web demo is available [here](http://sinc.unl.edu.ar/web-demo/mirdnn/). It can process fasta files with multiple sequences to calculate the prediction scores, and can generate the nucleotide importance plots. The full source code of this project is available [here](http://sourceforge.net/projects/sourcesinc/files/mirdnn) and [here](https://github.com/cyones/mirDNN).

## 1 Introduction

MicroRNAs (miRNAs) have a crucial role in post-transcriptional gene regulation, cell growth and other physiological processes. Thus, the discovery of novel miRNAs can be considered one of the most essential problems in computational biology today (Demirci *et al*., 2017). Since their discovery, miRNAs have reshaped the knowledge on gene regulation. They can determine the genetic expression of cells and influence the state of the tissues (Bartel, 2004). Therefore, finding new miRNAs and inferring their functions are necessary tasks for understanding their roles in gene regulation. Furthermore, given their importance in promoting or inhibiting certain diseases and infections, the prediction of new miRNAs is of high interest today (Chen *et al*., 2019; Bugnon *et al*., 2019; Huan and et al., 2015; Takahashi and et al., 2015). For example, in biomarkers developing and targeted drug delivery (Cheng and et al., 2015; Lai *et al*., 2011). However, when evaluating a large number of candidates in a full genome context, experimental methods are unfeasible and computational methods play an important role for their prediction.

One simple way of finding miRNAs in a genome could be using a local alignment search tool, such as BLAST, for finding regions of similarity between sequences. Having known and publicly available miRNAs of other species, similar sequences can be found by direct comparison, that is, similarity-based homology. However, it is known that RNA sequences can be considered context free languages, which means that direct similarity search like this can be very limited (Searls, 2002). For example, it is known that miRNAs precursors (pre-miRNAs) in animals usually have a stable stem-loop structure. A sequence could be relatively similar to a known pre-miRNA, but a small number of mutations could lead to a very different and unstable secondary structure. Another problem is that not all nucleotides are equally important in a pre-miRNA. For example, it is known that the stem region of the sequence is usually highly conserved in comparison to the loop region. A sequence alignment tool would give the same importance to mutations independently of the position on the sequence.

The limitations present in the methods based on alignment leads to the emergence of many machine learning (ML) approaches. The objective of these methods is to learn other intrinsic properties that defines a miRNA besides the raw sequence. But this task has many complex and challenging aspects. In spite of the pre-miRNAs having a typical stem-loop structure with few internal loops, a very large number of hairpin-like structures can be found in a genome. Moreover, the number of known pre-miRNAs, in order to be used as reference or positive class, is quite limited. For example, in the case of human genome, there are 1982 miRNAs deposited in miRBase (Kozomara *et al*., 2018) v22 (release oct 2018), while more than 48 millions of sequences with hairpin-like structure could be extracted from the complete genome. This constitutes a very large class imbalance problem (in the order of 1:24,000). This high imbalance is very difficult to handle by any ML model (Bugnon *et al*., 2019). In spite of the fact that there is a myriad of methods for pre-miRNA classification published (Li *et al*., 2010; Gomes and et al., 2013; Shukla *et al*., 2017; Stegmayer *et al*., 2018; Chen *et al*., 2019), a study has clearly shown that the prediction of pre-miRNAs is yet far-away from being satisfactory solved because existing methods have a very high rate of false positives (Demirci *et al*., 2017). That is, they provide an excessively long list of candidates to novel pre-miRNAs, that cannot be validated with wet-experiments. Moreover, in many of the published methods, the performance measures were obtained in very controlled conditions. For example, many methods were validated through cross validation over the miRNAs of one species, that is, using a high percentage of the miRNAs of the species under study as training example. This is not possible when studying a new sequenced genome, since in many cases there is not any confirmed pre-miRNA. Furthermore, some methods were tested on artificially balanced datasets, which leads to a overestimation of the precision.

Another very important aspect when looking for candidates in a full genome is the feature extraction required for training a classical ML classifier. A large number of features have been proposed for pre-miRNAs mainly based on the structure of the sequences, but they have the serious disadvantage of being dependant on feature engineering, that is, a process highly dependent on manual intervention (Ivani de ON Lopes and Alexander Schliep and Andre de Carvalho, 2014; Yones *et al*., 2015; Raad *et al*., 2019). The emergence of deep learning (DL) models has led to substantial improvements in the field of automatic information extraction and representation, precisely avoiding the feature engineering step (LeCun *et al*., 2015). Deep models can automatically extract relevant features by themselves, differently to the traditional procedure of hand-made definition of the features to extract, which is extremely time consuming and requires the involvement of domain experts. Deep learning is inspired by the representation of biological neural networks and it can be considered today among the best paradigms of ML for classification (Bengio *et al*., 2013). It has already been employed for small-RNA feature extraction (Zheng *et al*., 2019), identification and classification (Amin *et al*., 2019), showing promising results for genomics analysis. In fact, DL models have had recent success in several sequence classification studies (Zeng *et al*., 2016; Bugnon *et al*., 2019) because empirical analyses have shown that deep models can extract motifs from a set of homologous sequences, which are essential for distinguishing among different types of sequences, such as for example miRNA families (Seo *et al*., 2018; Tang and Sun, 2019). In Eraslan *et al*. (2019) authors analyze gaps and challenges for DL in genomics, mentioning the need for more DL-based tools capable of handling the real genome-wide scenario.

In this work we propose mirDNN, a novel pre-miRNA prediction algorithm based on deep learning. It has been designed for finding candidate sequences from genome-wide data. The model is a convolutional deep residual neural network (He *et al*., 2016) that can automatically learn suitable features from the raw data, without a manual feature engineering. It is capable of constructing a model that can successfully learn the intrinsic structural characteristics of precursors of miRNAs within a sequence. The proposal has been tested with several genomes of animals and plants, and compared with state-of-the-art algorithms.

## 2 Deep learning model for microRNA prediction

MirDNN takes as input RNA sequences, their corresponding predicted secondary structure and the minimum free energy (MFE) when folding (Lorenz *et al*., 2016). The input sequence and its secondary structure is represented as a one-hot-encoding tensor of shape 4 × *L* (see Figure 1), where *L* is the maximum sequence length considered. It has 4 rows for the 4 possible nucleotides (A,C,G,U) in each position. Each column represents a nucleotide in the sequence. The tensor size is fixed and completed with zero-padding for sequences of variable length. This tensor can have, in each column, a 1 and three 0s, according to the nucleotide value to represent at each position. For example, if the first nucleotide of the sequence is Adenine, the first element of the column is 1 at the first row and the rest of the column is 0; if the nucleotide is Cytosine, the second element of the column is a 1 and the rest is 0. The secondary structure of the sequence is represented as a tensor of shape 1 × *L*, where the i-*th* element is 1 if the i-*th* nucleotide of the sequence is left-paired (a left parentheses in the RNAfold format), -1 if it is right-paired and 0 otherwise. The distinction between left and right paired nucleotides help to identify bulges and loops in the sequence. These two tensors are concatenated over the first dimension to form a tensor of shape 5× *L*, which is the input tensor to the convolutional neural network (CNN).

**Figure 1:**
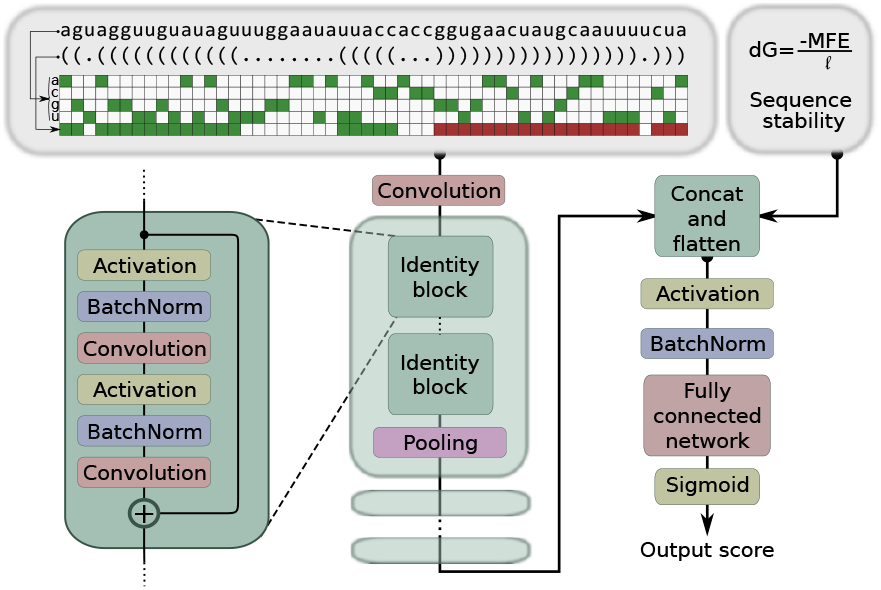
Schematic representation of mirDNN end-to-end architecture. In the matrix, a green cell represents a 1, a red cell represents a -1, and a white cell represents a 0.

The first layer of the network is a one-dimensional convolution of length *F* with *W* filters. This layer generates a sequence of length *L* and *W* features, which is the width of the network in all the next layers. Then, several stages composed of identity blocks (He *et al*., 2016) and pooling layers are stacked. The identity blocks allow the model to auto-define the number of convolutional layers needed during training, which avoids optimization of the number of hidden layers. When there are many identity blocks available, the model is capable of automatically selecting how many of them are necessary while non-necessary blocks are just skipped. Each block is composed of two activation functions, two batch normalization layers, and two convolutions that can be seen in the detail (left part) of Figure 1. All these operations are done over the input tensor and the result is summed up to the input of the next identity block. This helps to back-propagate the error during training, allowing the addition of more convolution layers without difficulting the training of the model. After *B* identity blocks, a pooling layer is used to reduce the length *L* of the sequence by a factor of 2. After *M* of these stages, a tensor of shape *W* × *L/*2*M* is obtained. This tensor is converted in a one-dimensional vector, and then it passes through activation and batch normalization layers. Then, the input sequence stability, *DG*, is appended in order to form a tensor of *WL/*2*M* + 1 elements. It is calculated as *DG* = *MFE/*ℓ, where ℓ is the original sequence length (not *L*, which is the zero-padded one). After that, this tensor feeds a fully connected network that generates the corresponding output score for prediction of the input sequence.

For training the mirDNN model, two strategies were tested to tackle the imbalance present in genome-wide data. The first one was positive class oversampling, that is, each batch is built with a proportion of positive examples that is higher than the real proportion present in the dataset. To achieve this, the positive examples are sampled with replacement. The second strategy tested was focal loss (FL), proposed by Lin *et al*. (2017). When the negative examples (the majority class) are classified, they generate an error to be back-propagated through the model. Due to the high imbalance, the sum of these contributions is much larger than the contribution of the (few) positive examples, and the model is heavily biased towards the negative class. Thus, it does not learn the positive class correctly. In order to solve this, a FL function is defined as

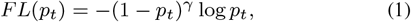

where *pt* is the predicted probability (output score) for the sequence under analysis, and the parameter *γ* can be used to increase or reduce the weight given to the examples correctly classified. This way, if the majority samples are correctly classified (with a small label error), the error back-propagated is much smaller than with traditional binary cross entropy loss (BCE). Therefore, in an imbalance escenario the model errors for the minority (in this case, the positive) class get more importance and drive the learning of the network.

## 3 Materials and experimental setup

### 3.1 Data

Genome-wide data from 4 species was used in this work: *Arabidopsis thaliana, Caenorhabditis elegans, Anopheles gambiae*, and a much more larger dataset with the sequences of the *Homo sapiens* genome. First, HextractoR1 was used to extract all the stem-loop sequences from each genome with standard settings, with window size of 320 nt for plants and 160 nt for animals. The positive examples were taken from mirBase v22. Only the animal sequences were used on test with animal genomes and only the plant sequences were used on the test with *A. thaliana*.

### 3.2 Experimental setup

In order to have an experimental setup that emulates a real situation in genome-wide, for example the case of the genome of a recently sequenced species, all the pre-miRNAs of each genome were excluded from the corresponding training sets. This is a very challenging test condition, a much harder problem to solve than usually published experimental setups. Thus, training and testing follow a leave-one-species-out scheme, presented in Figure 2. For training mirDNN with a full-genome, the positive set has the pre-miRNAs of the rest of the species of the kingdom, without the specific pre-miRNAs of the genome under analysis. The negative-class training set has the 95% of the stem-loop sequences extracted with HextractoR, within the genome under analysis. It is important to point out that the negative training set might probably have pre-miRNAs, but these are left because in a real case escenario false positives cannot be filtered out from the training set. For testing, the specific known pre-miRNAs of the species under analysis are used, together with the remaining 5% of stem-loops of the genome. In this last testing partition, the known pre-miRNAs (false negatives) are filtered in order to accurately measure the performance of the methods. The total number of sequences of each train and test partition is reported in Table 1.

**Table 1:**
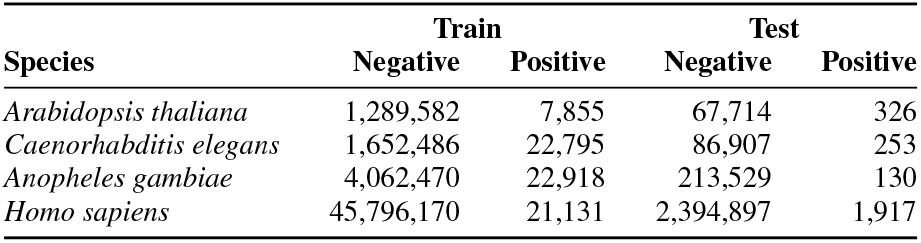
Number of sequences in train and test partitions of each genome, using the leave-species-out validation schema.

**Figure 2:**
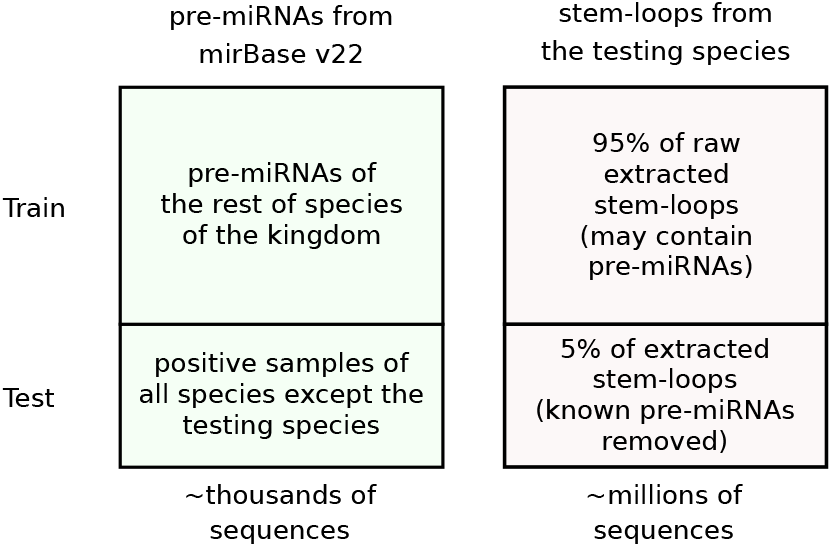
The leave-one-species-out scheme for training and testing the mirDNN model.

To select the optimal hyper-parameters for mirDNN, a grid search was performed using the data of three full genomes: *A. thaliana, C. elegans* and *A. gambiae*. The training sequences of each genome were splitted in a 90%-10% training-validation scheme to measure the performance of each hyper-parameter set. The hyper-parameters tuned were: the number of filters *W* (16, 32, 64), the number of identity blocks *B* on each stacked stage (1,3,5), the filter length *F* (3, 5), and the loss function (BCE or FL). Also, two imbalance conditions were tested during training: i) oversampling the minority class; ii) using the training data with the natural imbalance.

In Figure 3, the effect of different parameters over the AUCPR is shown. First, Figure 3 a) shows the effect of different alternatives during the training process. From the figure, it can be concluded that over-sampling the positive class generates a significative fall in the AUCPR. Training the model with the natural class imbalance of the dataset leads to the best results. With respect to the loss function, using FL achieves the best results in all cases. Secondly, in Figure 3 b) the AUCPR of different architectures using FL and without oversampling is shown. Here, the differences are not very conclusive: there are several good pairs of parameters. In *C. elegans* and *A. thaliana* the best results were achieved with *B* = 3 and *W* = 64, while in *A. gambiae* this configuration reached the second best AUCPR. Therefore, these values will be used in the final model. Finally, in Figure 3 c) the effect of filter size *W* on AUCPR (left axis) in the three datasets using FL and without oversampling is shown. As it can be seen, larger filters do not help improving the results, and since filters of length 5 are computationally intensive, filters of length 3 were chosen.

**Figure 3:**
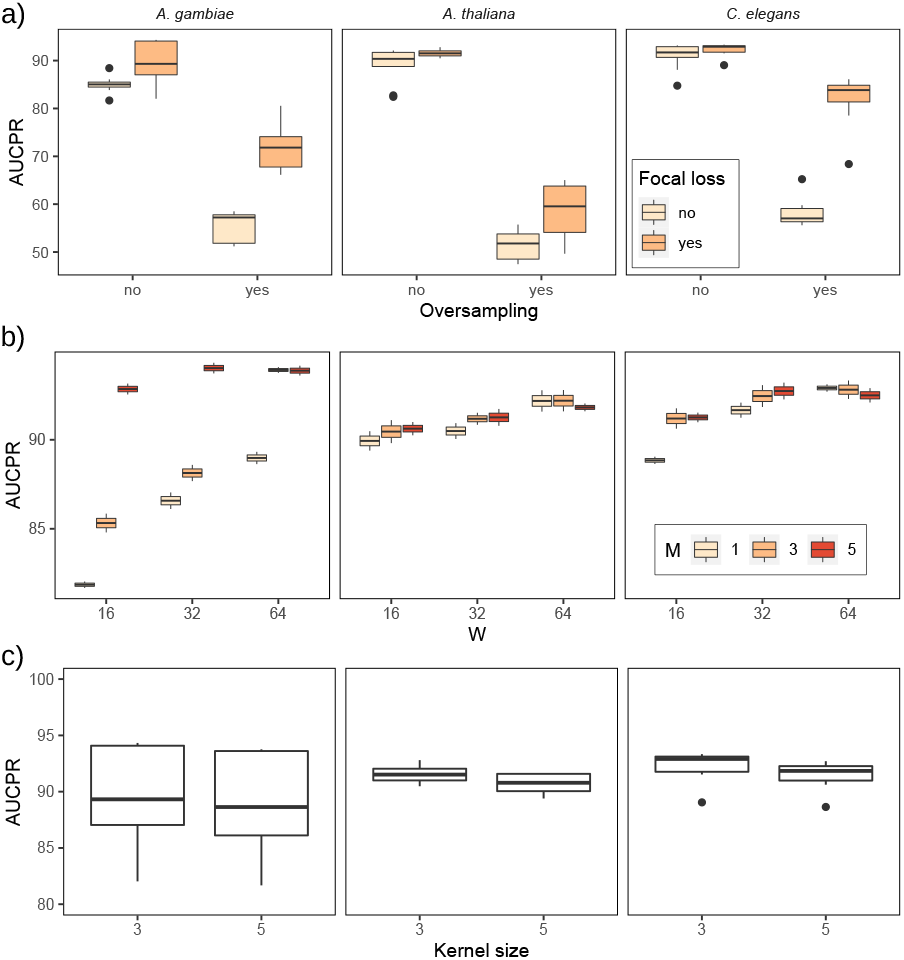
Effect of different hyper-parameters sets in the AUCPR on the 3 genomes: a) effect of focal loss and oversampling; b) different combinations of filters W and number of identity blocks B, using FL without oversampling for training; c) effect of increasing the size of the filters.

The mirDNN model has been compared against the following methods: i) BLAST alignments against complete pre-miRNAs and using the e-value reported for each sequence as a prediction score (BLAST version 2.6 was used with default parameters); ii) DeepMir (Tang and Sun, 2019) a CNN networks for miRNA family prediction, which obtained one of the best performances in a very recent review of genome-wide methods (Bugnon *et al*., 2020); iii) miRNAss (Yones *et al*., 2017), a semi-supervised ML prediction method specifically designed for taking advantage of unlabeled sequences even when there are just very few positive examples; iv) a classical Random Forest (RF) as representative of supervised model for pre-miRNAs prediction (Gudy *et al*., 2013), which was among the best supervised models according to (Stegmayer *et al*., 2018); and v) a Gradient Boosting Machine (GBM), which is currently one of the best methods based on ensemble learning (Ke *et al*., 2017).

The full source code to reproduce the experiments is available here. Furthermore, to provide fast an easy access to mirDNN, a web demo (Stegmayer *et al*., 2016) is available here.

### 3.3 Performance measures

The prediction quality of the model was assessed using the classical classification measures of precision (*P*) and recall (*R*), defined as

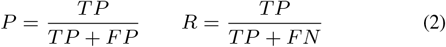

where *T P, FP* and *FN* are true positive, false positive and false negative predictions, respectively. This measures were used to generate precision recall curves (PRC), which is a well-known performance indicator. It has been shown (Saito *et al*., 2015) that this measure is preferred over the classical receiver operating characteristic (ROC) curve to assess binary classifiers with highly imbalanced data. When there is a large class imbalance in a dataset, a classifier can reach a good performance in terms of specificity, but can perform poorly in providing good quality candidates, with a large amount of false positives. A PRC can provide a better assessment of performance because it also evaluates the fraction of true positives among the total positive predictions. The area under the precision-recall curve (AUCPR), which is a single numeric summary of the information, will also be reported.

## 4 Results and discussion

### 4.1 Comparison with state-of-the-art methods

Figure 4 shows the PRC plots corresponding to mirDNN and the other methods for *A. thaliana, C. elegans, A. gambiae* and *H. Sapiens* full genomes. In these plots, precision is shown in the y-axis and recall in the x-axis. The curves are generated varying the threshold applied to the output score, to separate the positive from the negative samples and calculating the precision and recall for each possible threshold.

**Figure 4:**
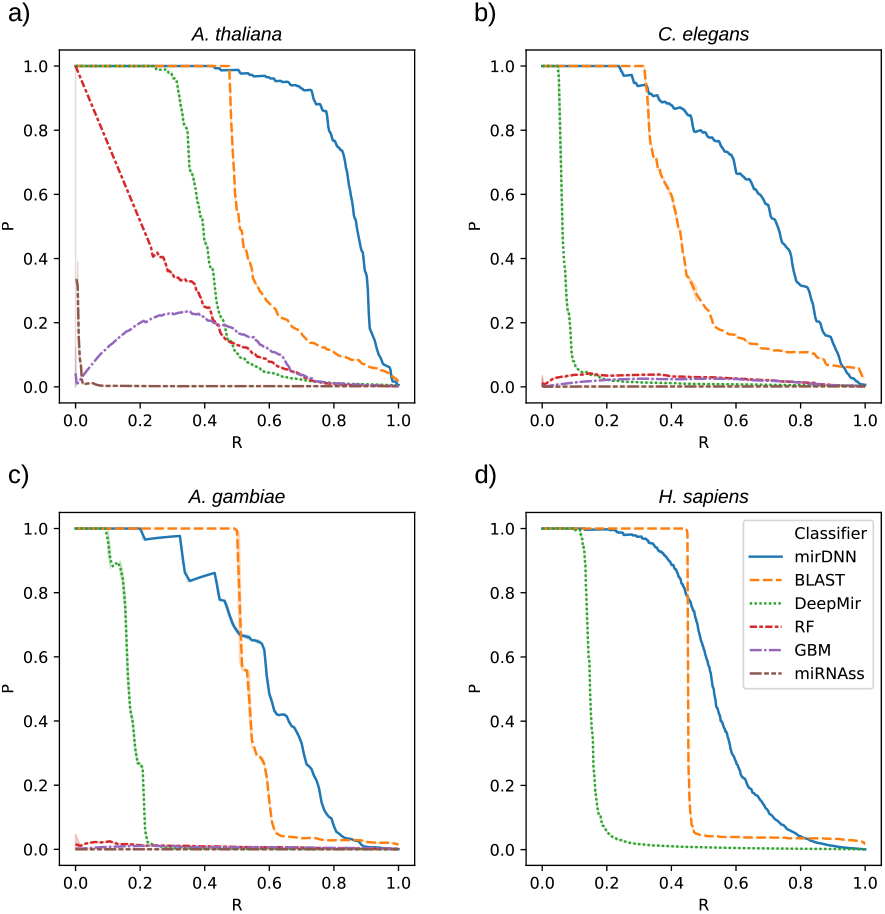
PRC plots for mirDNN on different genomes. a) *A. thaliana*, b) *C. elegans*, c) *A. gambiae*, d) *H. Sapiens*.

With the *A. thaliana* full genome, mirDNN provides a precision of 0.93 and, at the same time, a recall of 0.70. BLAST reaches a precision of 1.00 up to a recall of approximately 0.50, where it abruptly decreases performance. DeepMir presents a similar behaviour but with a lower performance. The feature-based ML methods obtained the worst results: RF has an almost linearly decreasing behavior with a much worse performance than mirDNN and BLAST; miRNAss and GBM barely reach a precision of 0.40 for low recall rates, therefore they are not competitive. For a recall of 0.70, mirDNN has a precision nearly 5 times higher than the second method, BLAST.

In *C. elegans* the best performing methods are mirDNN and BLAST, while DeepMir, RF, GBM and miRNAss have very low performances. BLAST obtains a very good precision, but only for the positive identical sequences, up to a recall of 0.40, approximately. Above that value, the precision falls. MirDNN has the same precision for the identical sequences, but also maintains a good precision for recalls up to 0.80.

In the case of *A. gambiae*, BLAST again has shown high precision for sequences that are extremely similar to the positives class samples (known pre-miRNAs). However, for a recall of about 0.60, precision rapidly decreases to a very low value (under 0.15) value. DeepMir again presents a similar behavior, but with a much lower performance. Meanwhile, mirDNN can maintain a precision of 0.67 for a recall of 0.60. That is, with BLAST there are 152 TP and 769 FP, while mirDNN provides a list of 152 TPs with only 75 FPs.

In order to see the scalability of mirDNN with a larger dataset, it was tested on *Homo sapiens* full genome. Here, mirDNN was compared with BLAST and DeepMir, because the other methods were highly outperformed in the other three genomes. As it can be seen, DeepMir achieve a good precision but only for recall below 0.20. BLAST achieves better results, with good precision for recall smaller than 0.45,

which could be the approximate size of the set of pre-miRNAs that have very similar sequences in other species. But for recalls higher than 0.45 the precision abruptly falls to very small values. Differently, mirDNN maintains a relatively good precision for recalls up to 0.60. This means that, for example, with a precision of 0.50, mirDNN founds 1,016 true pre-miRNAs while BLAST founds only 863. These results emphasizes the fact that the proposed method is really useful for discovering new pre-miRNAs, while the best competitor only allows finding sequences very similar to those already known.

Finally, a summary of results with all genomes is presented in Figure 5, comparatively showing the AUCPR for all the methods methods. It can be clearly seen here that mirDNN has the best results in all genomes, no matter the species. This highlights the capability of the deep model for analyzing the complete sequence, exploiting both sequential and structural information, and without the need of hand-crafted features. The second best was BLAST, which has shown better performance than the other classical ML methods. This can be explained due to the fact that many pre-miRNAs are highly conserved in many species. This gives BLAST very good precision for these conserved sequences, and also explains why the precision for BLAST abruptly falls when trying to predict pre-miRNAs that are not very conserved across species. In the case of DeepMir, in spite of being a CNN that learns features directly from the sequence, its performance is much lower than the achieved by mirDNN. This difference could be due to several reasons. First, the lower number of layers and a simpler architecture, without identity blocks, such as those of mirDNN. These blocks allow the model to auto-define the number of convolutional layers needed during training, avoiding optimization of this critical hyper-parameter. Second, the lack of a strategy to tackle the class imbalance during training in comparison to the positive class oversampling and focal loss of mirDNN, which allows obtaining a better performance in a real genome-wide scenario. The other ML methods, in spite of using features calculated over the full sequence and considering the secondary structure, fall behind the other methods in terms of performance. This could be mainly attributed to the fact that automatically learned features are more flexible and discriminant than hand crafted features. The convolutions are able to learn most of the discriminative patterns found in the sequence/structure for each species, while the hand crafted features are fixed for all of them.

**Figure 5:**
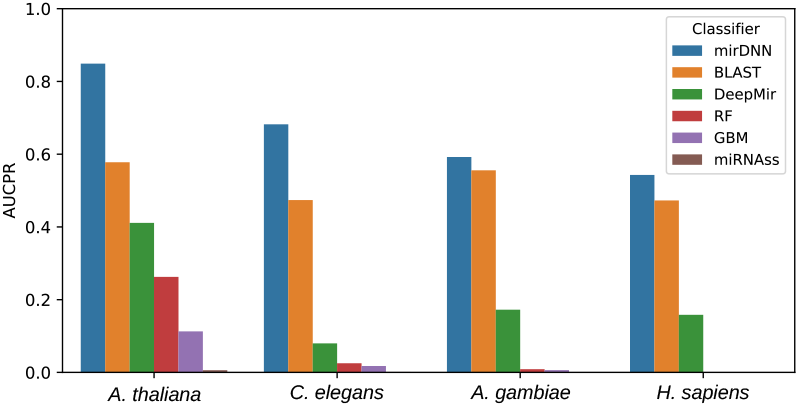
Comparative AUCPR of mirDNN on the four test genomes.

### 4.2 Class activation maps

An interesting insight regarding the deep learning model would be to know which characteristics of the pre-miRNAs were learnt. This could provide useful details on which patterns are important in a sequence to produce a miRNA, using the information learnt from a big database of known pre-miRNAs (for example, the complete animal pre-miRNAs deposited in mirBase v22). Given the number of parameters of the model (321,315), it would be difficult to obtain interpretations of the predictions directly from those. While the first layers learn patterns related to the prediction tasks, these patterns are then heavily processed by the deepest layers of the model. Therefore, it is important to study the inner behaviour of the model, as a whole.

To achieve this, the importance of each nucleotide in the prediction task was measured. First, the whole input sequence was evaluated with the trained mirDNN model, and the prediction output was used as reference score for the sequence. Then, all nucleotides were masked one by one, that is, the corresponding column was converted to a all-zeros vector, and each masked version of the sequence was evaluated again with mirDNN. Thus, the decrease in the score for the sequence was used as a measure of each nucleotide importance. Since the mature miRNA is encoded in a 21nt sequence, generally found next to the terminal loop, it could be expected that convolutional layers pay particular attention to this section of a sequence.

Figure 6 presents the importance that mirDNN assigns to each nucleotide in six sample sequences of known pre-miRNAs. The vertical axis shows the importance levels of each nucleotide, while the horizontal axis shows the nucleotides of the sequence. In addition, the area of the sequence where the corresponding mature miRNA is located has been highlighted. As it can be seen, in the region where the mature is located, the importance values are higher. Figure 6.a) shows how mirDNN gives more importance to the location of the mature in the 5p region. It can be seen in Figure 6.b) how the network assigns more importance to the nucleotides that are in the exact 3p region that is known to encode the mature. In particular here, the nucleotides that encode the seed within the mature are those that present the highest level of importance. A similar importance to the mature region can ben seen in Figures 6.c) and 6.d) for another species. In Figure 6.e) and 6.f) are presented two examples of pre-miRNAs that have two mature regions each. In these examples it is clearly shown that both mature regions 3p and 5pm are recognized and given high importance by mirDNN. To access the statistical significance of these result, the distributions of nucleotide importances among all sequences were compared, through a t-test with the distribution of those nucleotides that do not lay in the mature region. The difference has shown to be statistically significative with a p-value of 8.3E-13, which shows that this trend is present in all the pre-miRNAs used for tests. The nucleotide importance plots are provided for all test sequences in the Supplementary Material.

**Figure 6:**
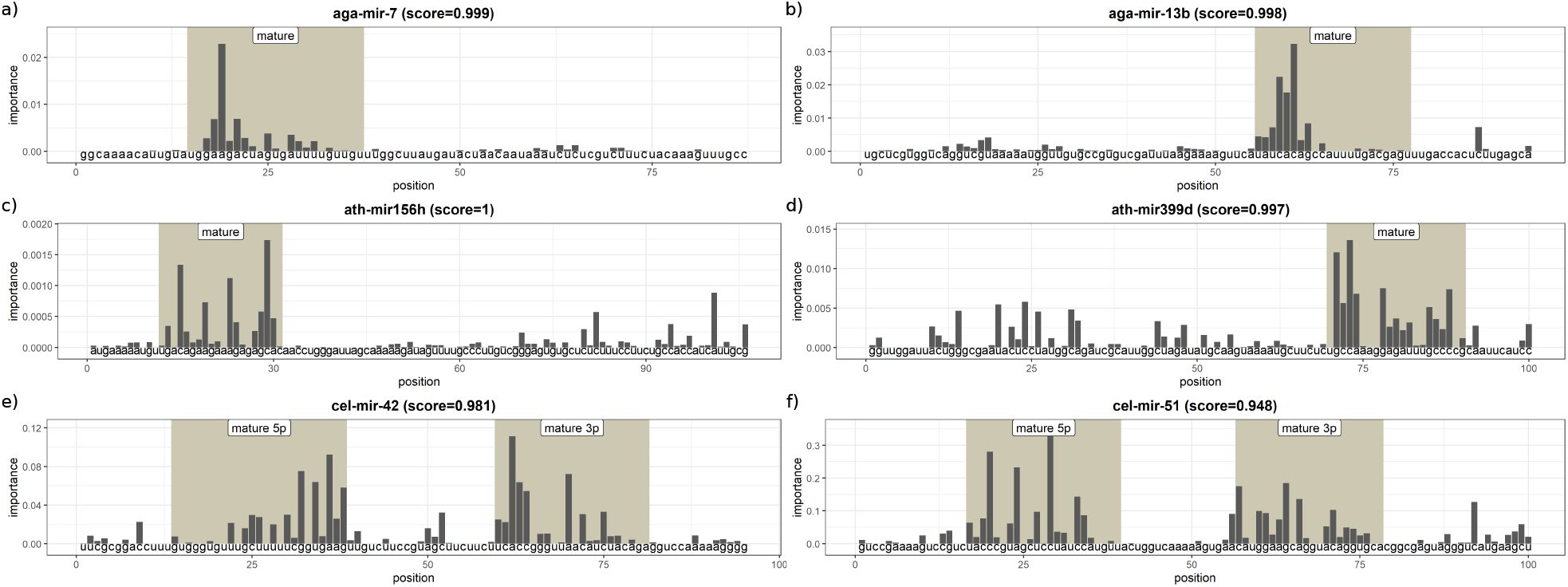
Nucleotide importance of six known pre-miRNA sequences. a) aga-mir-7, b) aga-mir-13b, c) ath-mir156h, d) ath-mir399d, e) cel-mir-42, f) cel-mir-51.

## 5 Conclusions

This work provided a novel approach based in convolutional deep residual networks for dealing dealing with the computational prediction of pre-miRNAs. The comparative results obtained show that this novel approach can achieve a better performance than other machine learning and sequence alignment methods in several genomes. It has been shown that this deep learning model is capable of using both the sequential information and the structure of the sequences, which has significantly improve the predictions in comparison to feature engineering based methods. Also in this work, a validation methodology closer to a real prediction task has been used to test the model, achieving very high precision. Additionaly, a method to generate nucleotide importance values was presented, which can be used to get insights about the newly discovered pre-miRNA candidates. In summary, it can be stated that the inference of pre-miRNAs with mirDNN is fast and scalable, making it suitable to process whole genomes.

## Acknowledgements

The authors acknowledge the support of NVIDIA Corporation with the donation of the Titan Xp GPU used for this research.

## Footnotes

1 https://cran.r-project.org/web/packages/HextractoR

