## Supplementary figures and images for "High precision in microRNA prediction: a novel genome-wide approach based on convolutional deep residual networks"

### aga-mir-7.png

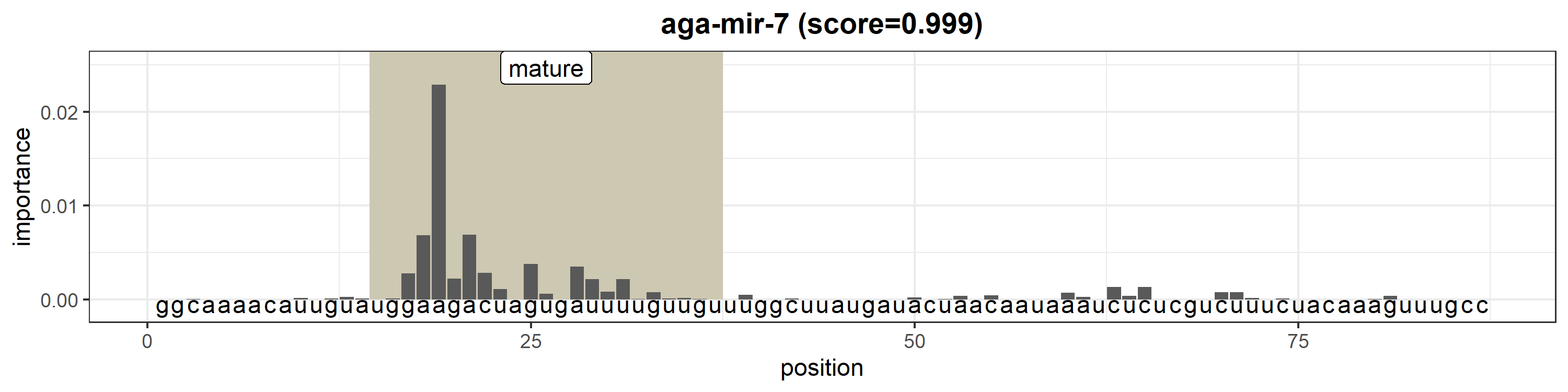

### aga-mir-8.png

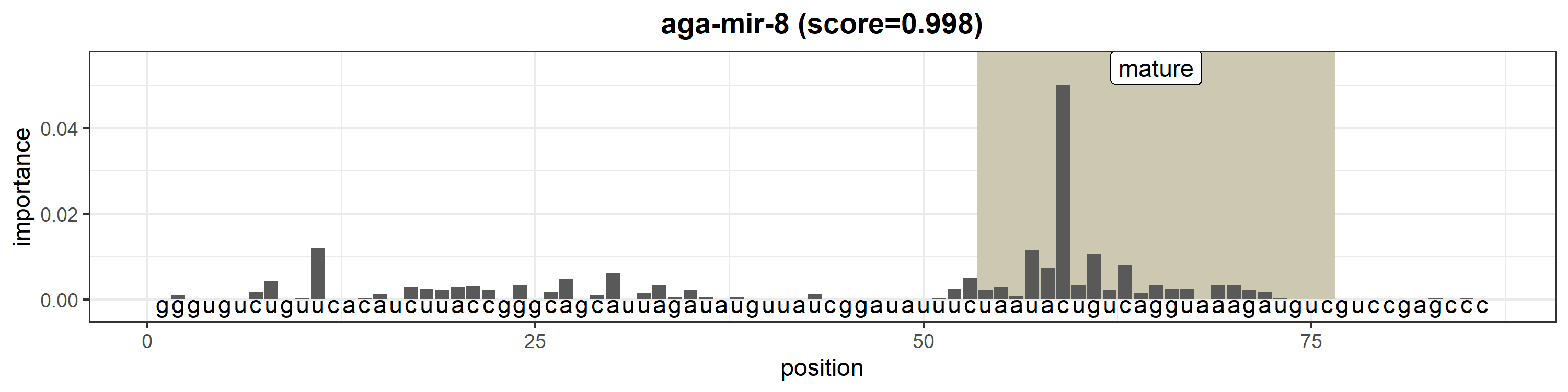

### aga-mir-9a.png

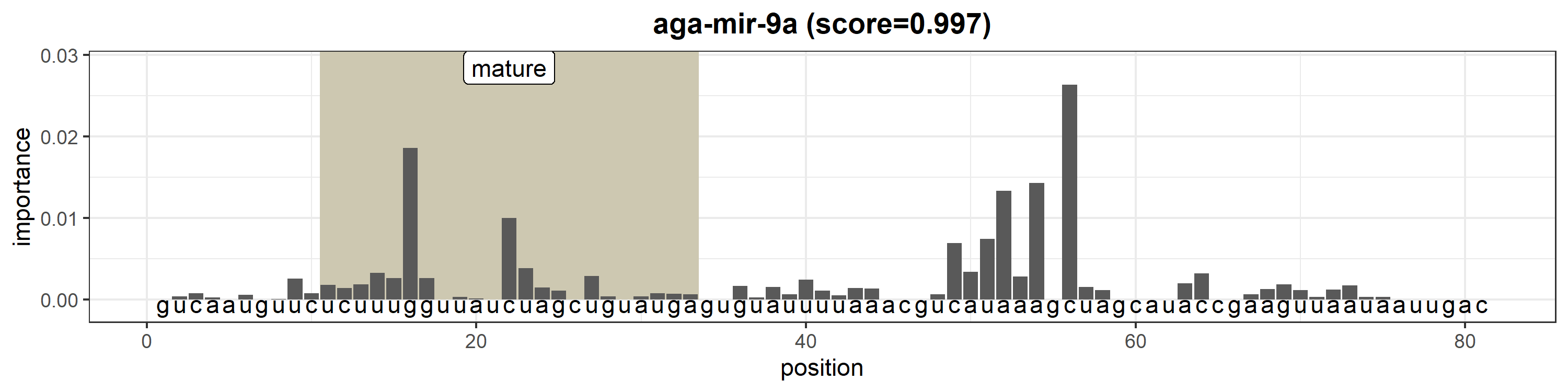

### aga-mir-9b.png

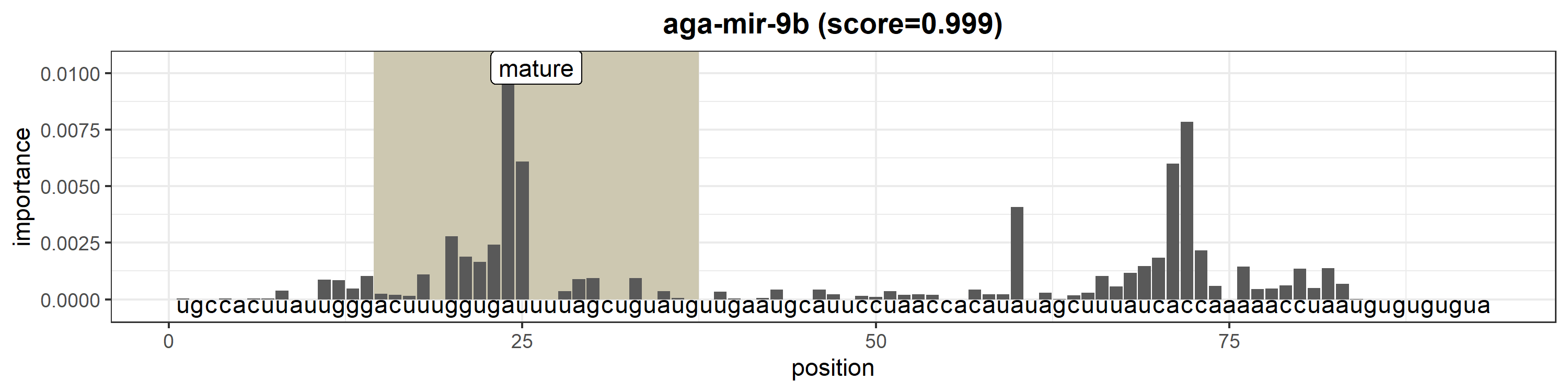

### aga-mir-9c.png

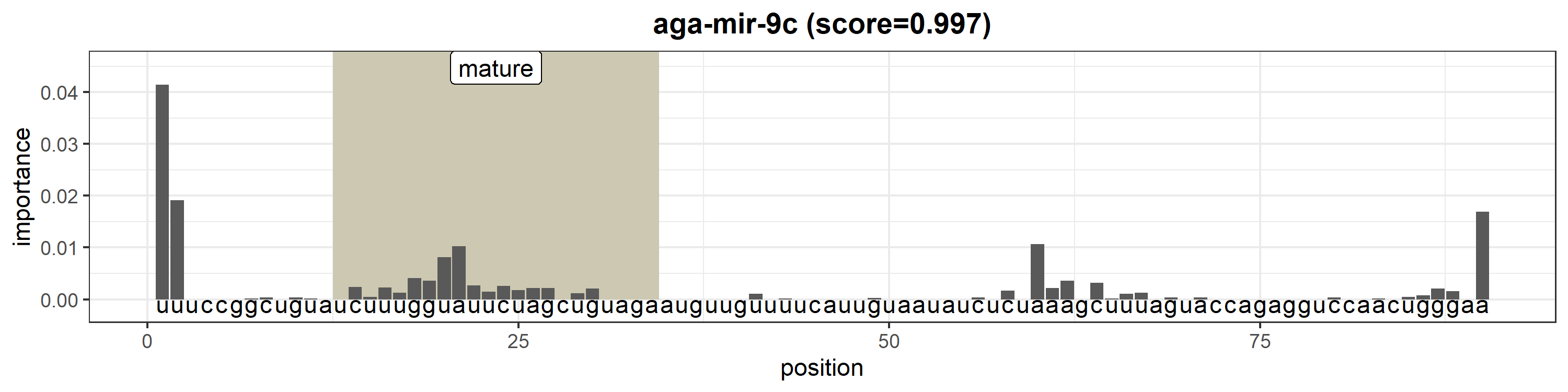

### aga-mir-33.png

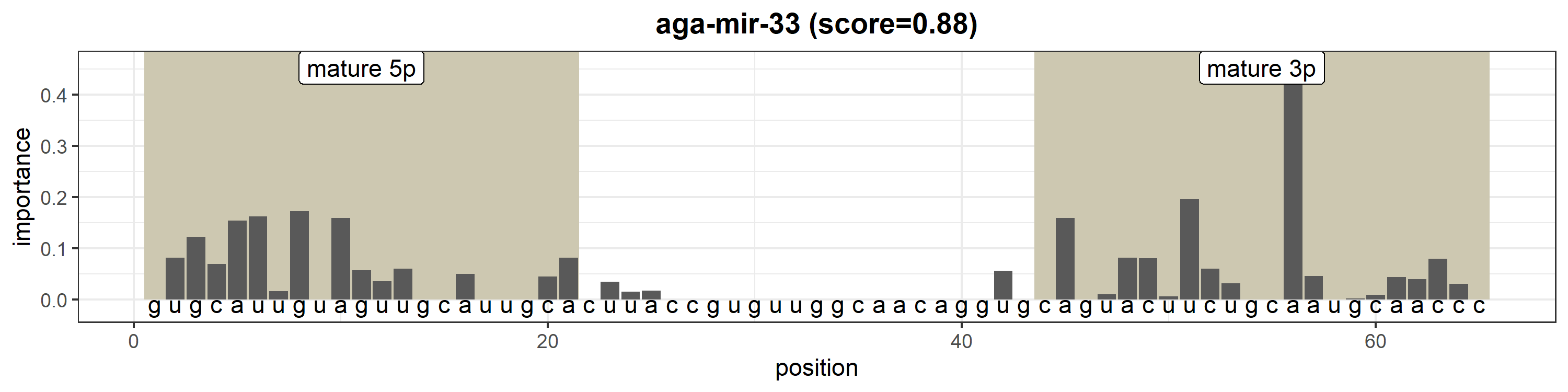

### aga-mir-34.png

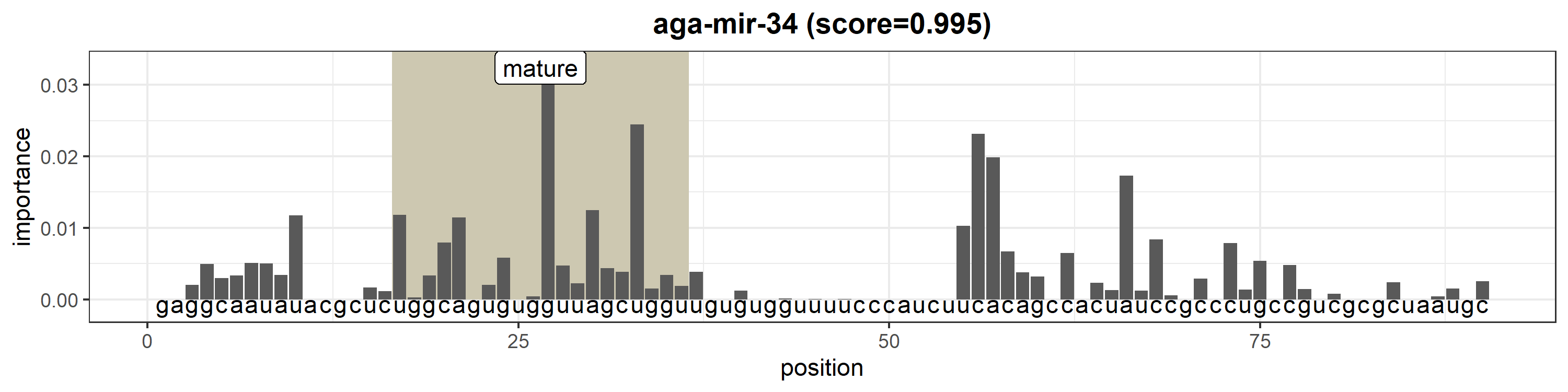

### aga-mir-79.png

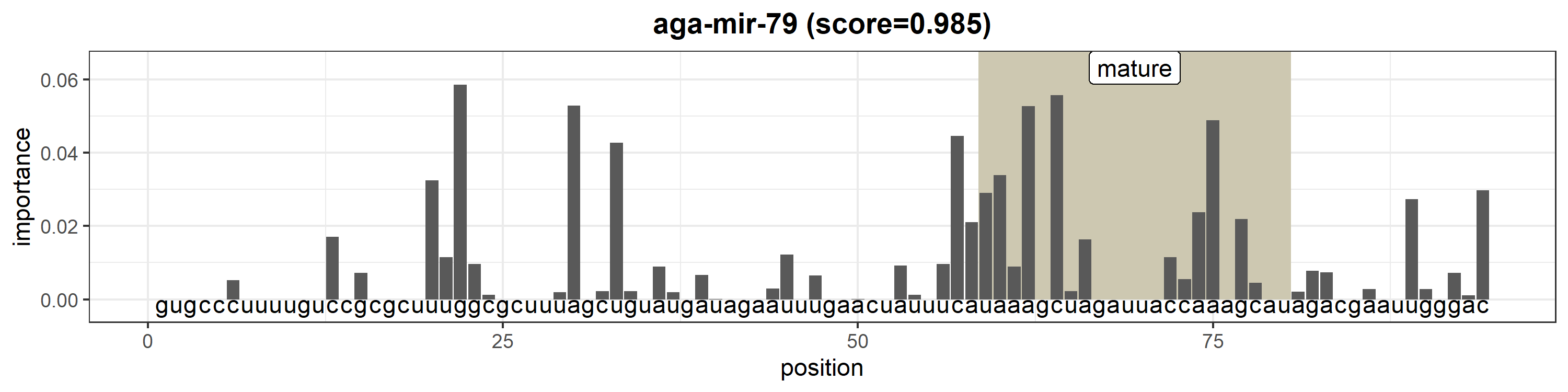

### aga-mir-87.png

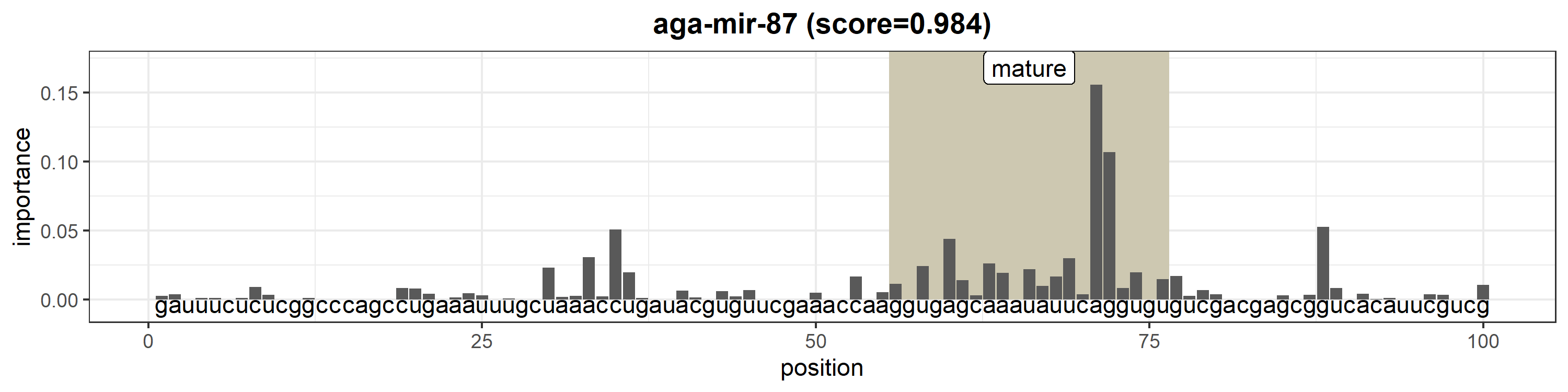

### aga-mir-92a.png

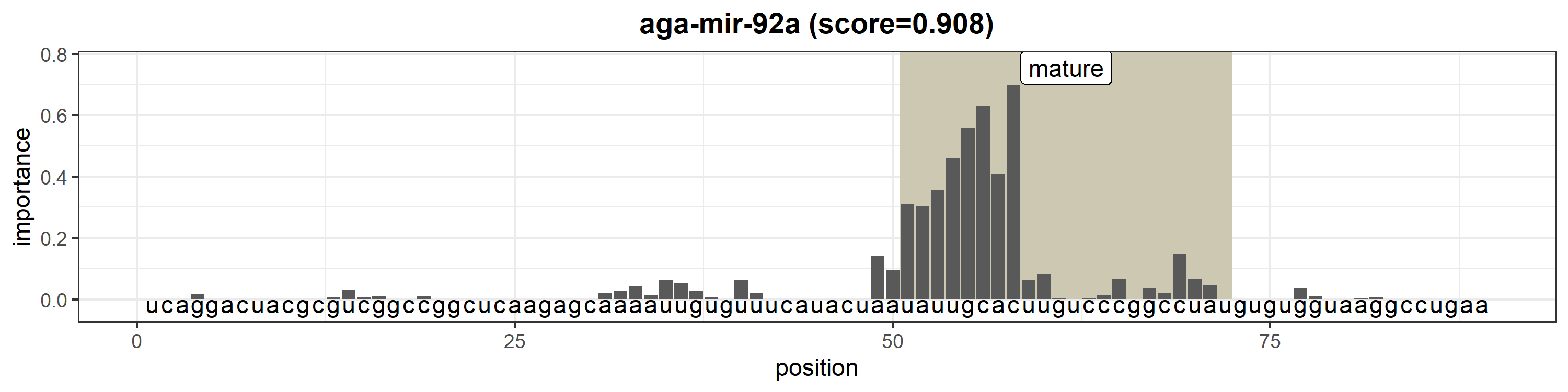

### aga-mir-92b.png

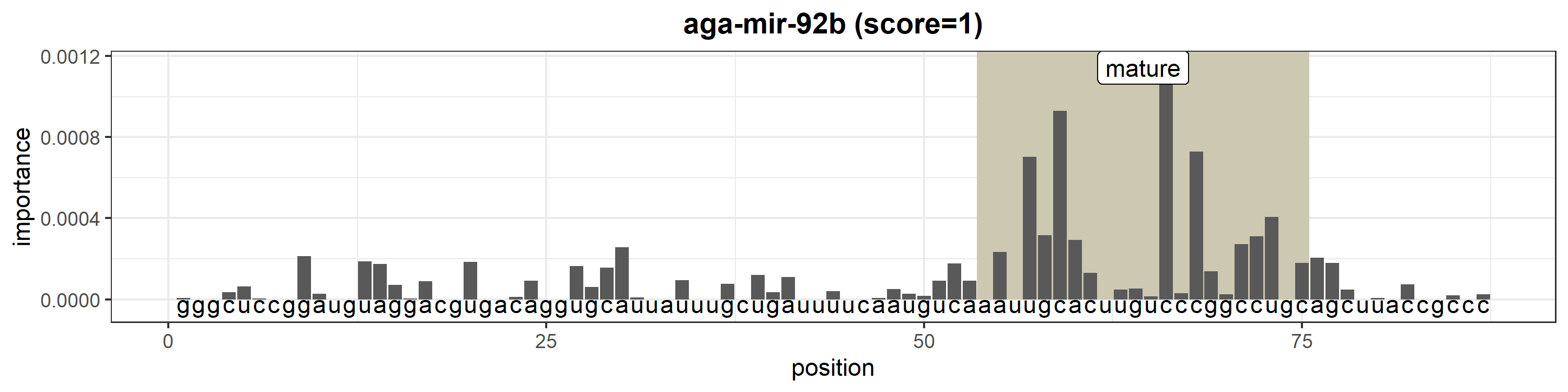

### aga-mir-315.png

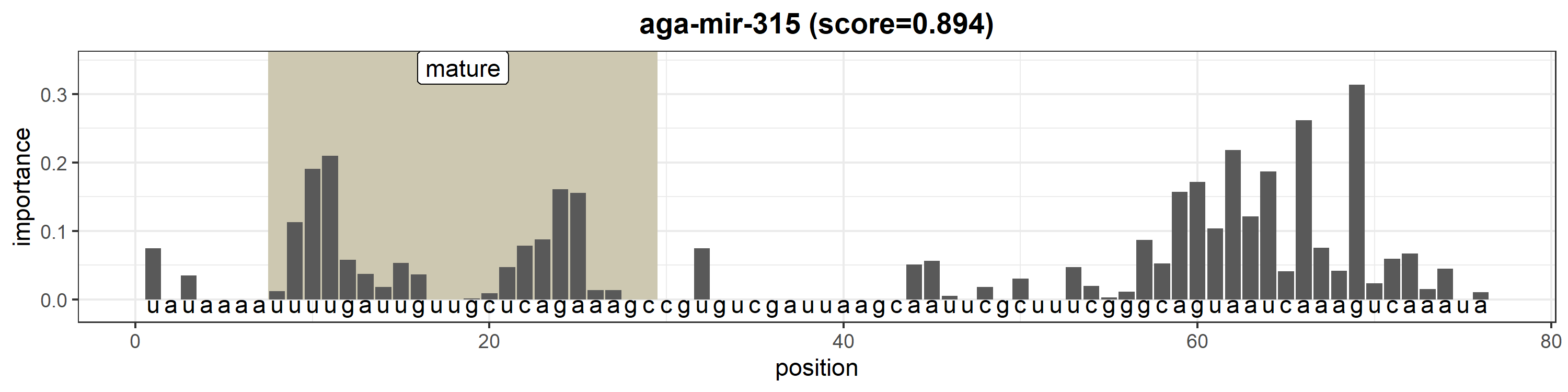

### aga-mir-317.png

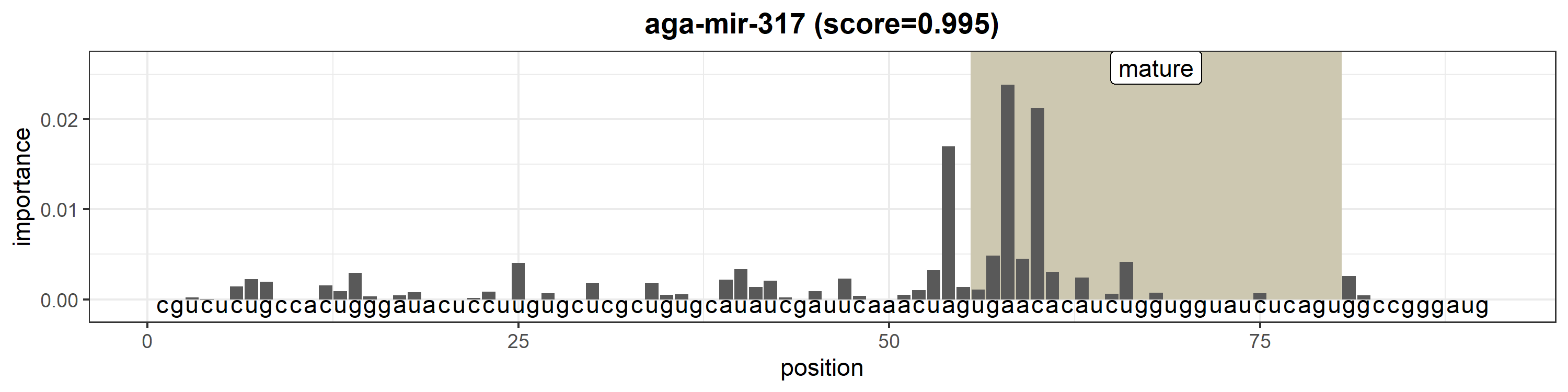

### aga-mir-375.png

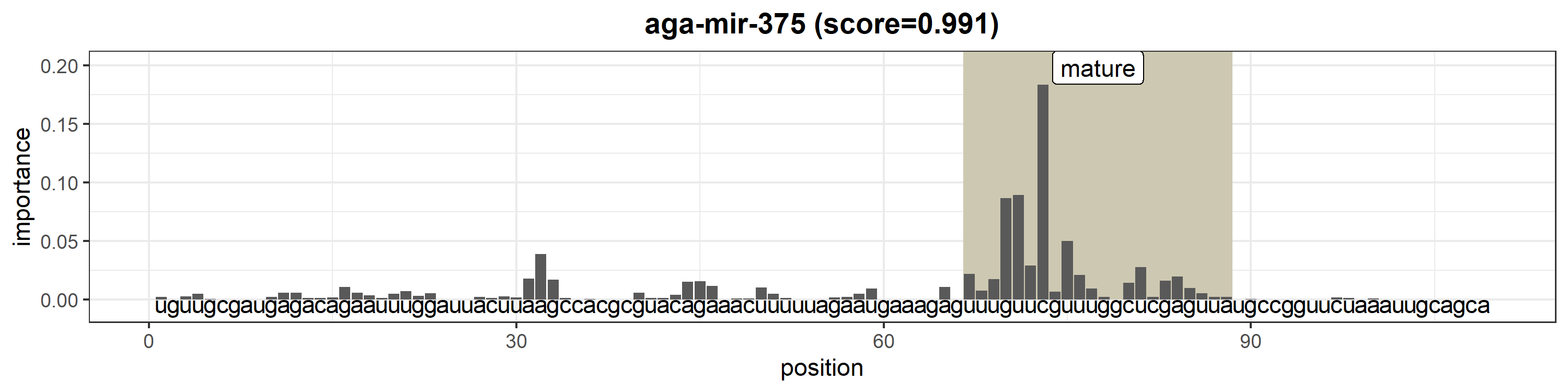

### aga-mir-927.png

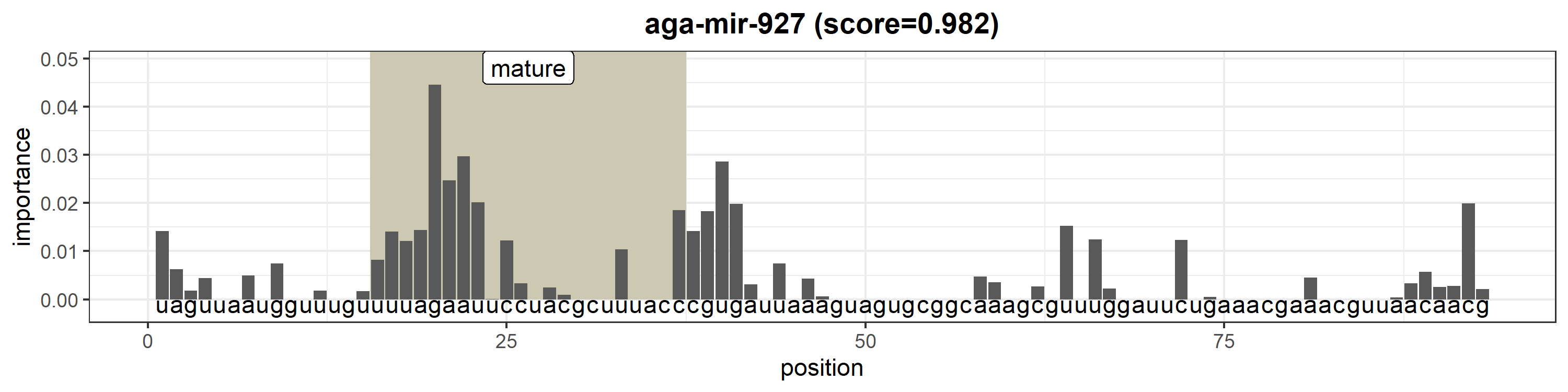

### aga-mir-929.png

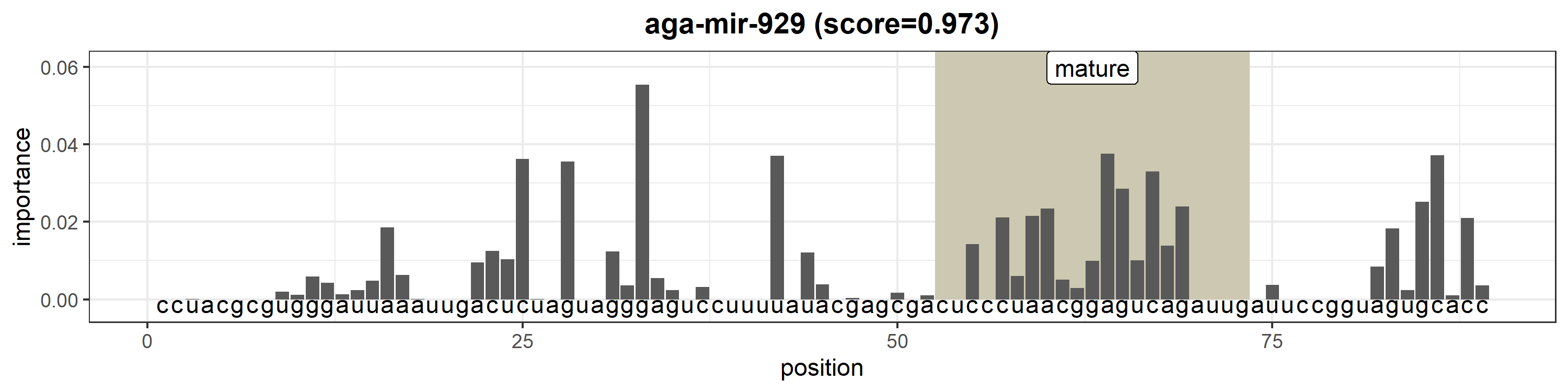

### aga-mir-932.png

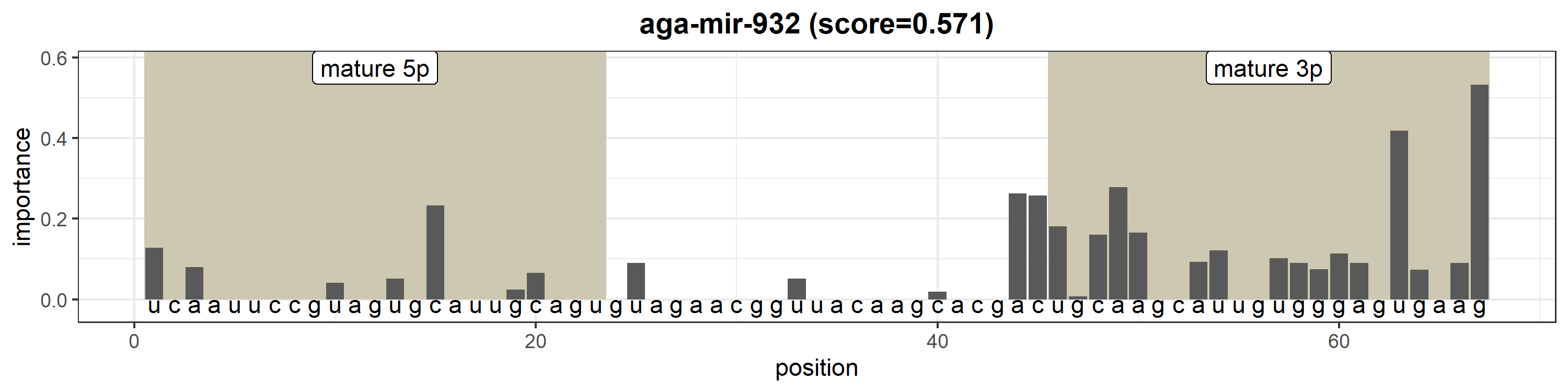

### aga-mir-957.png

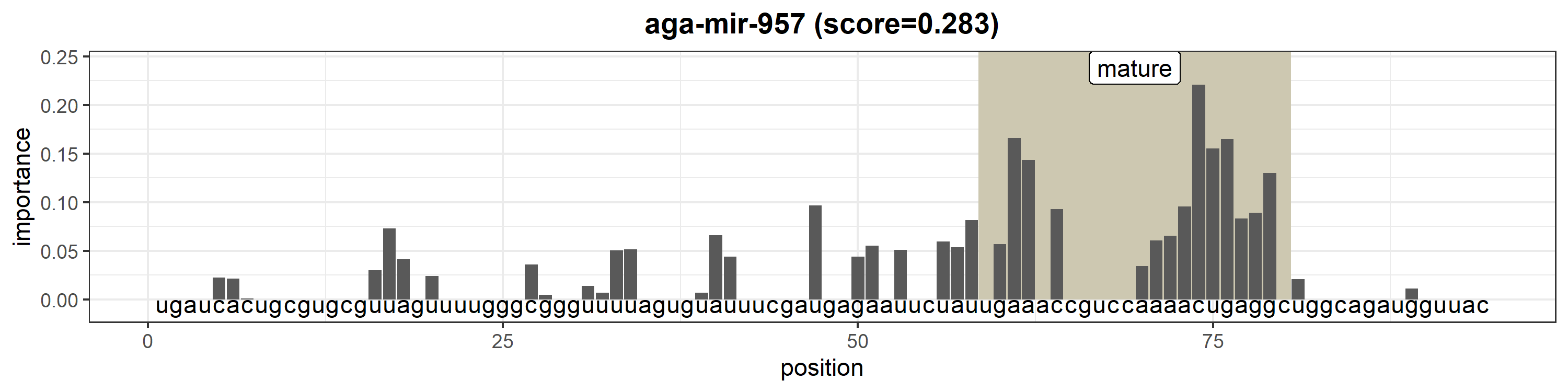

### aga-mir-965-1.png

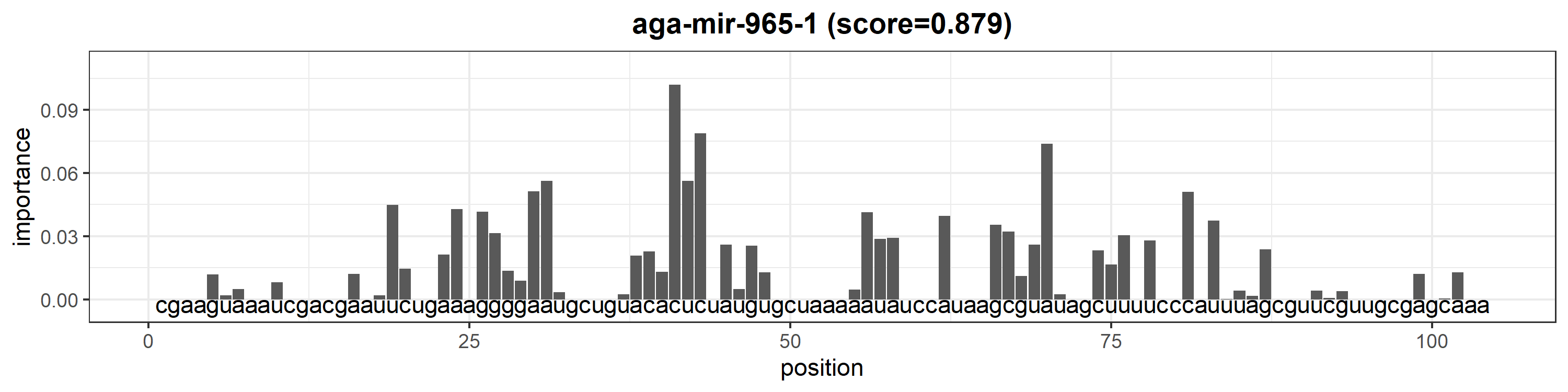

### aga-mir-965-2.png

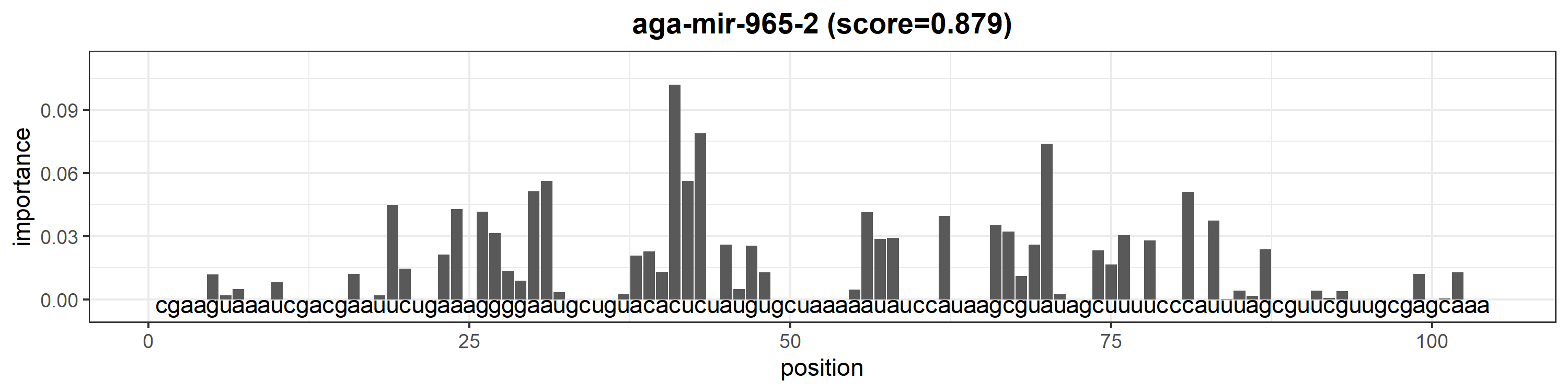

### aga-mir-970.png

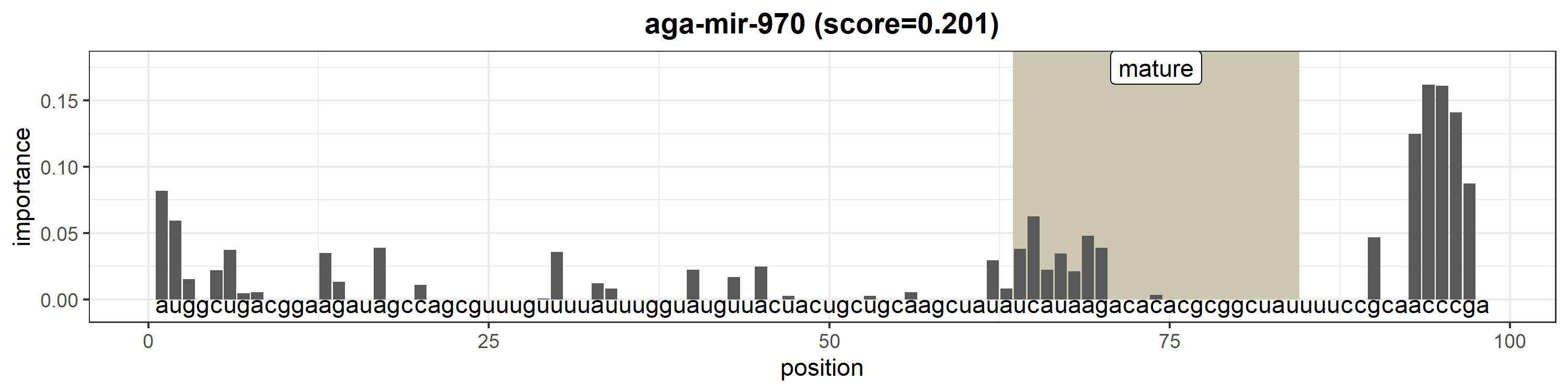

### aga-mir-980.png

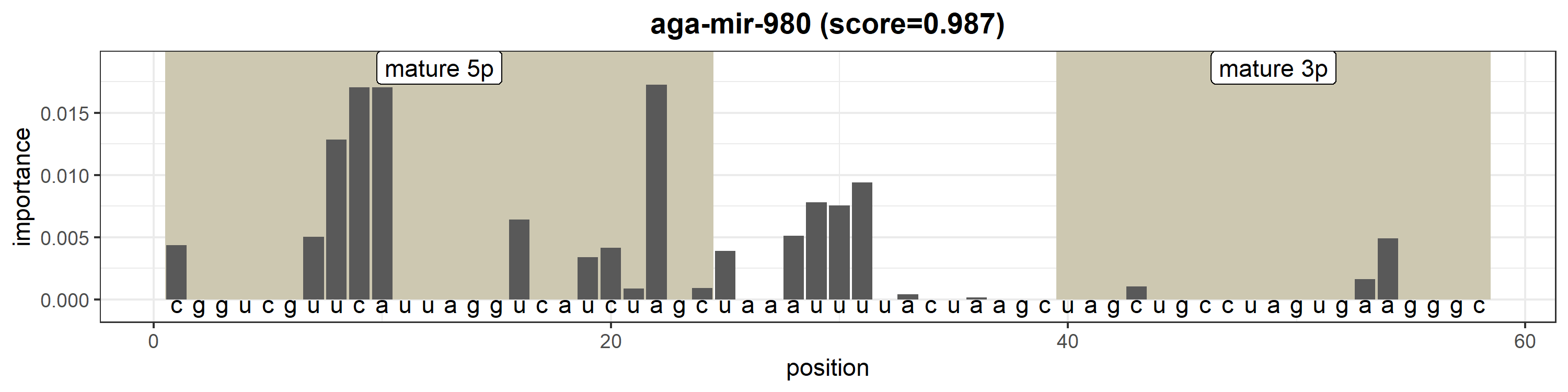

### aga-mir-981.png

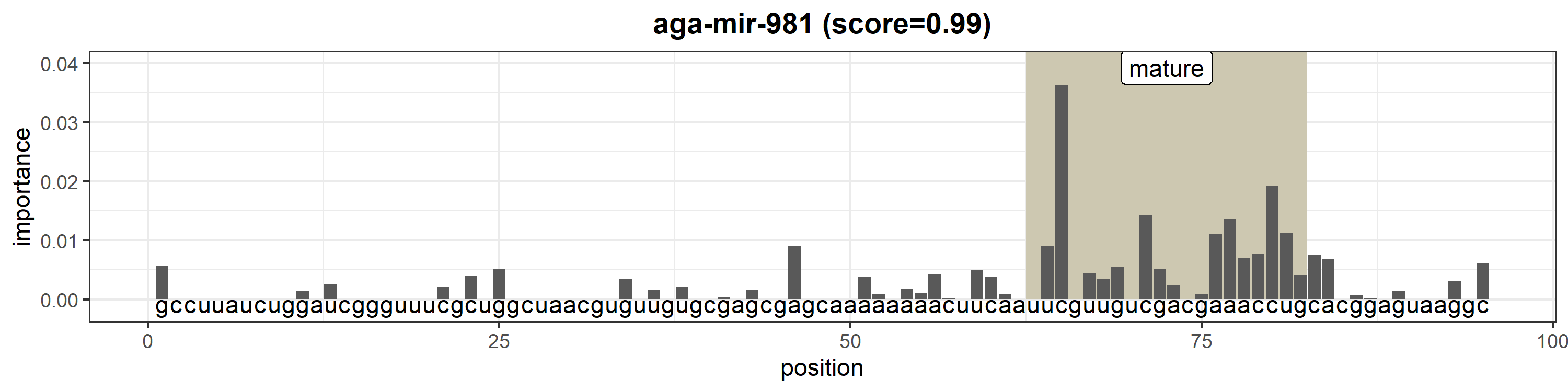

### aga-mir-988.png

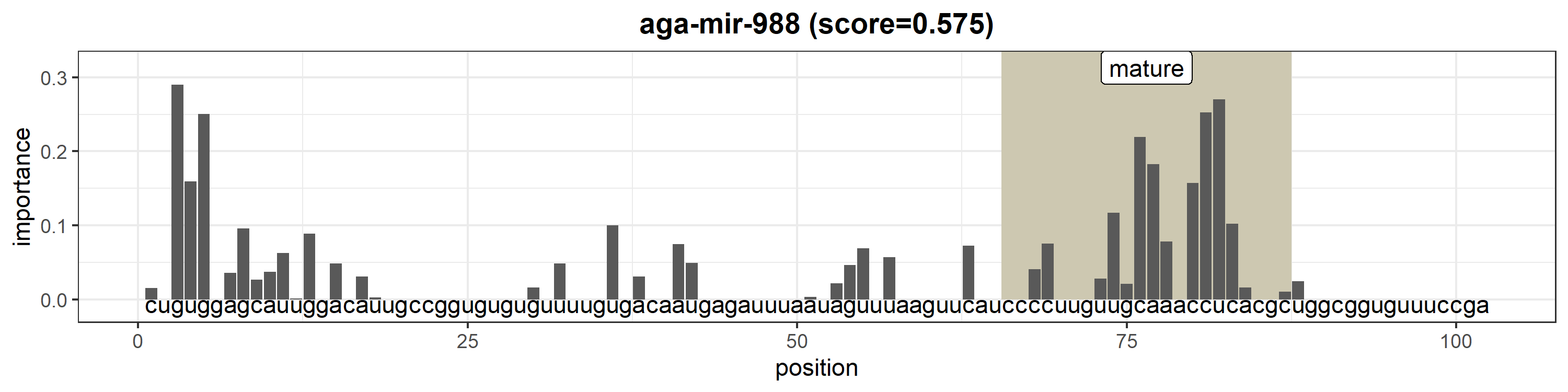

### aga-mir-989.png

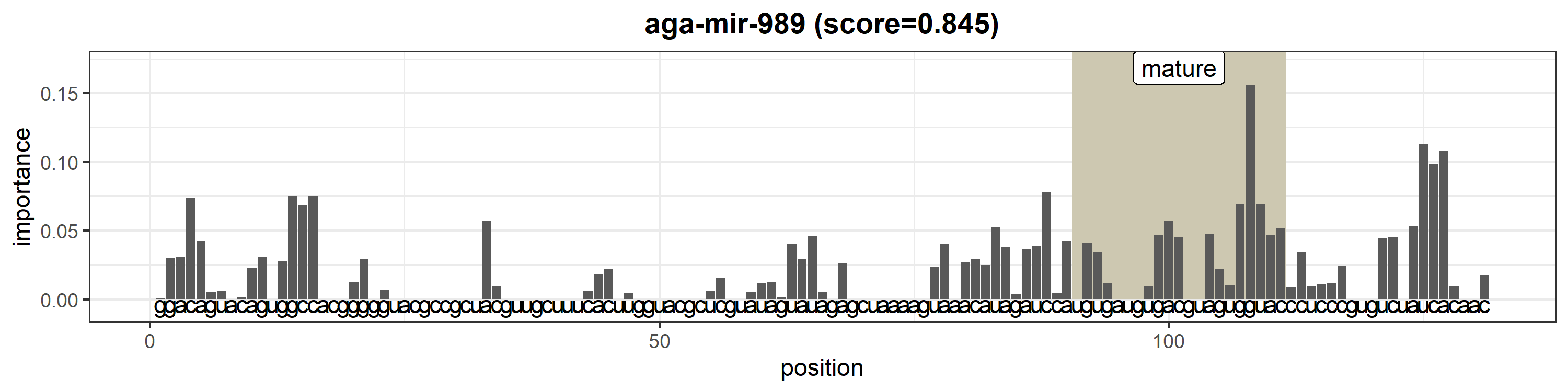

### aga-mir-993.png

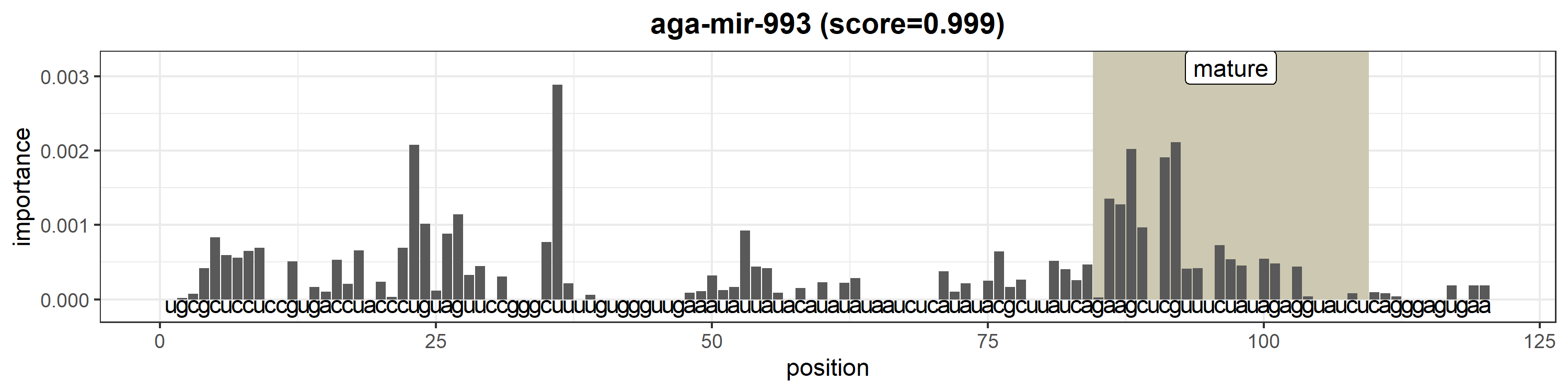

### aga-mir-996.png

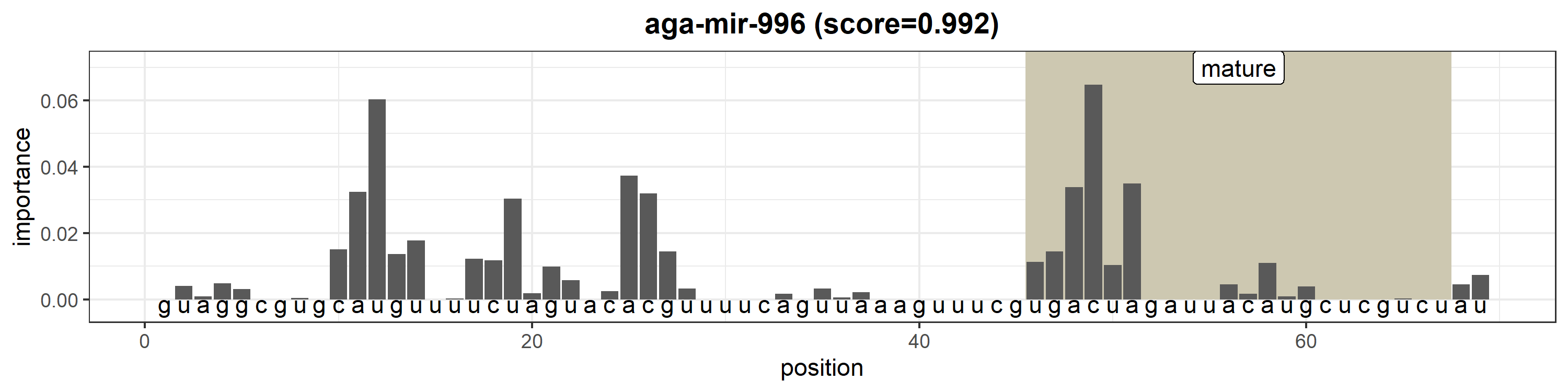

### aga-mir-998.png

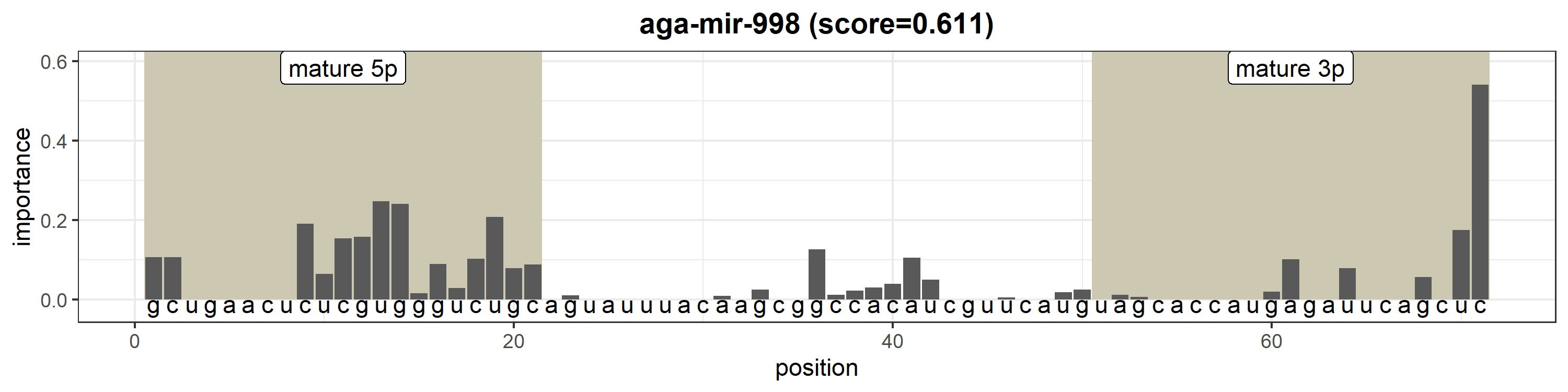

### aga-mir-iab-4.png

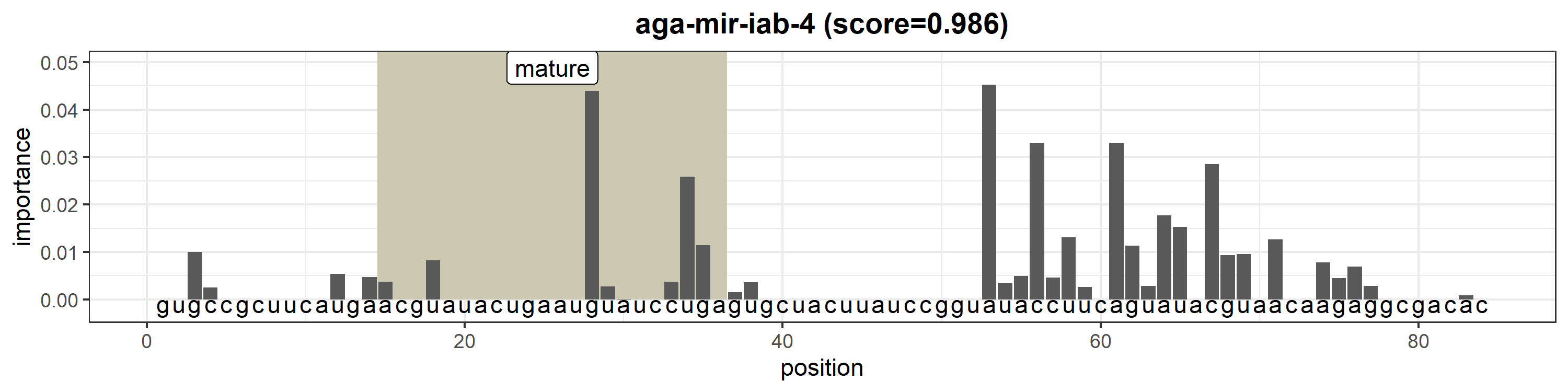

### ath-MIR156a.png

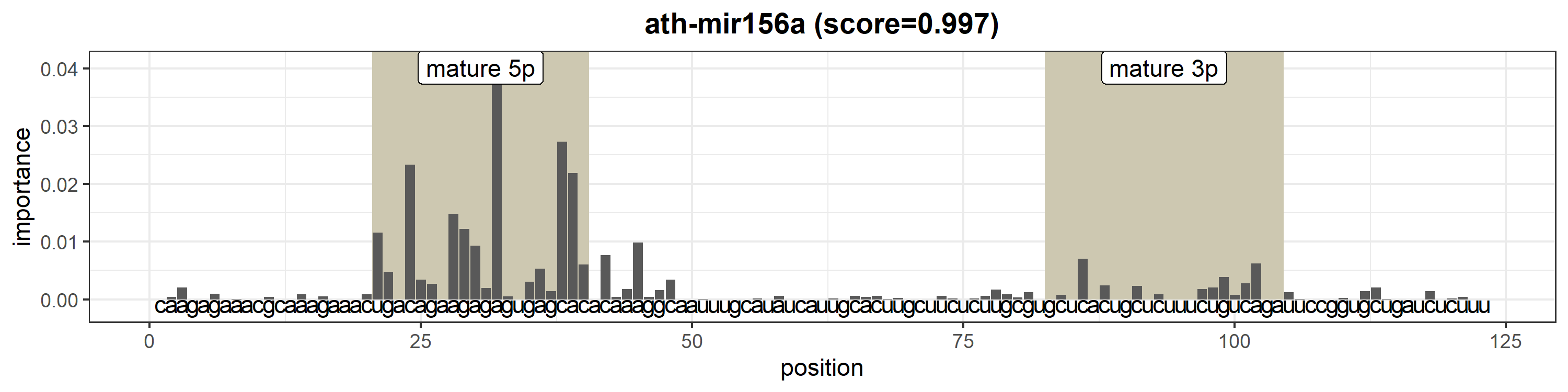

### ath-MIR156b.png

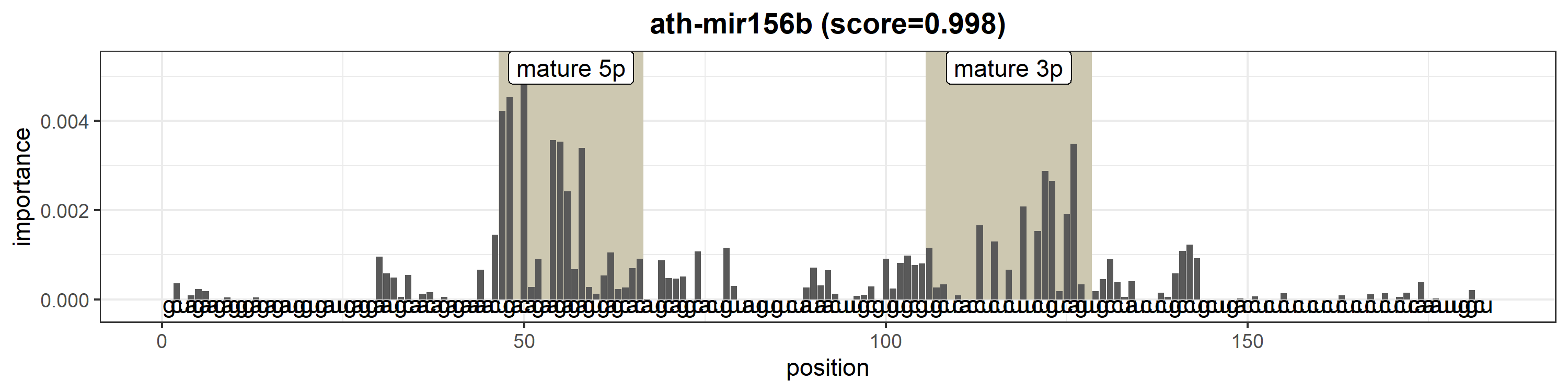

### ath-MIR156c.png

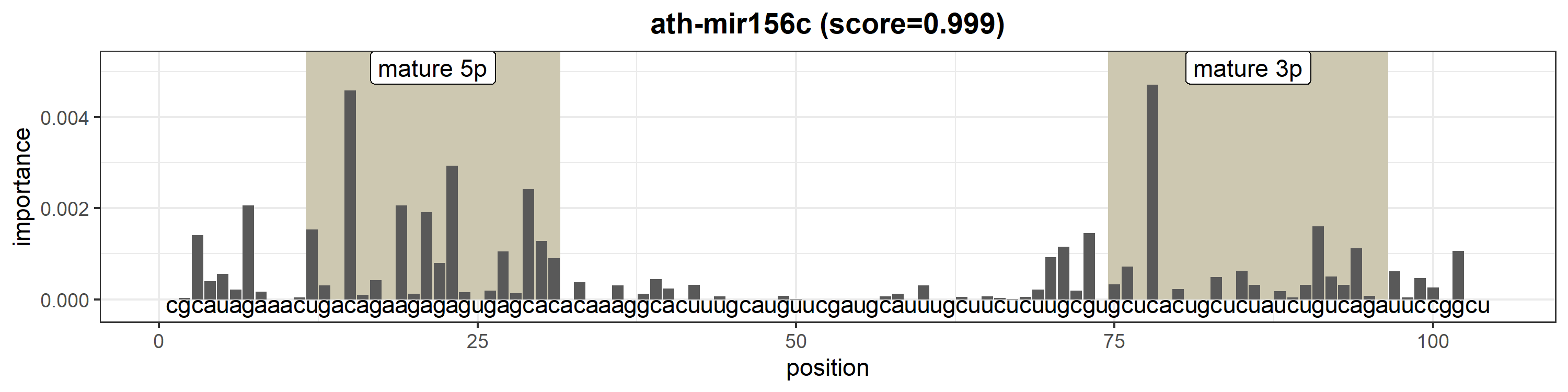

### ath-MIR156d.png

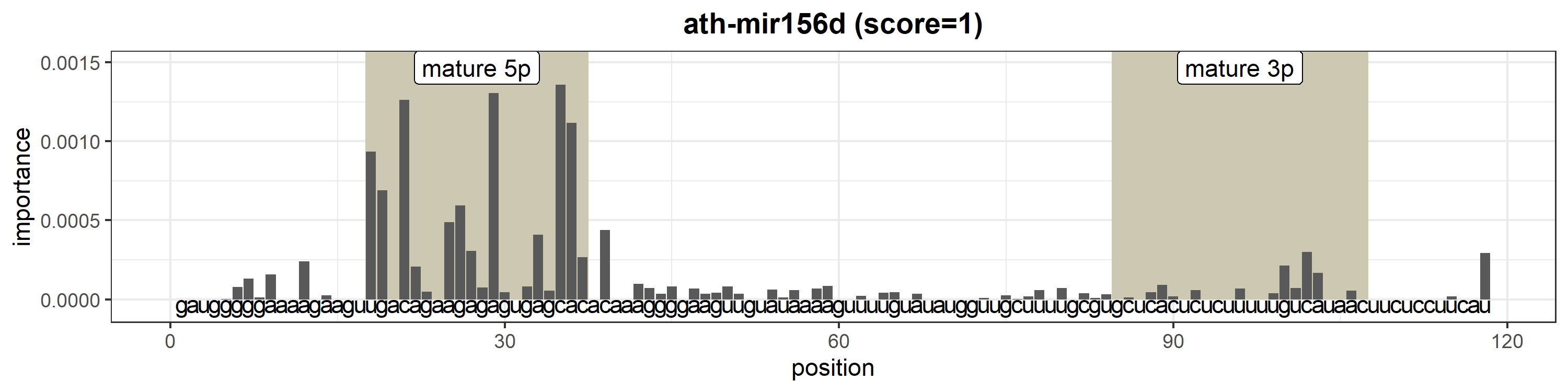

### ath-MIR156e.png

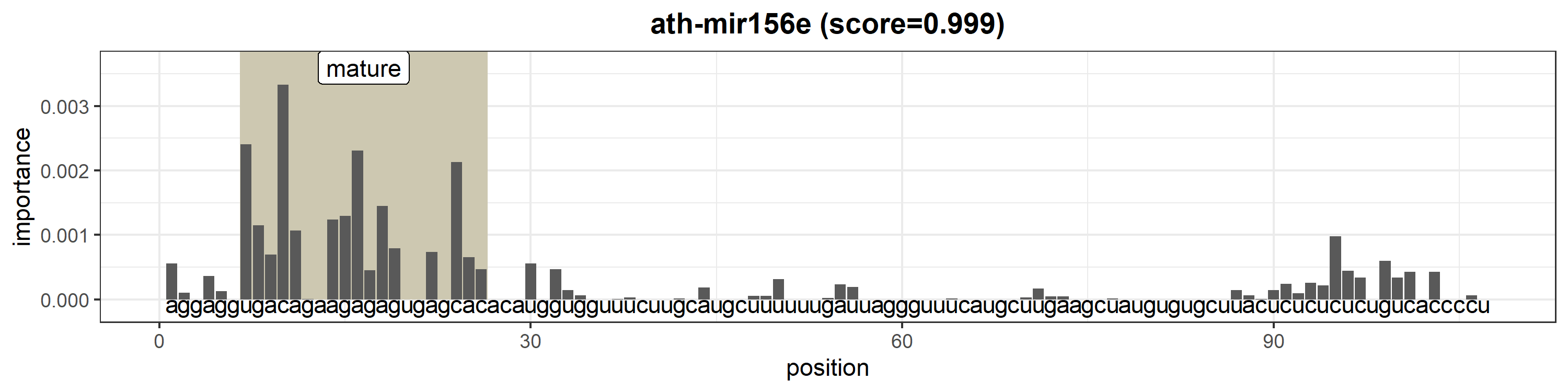

### ath-MIR156f.png

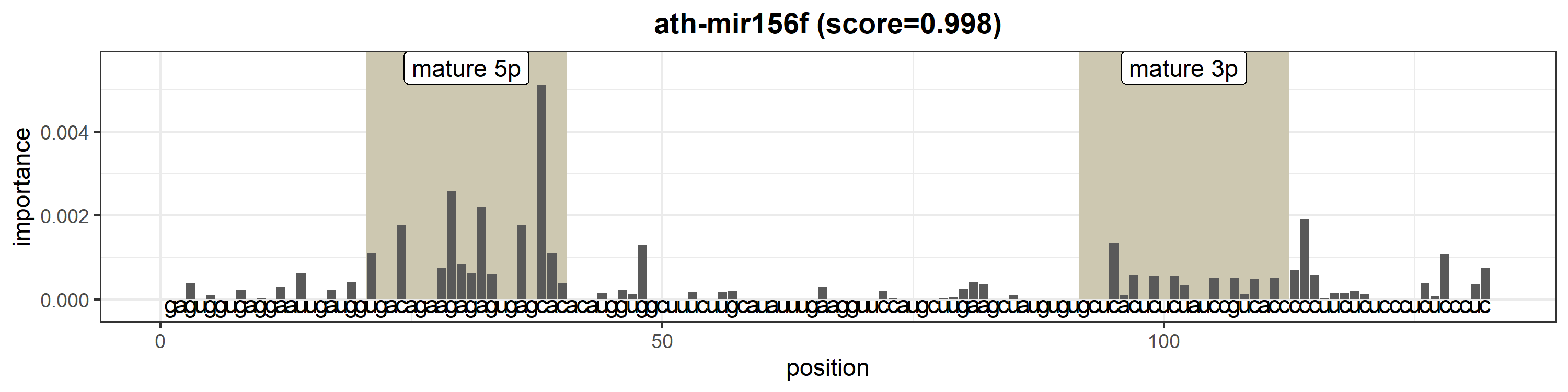

### ath-MIR156g.png

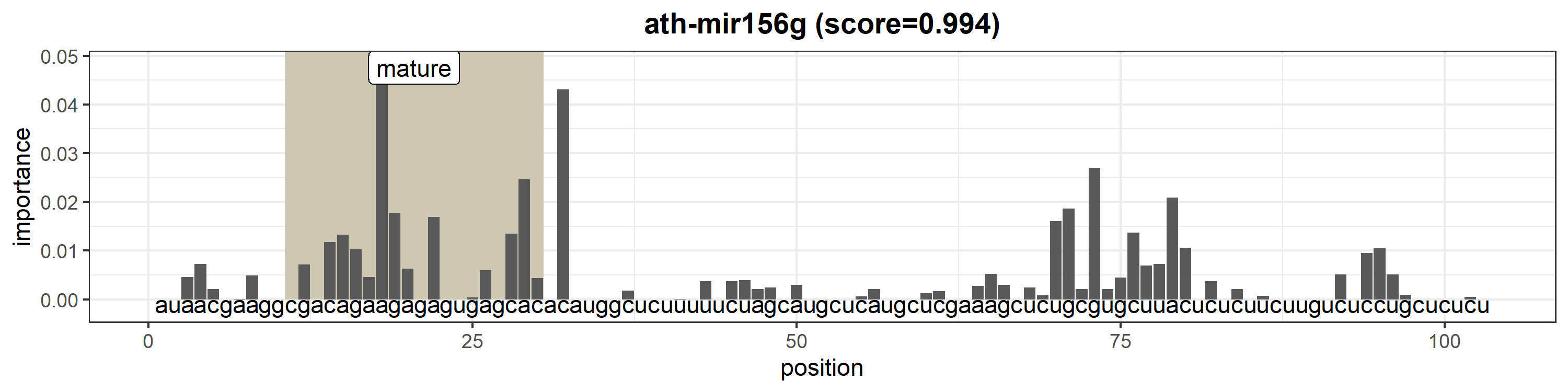

### ath-MIR156h.png

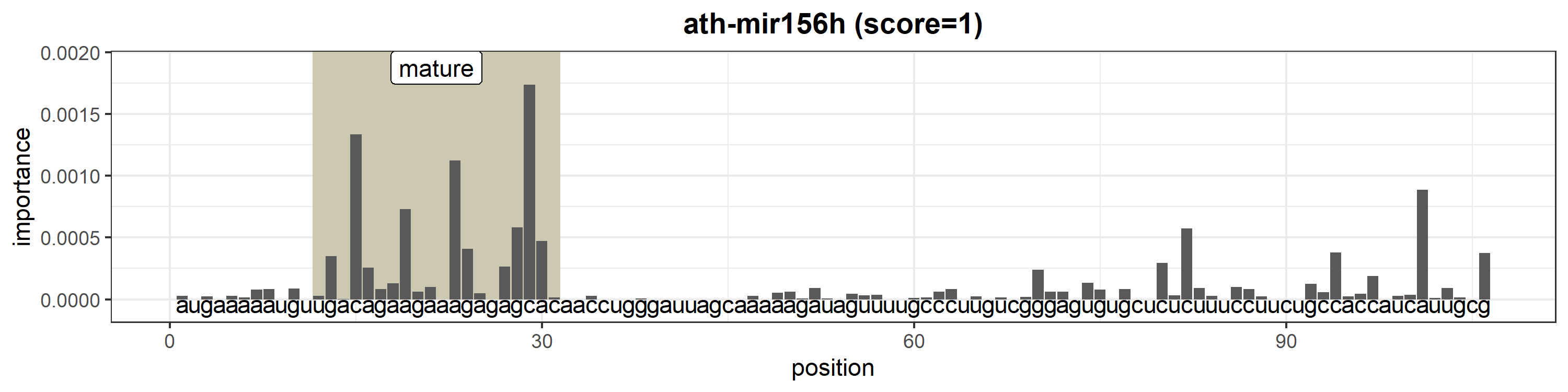

### ath-MIR156i.png

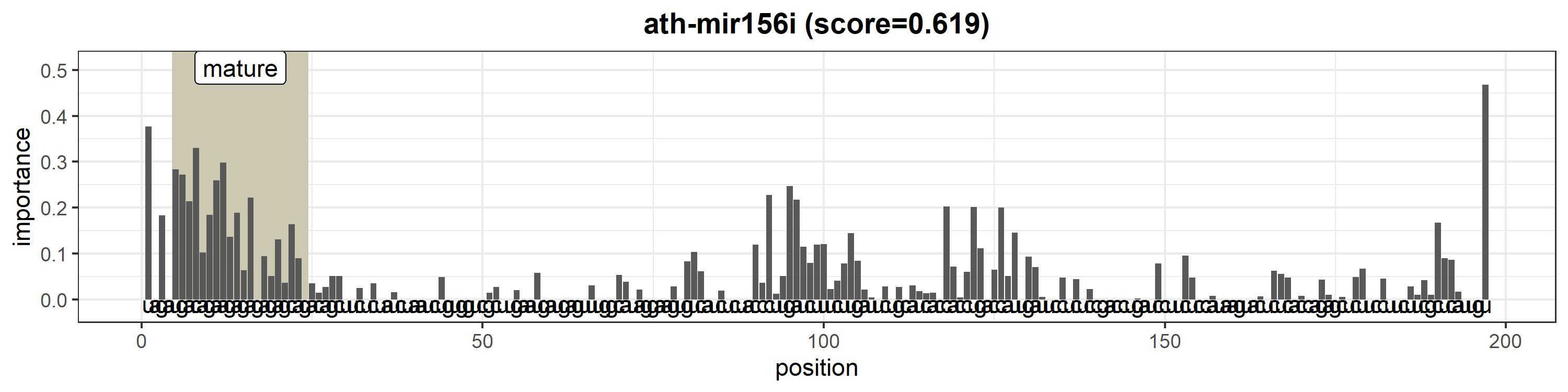

### ath-MIR156j.png

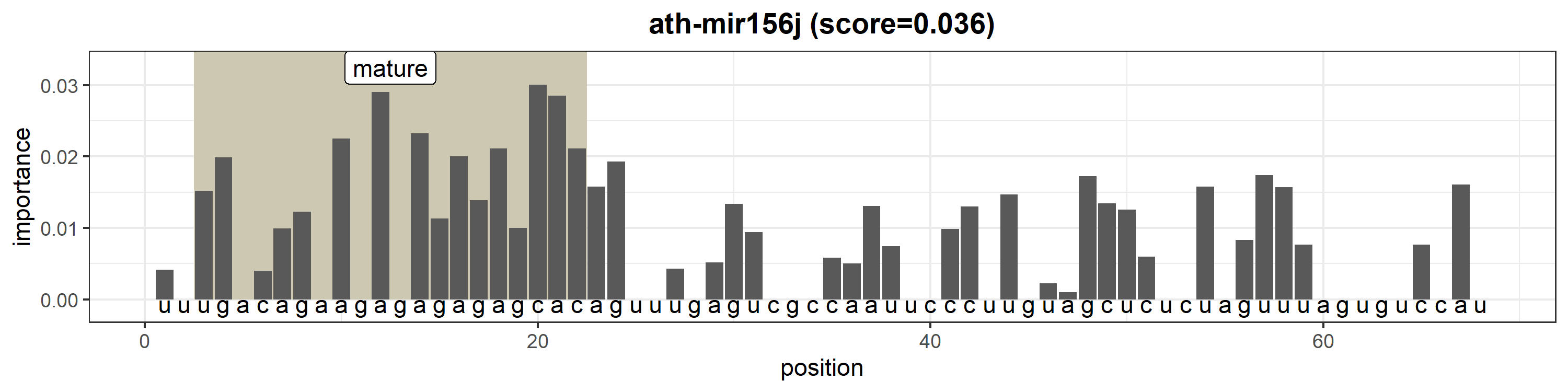

### ath-MIR157a.png

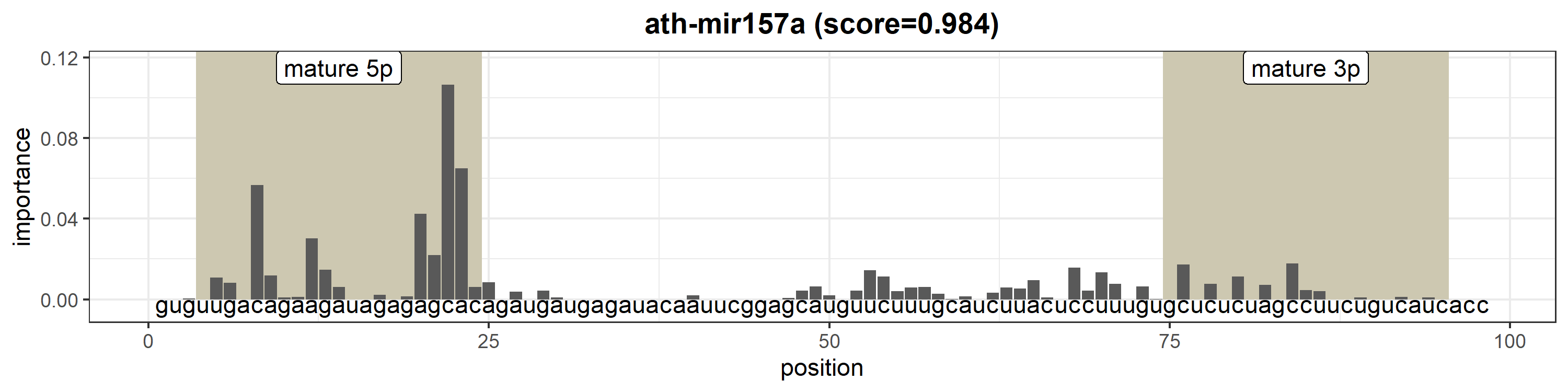

### ath-MIR157b.png

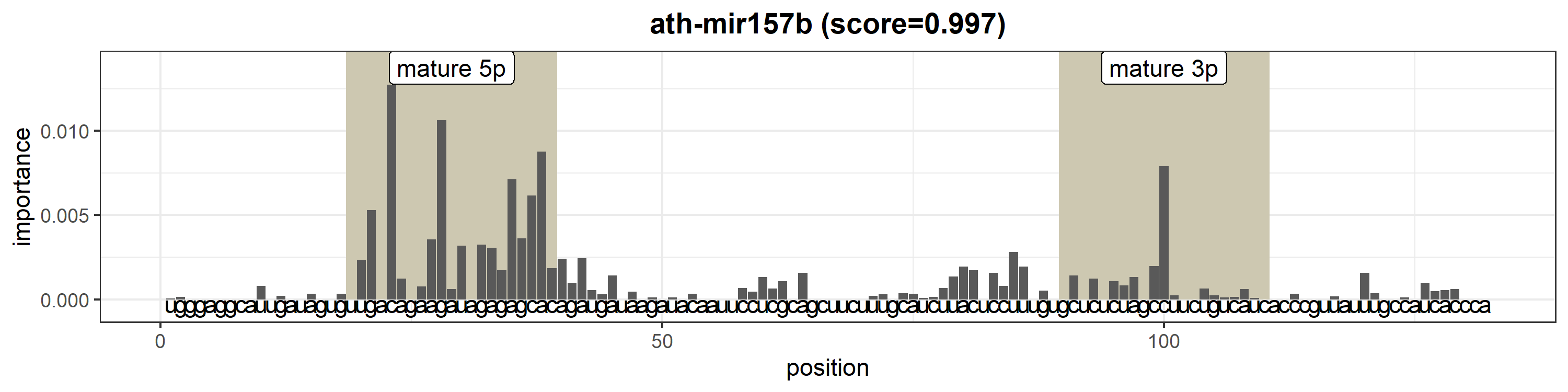

### ath-MIR157d.png

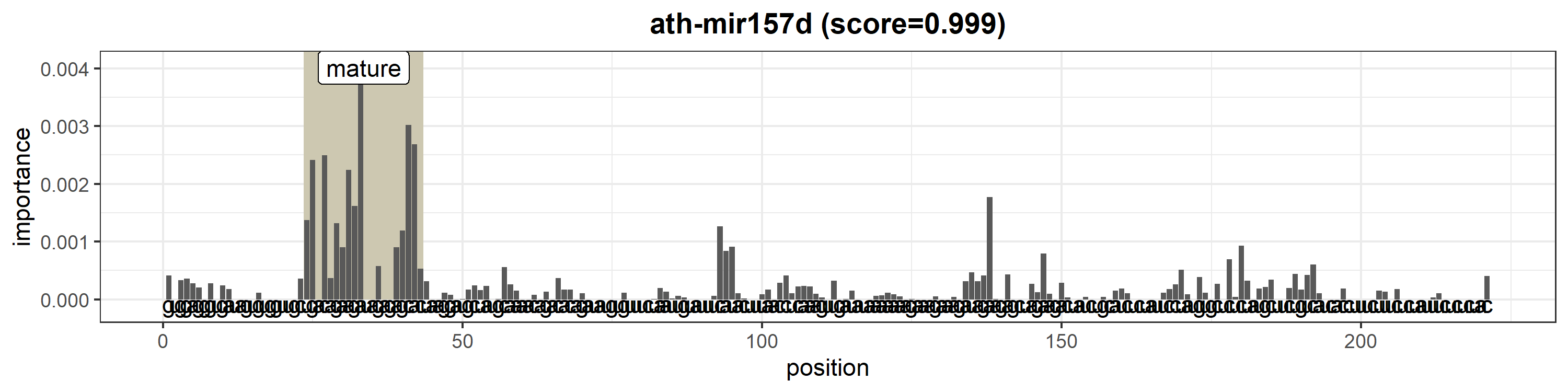

### ath-MIR158a.png

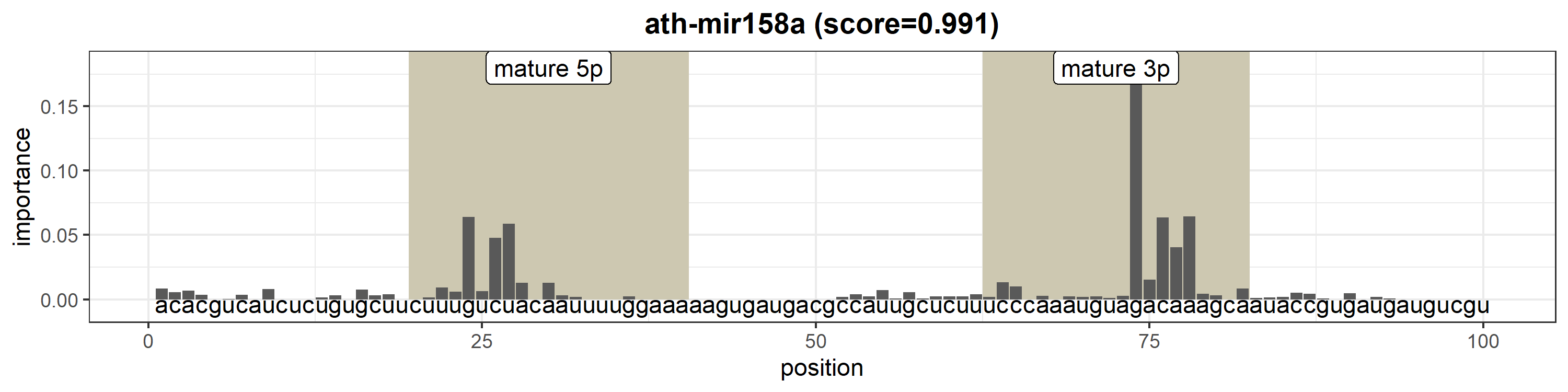

### ath-MIR158b.png

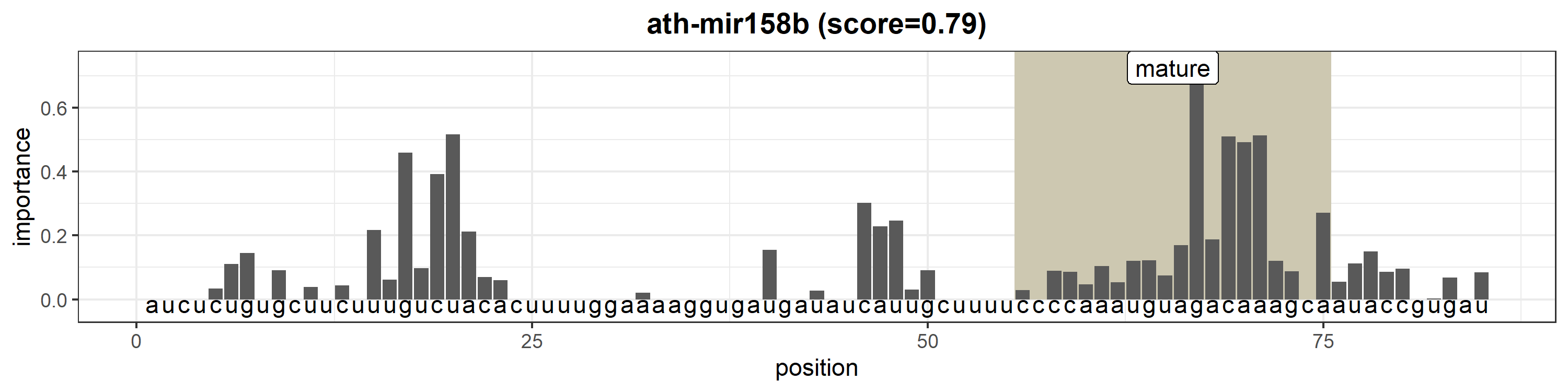

### ath-MIR159a.png

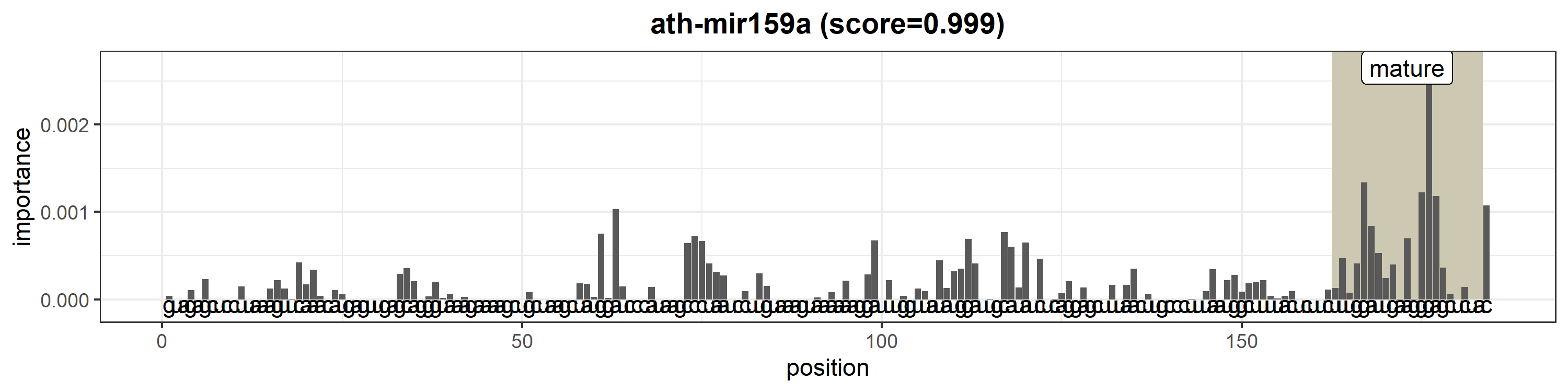

### ath-MIR159b.png

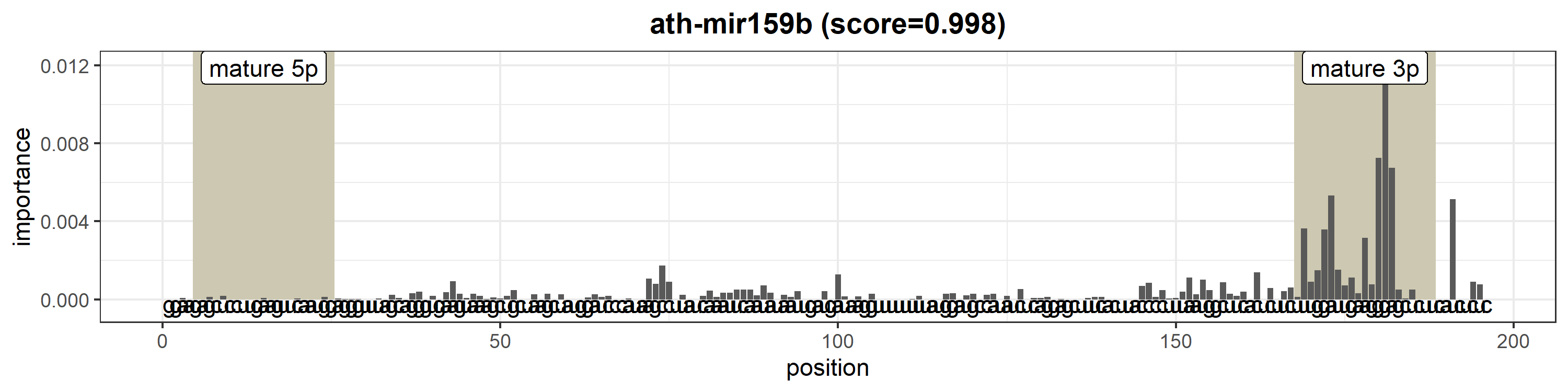

### ath-MIR159c.png

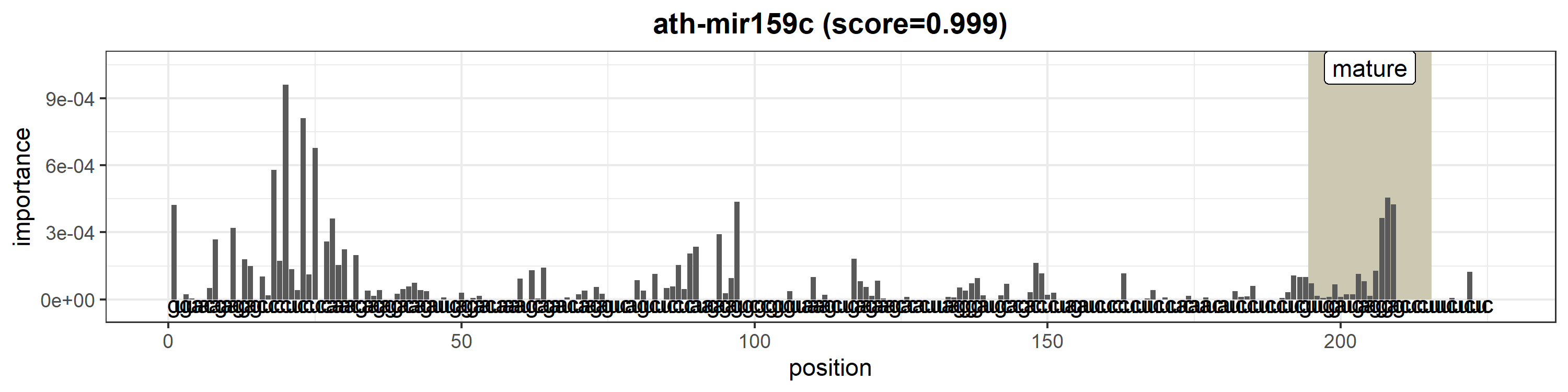

### ath-MIR160a.png

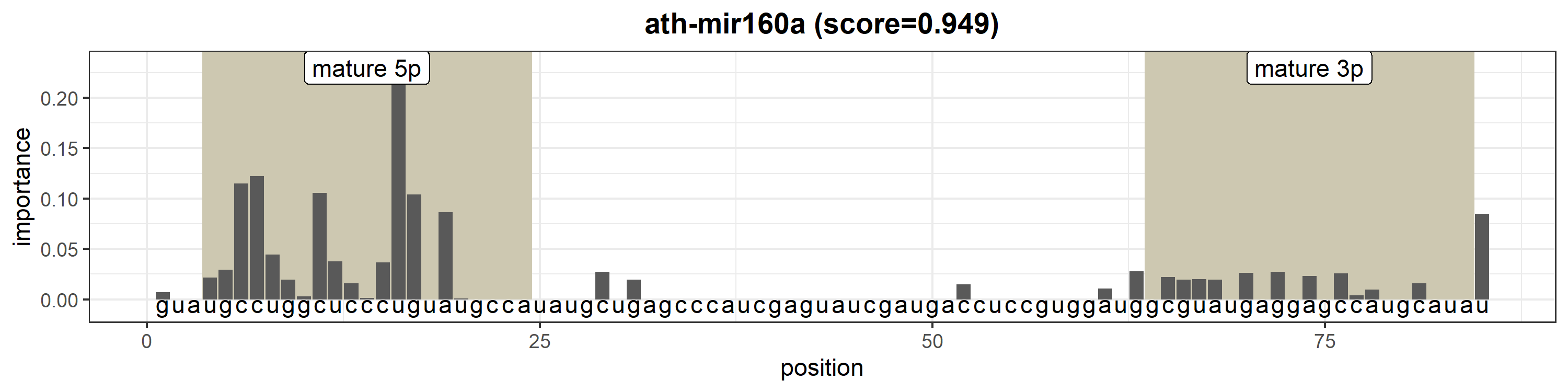

### ath-MIR160b.png

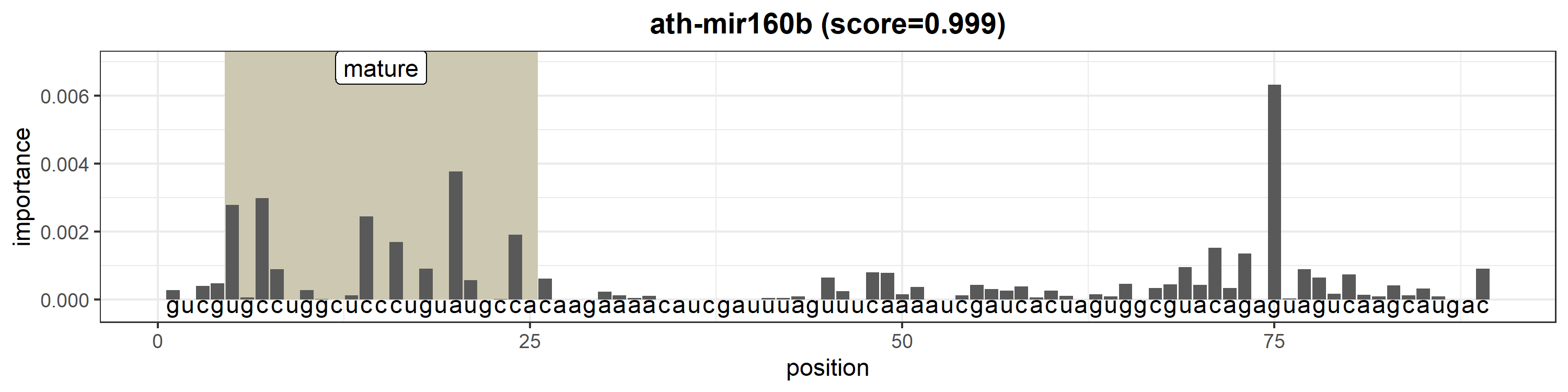

### ath-MIR160c.png

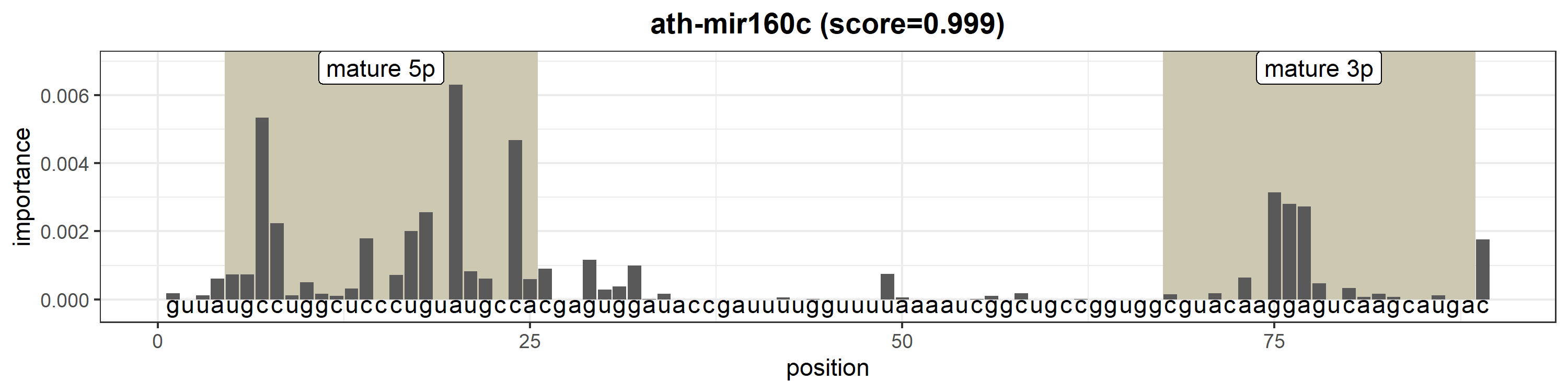

### ath-MIR161.png

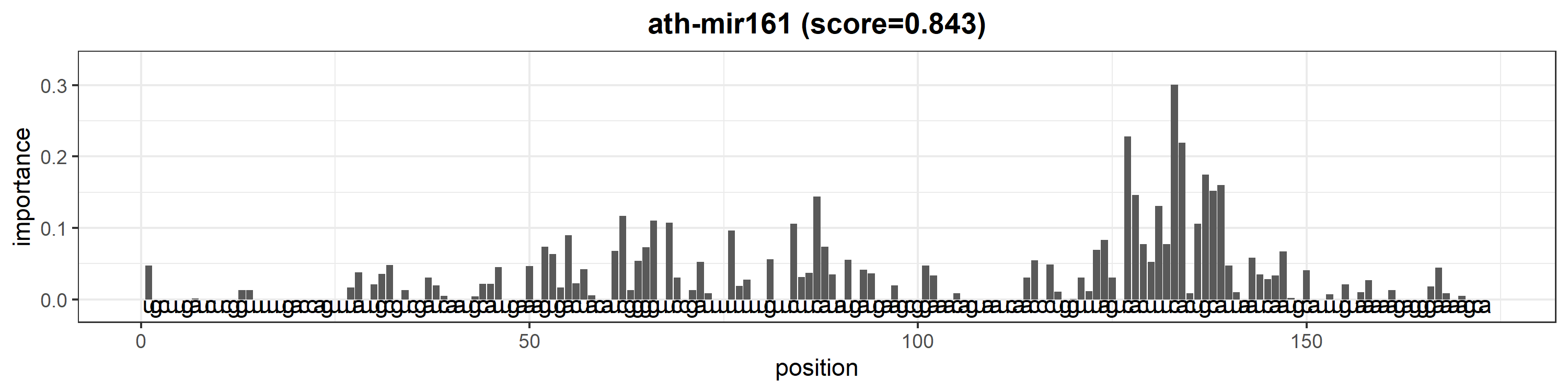

### ath-MIR162a.png

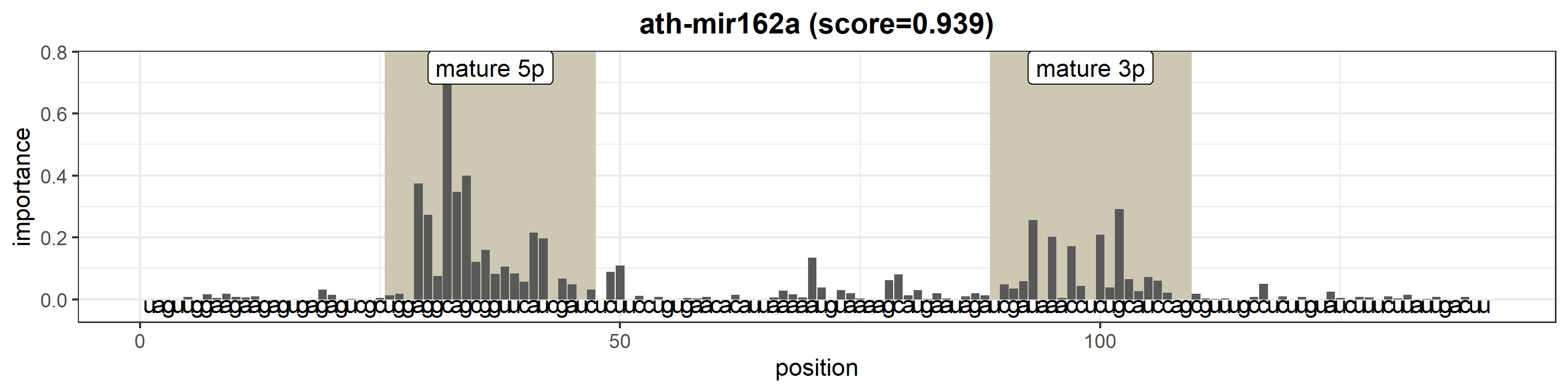

### ath-MIR162b.png

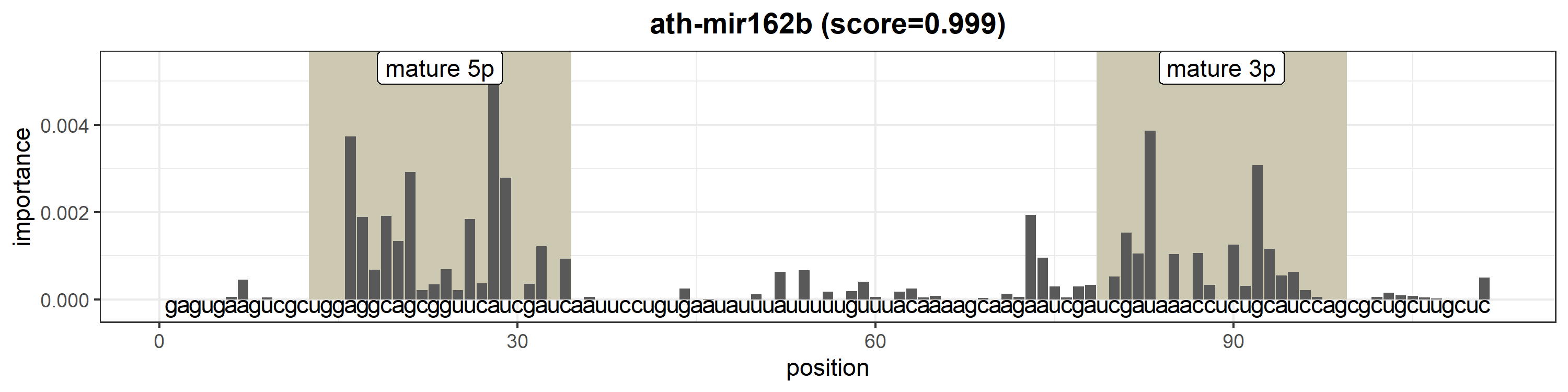

### ath-MIR164a.png

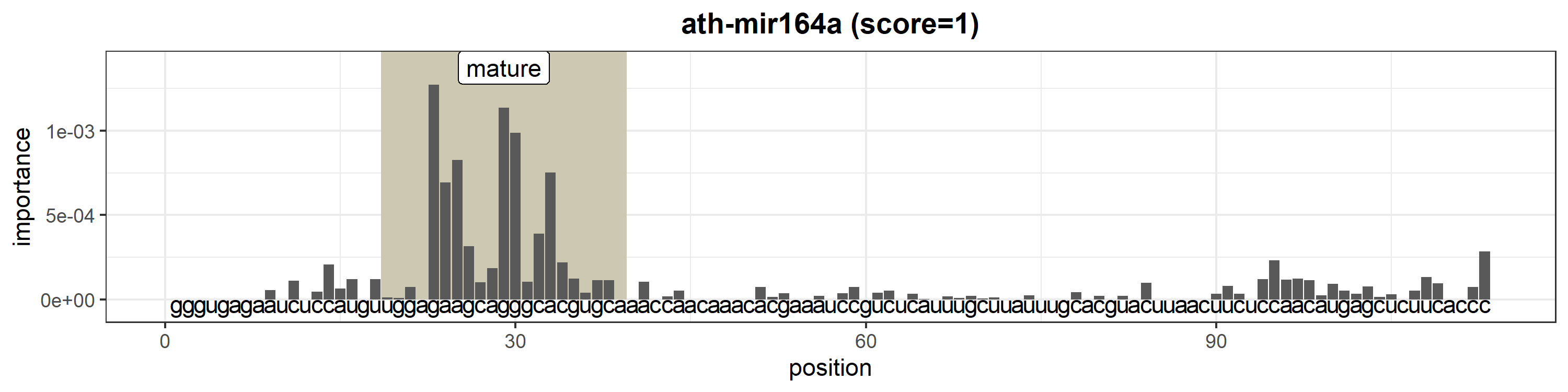

### ath-MIR164b.png

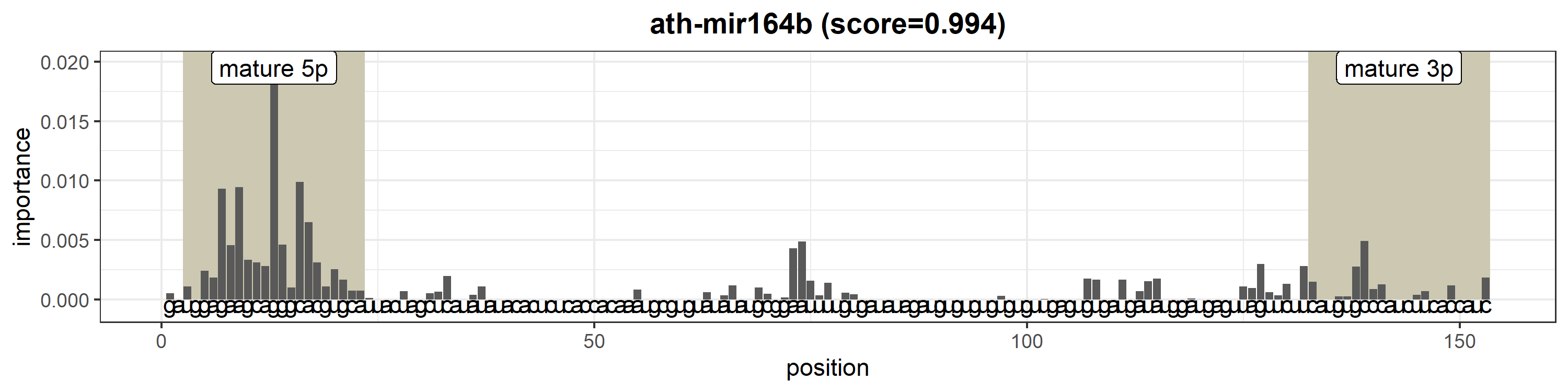

### ath-MIR164c.png

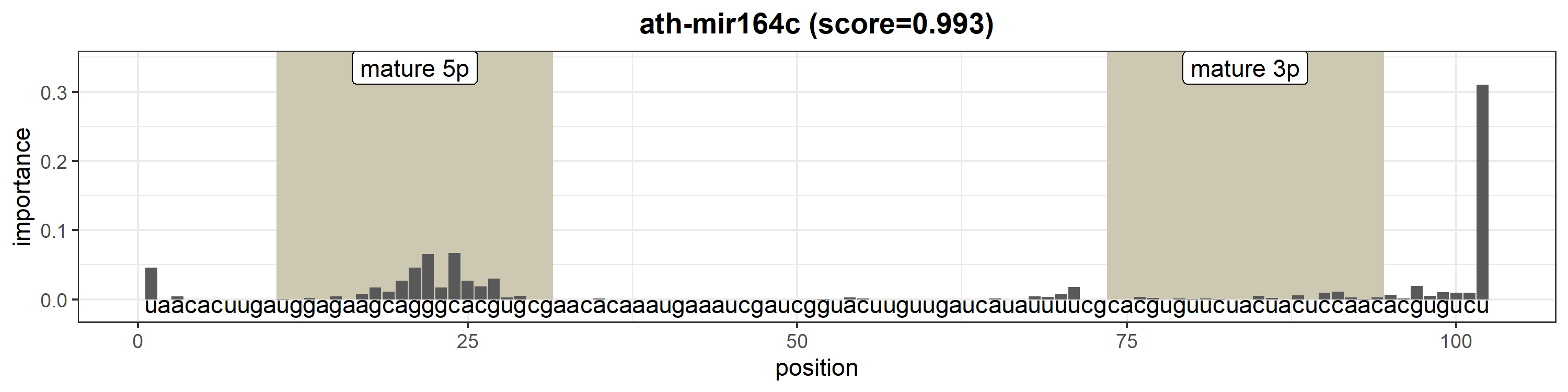

### ath-MIR165a.png

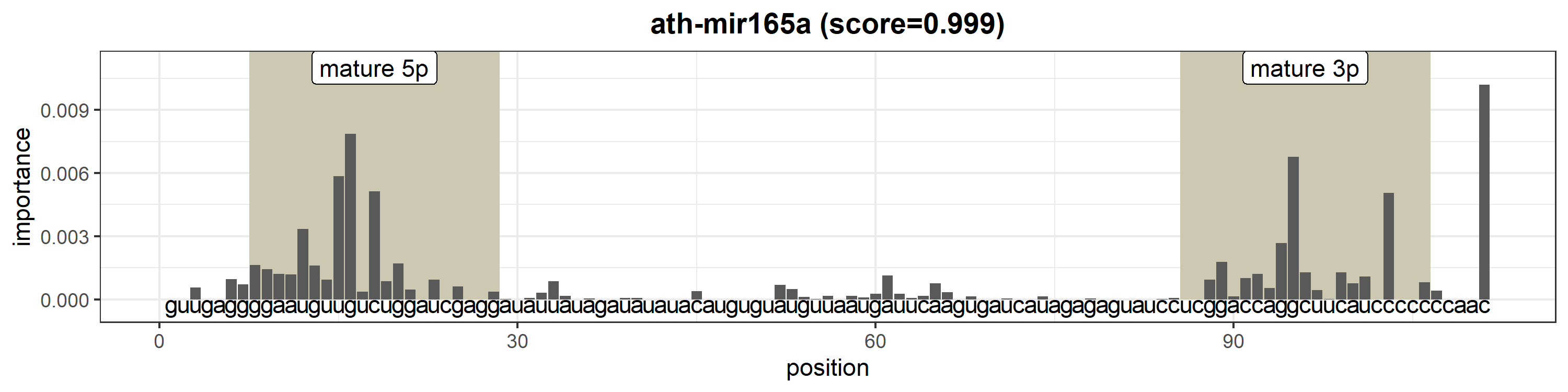

### ath-MIR165b.png

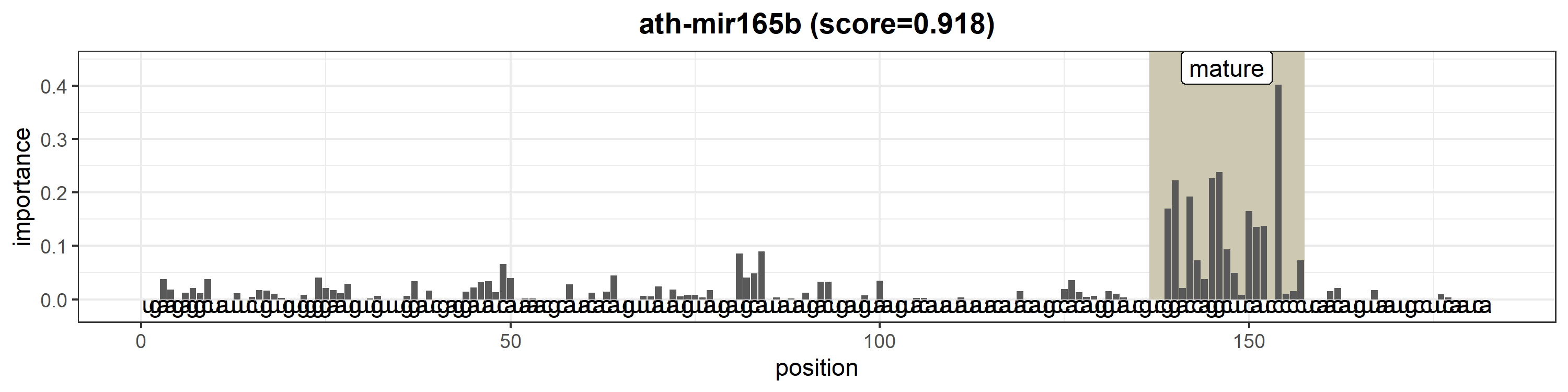

### ath-MIR166b.png

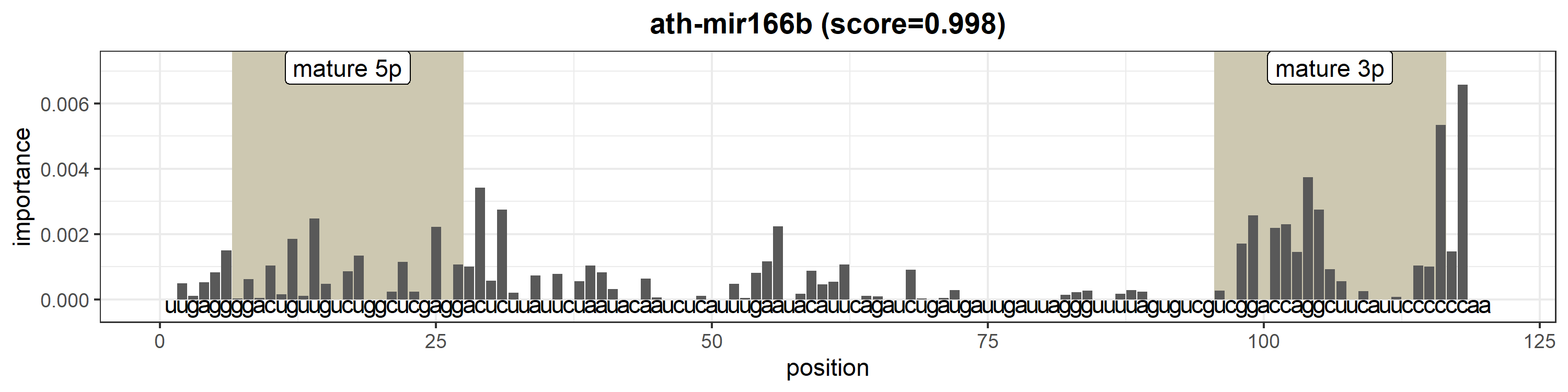

### ath-MIR166c.png

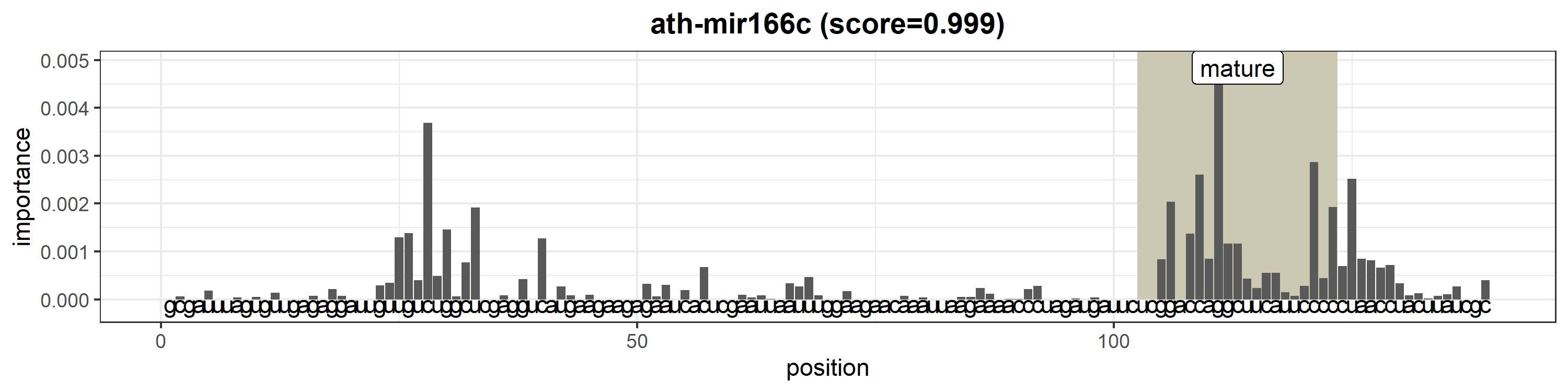

### ath-MIR166d.png

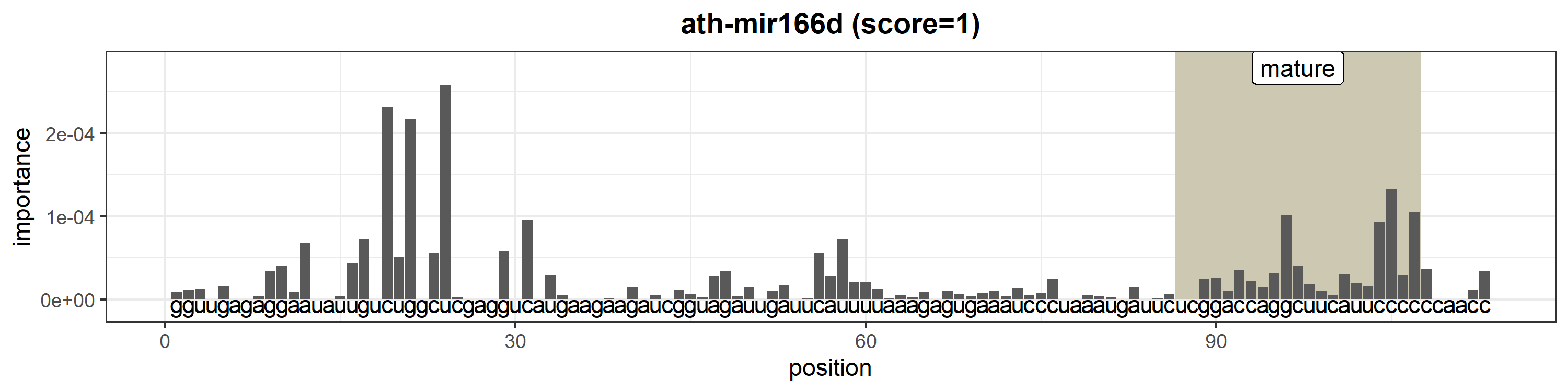

### ath-MIR166e.png

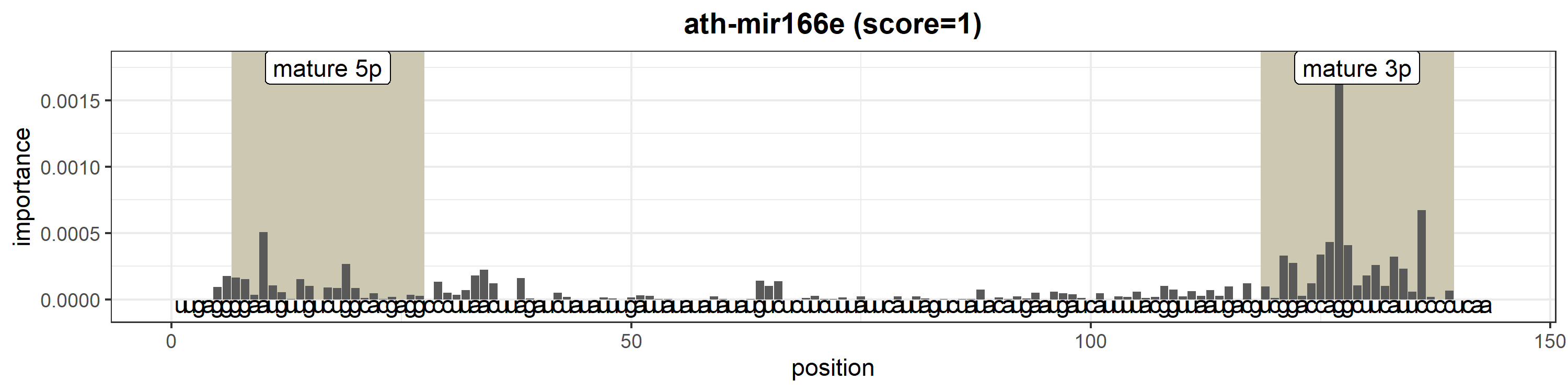

### ath-MIR166f.png

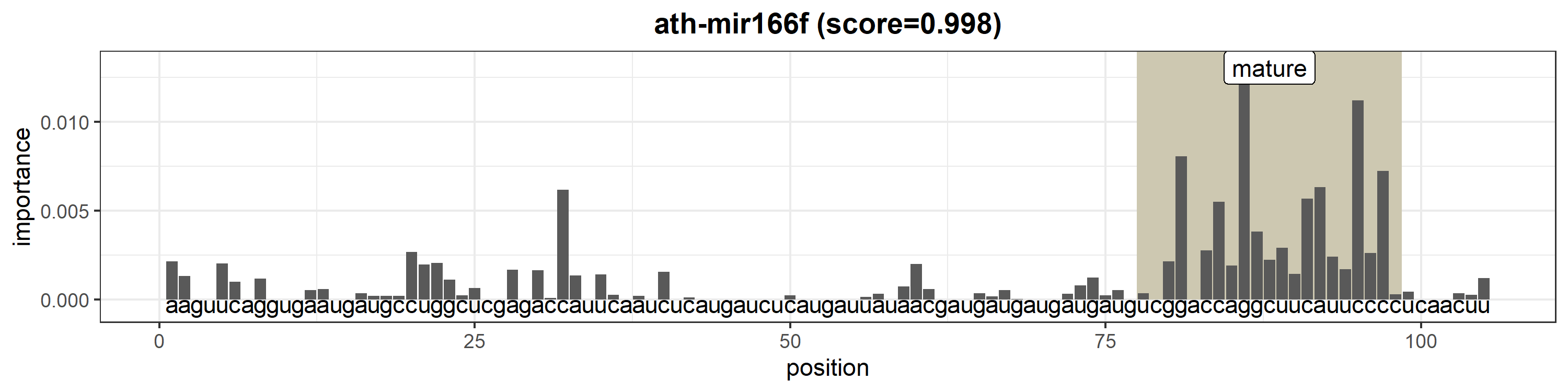

### ath-MIR166g.png

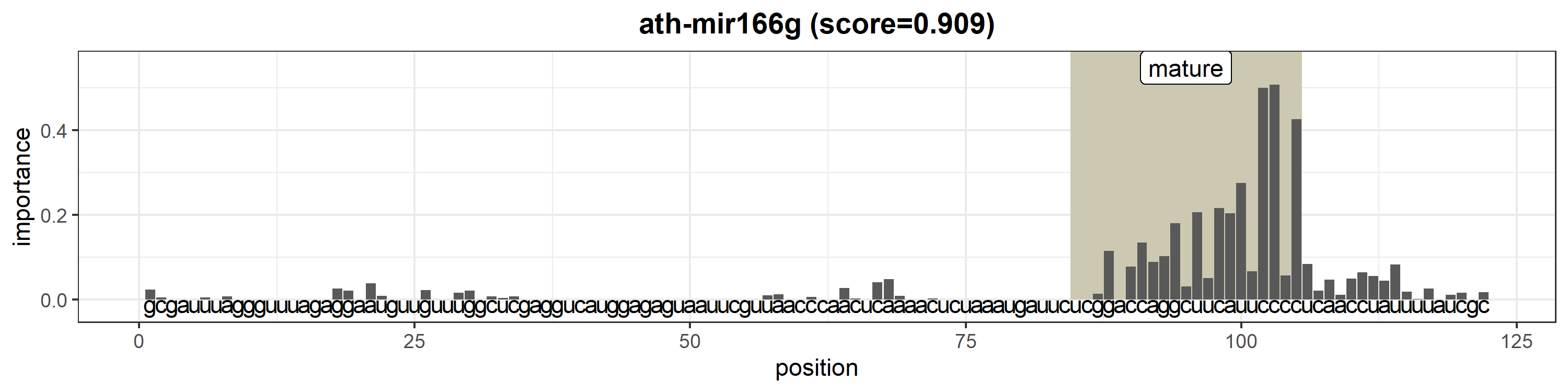

### ath-MIR167a.png

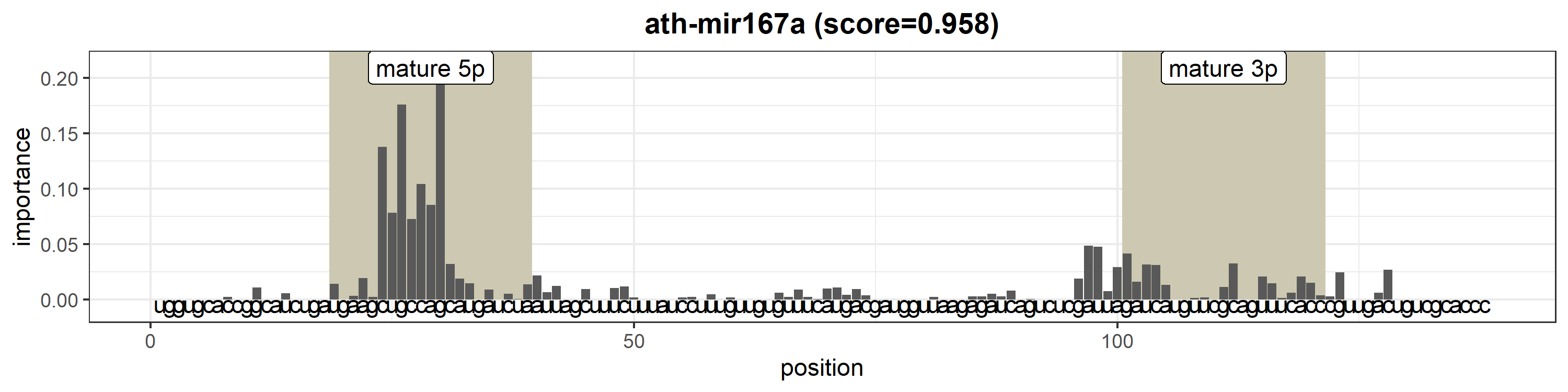

### ath-MIR167b.png

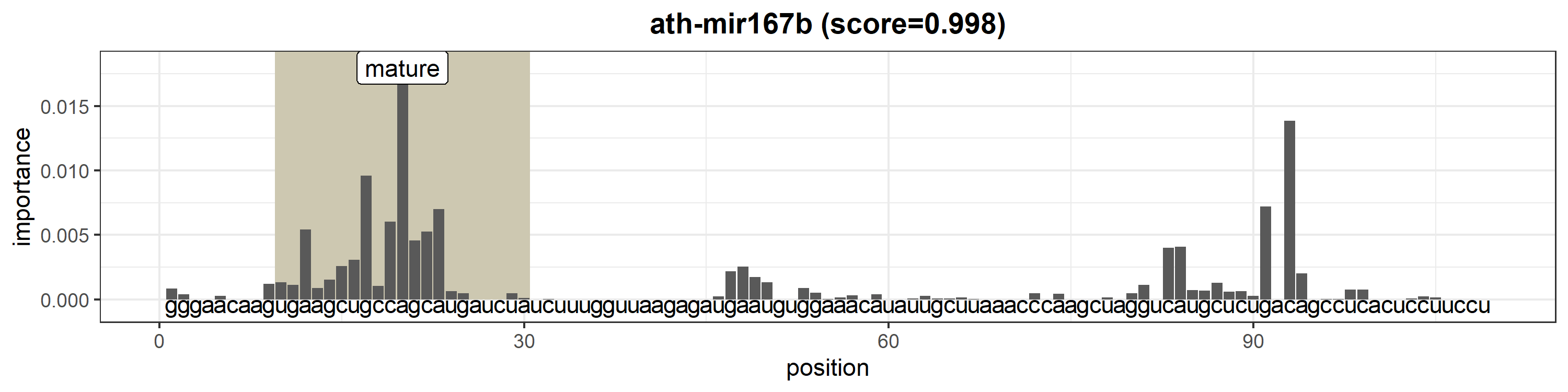

### ath-MIR167c.png

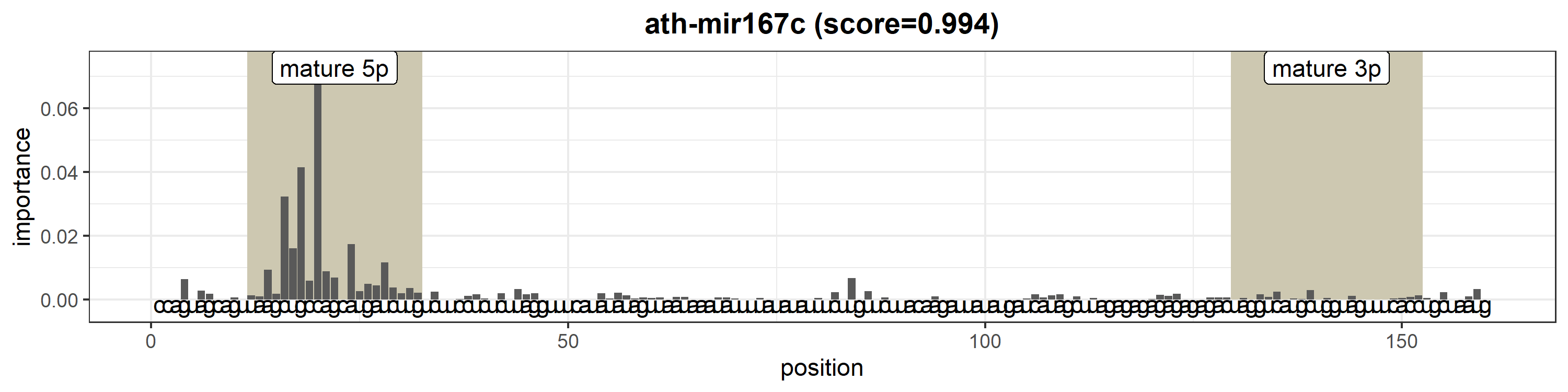

### ath-MIR168a.png

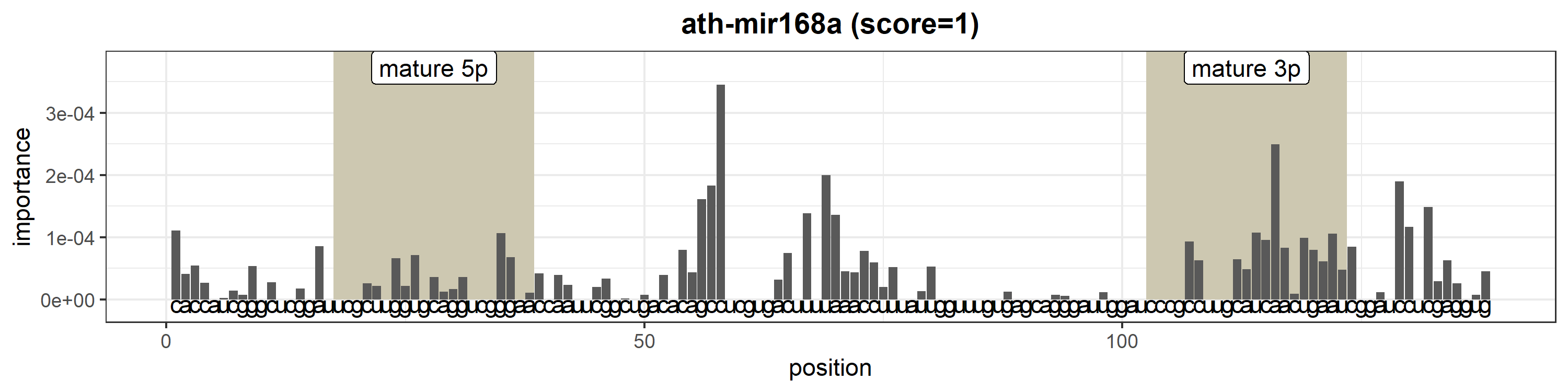

### ath-MIR168b.png

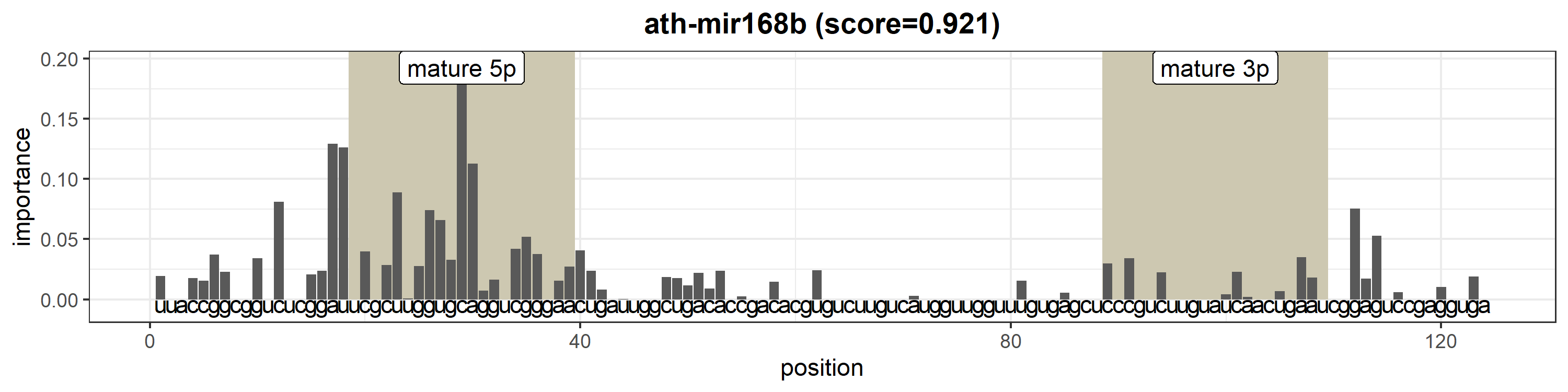

### ath-MIR169b.png

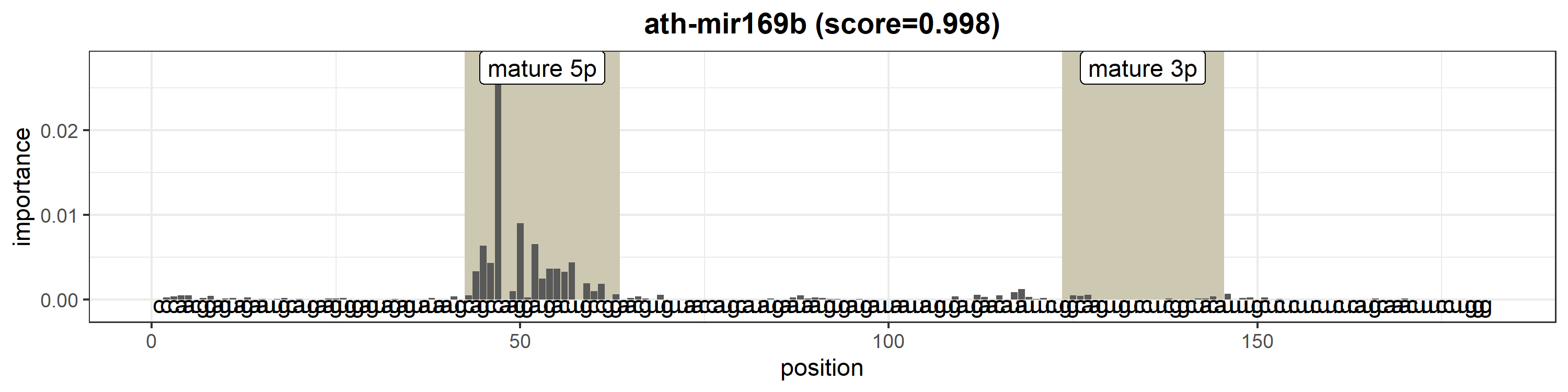

### ath-MIR169d.png

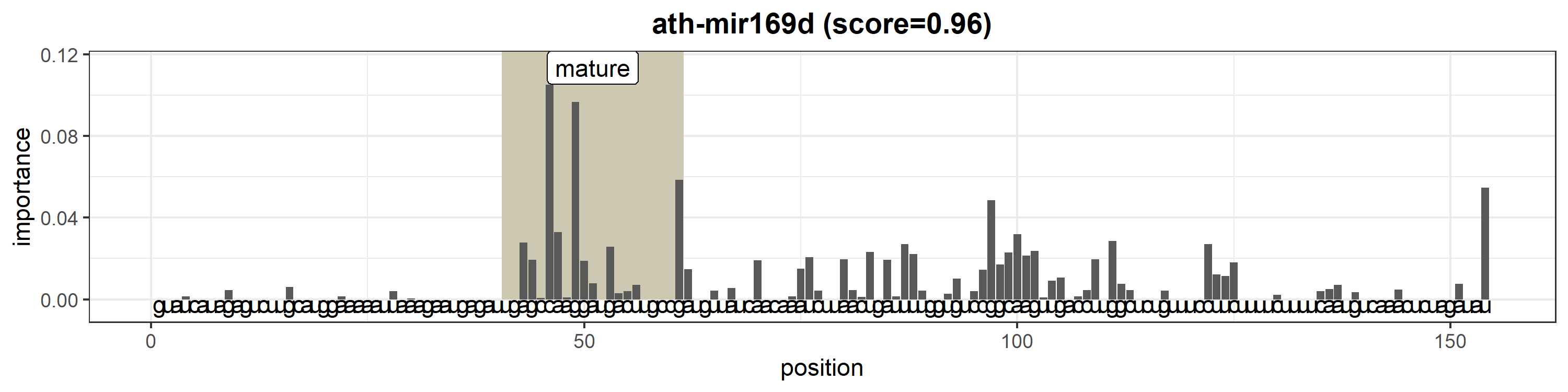

### ath-MIR169e.png

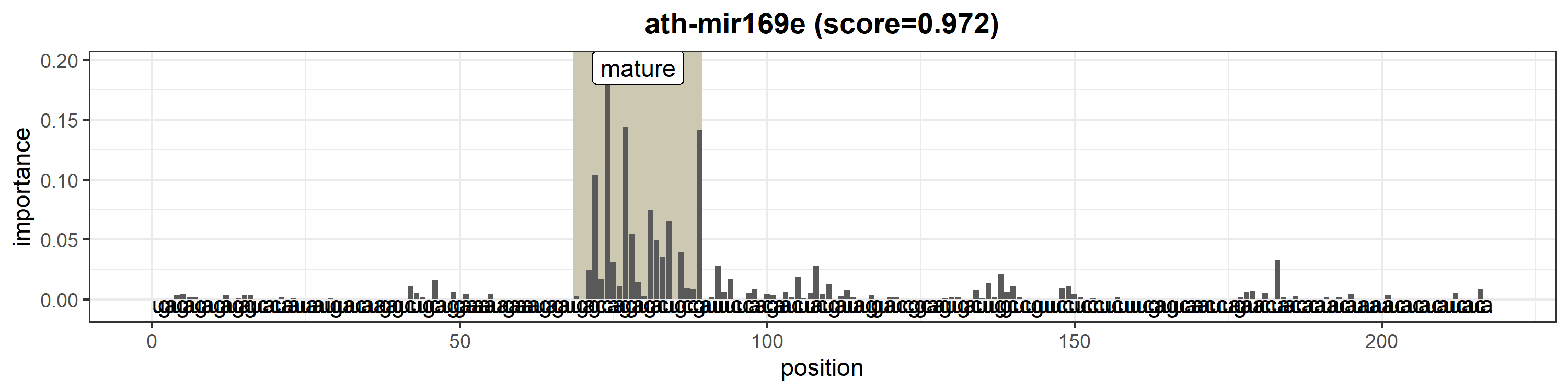

### ath-MIR169f.png

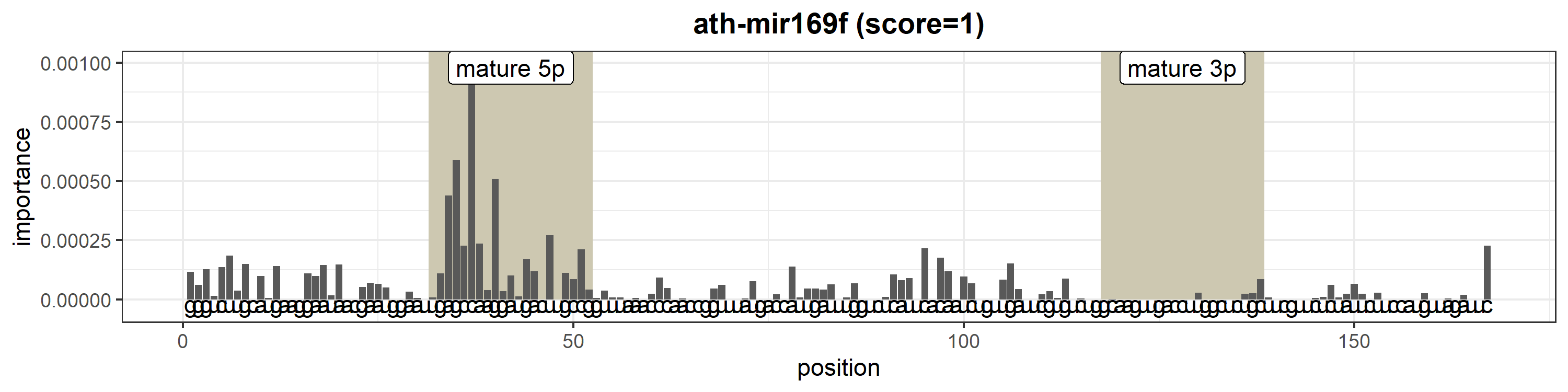

### ath-MIR169h.png

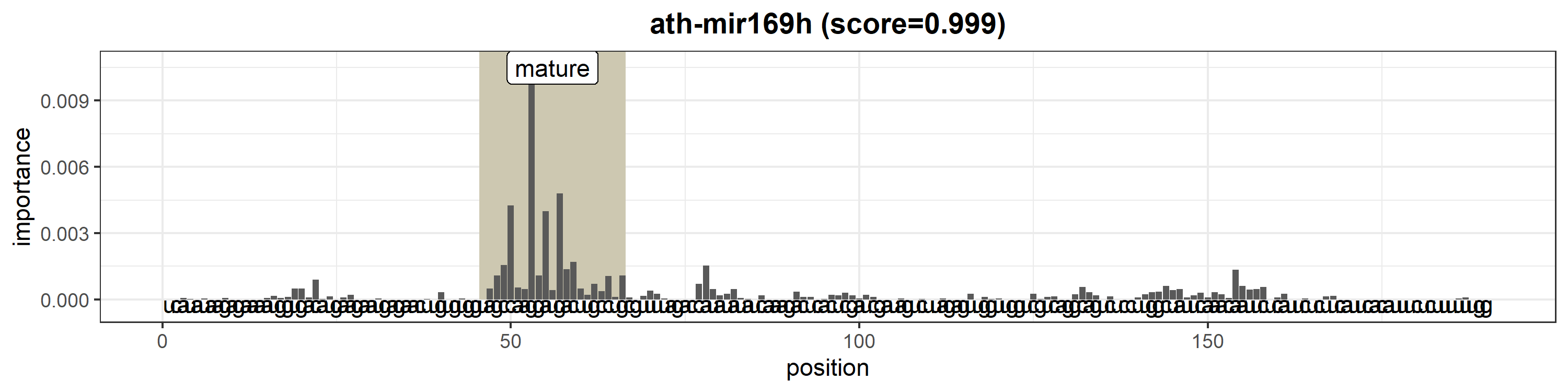

### ath-MIR169i.png

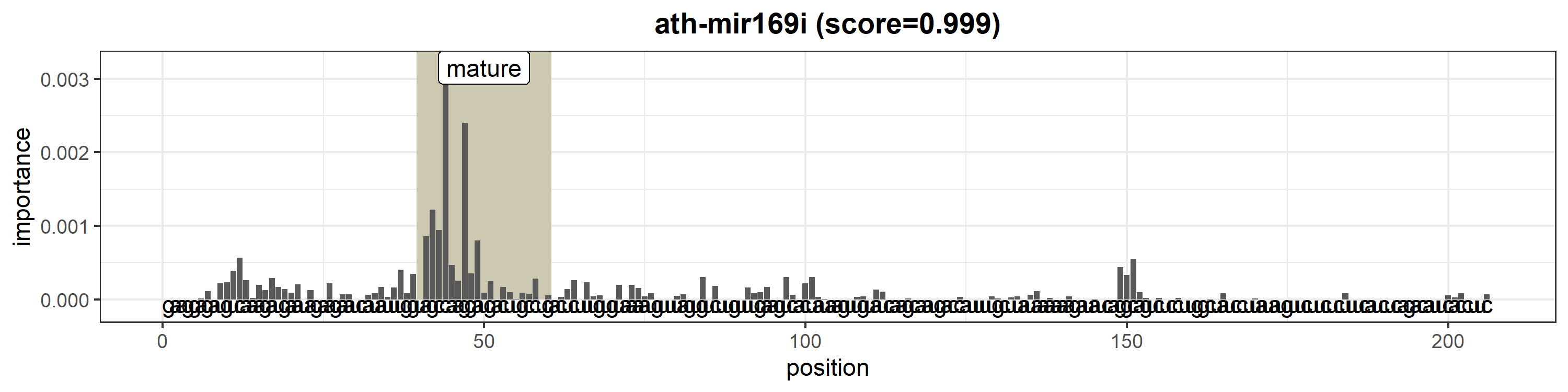

### ath-MIR169j.png

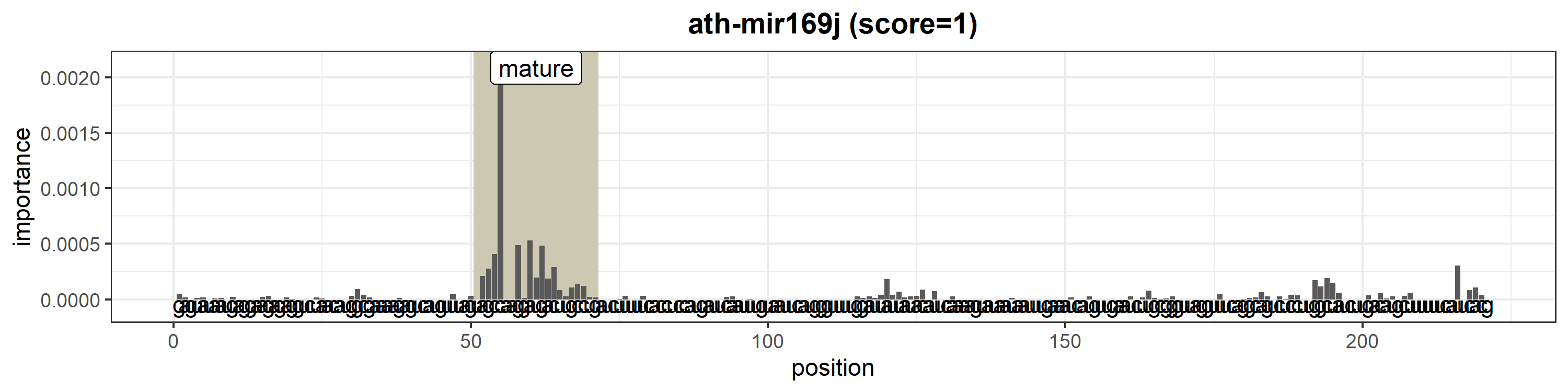

### ath-MIR169k.png

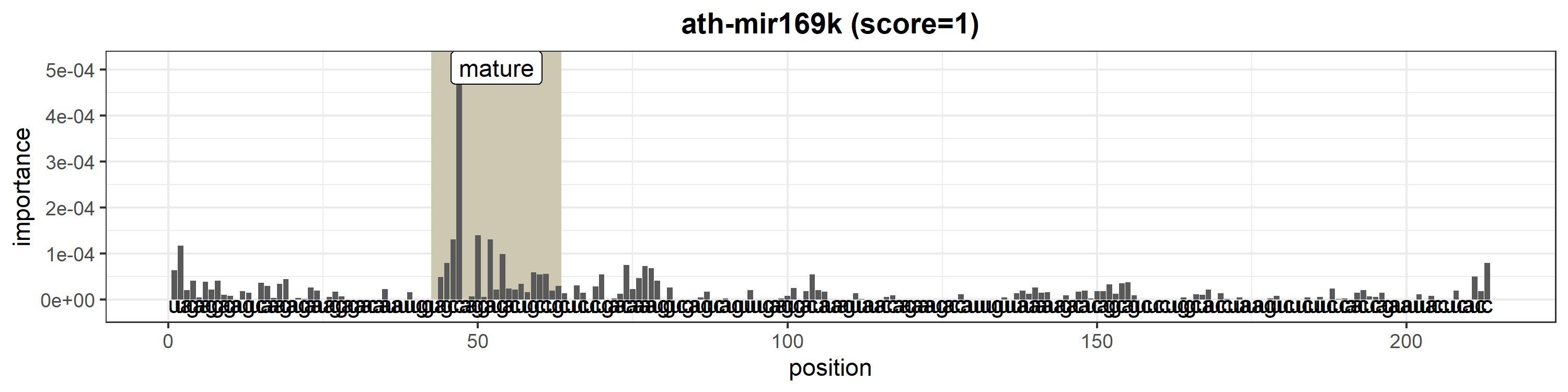

### ath-MIR169l.png

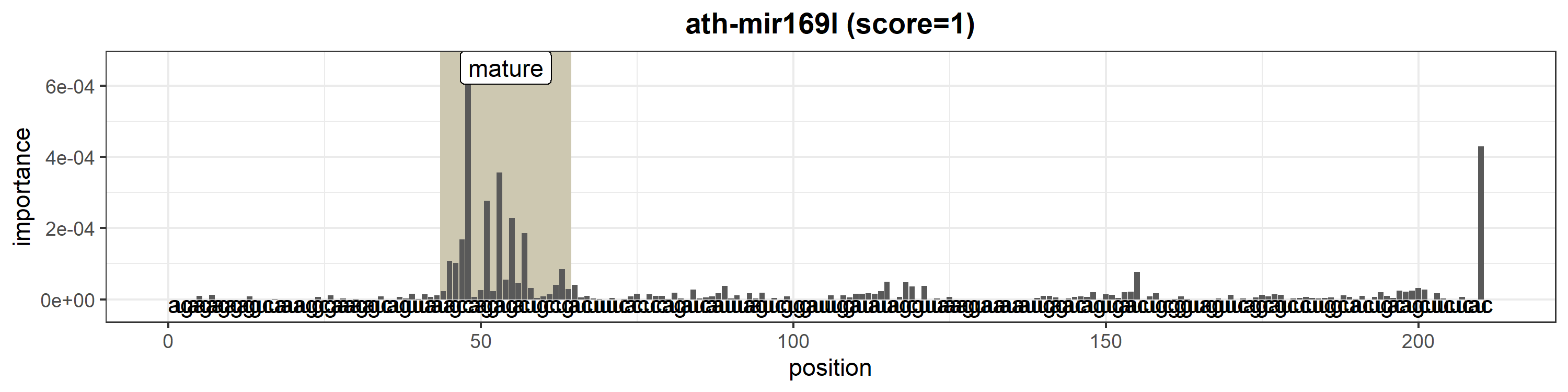

### ath-MIR169m.png

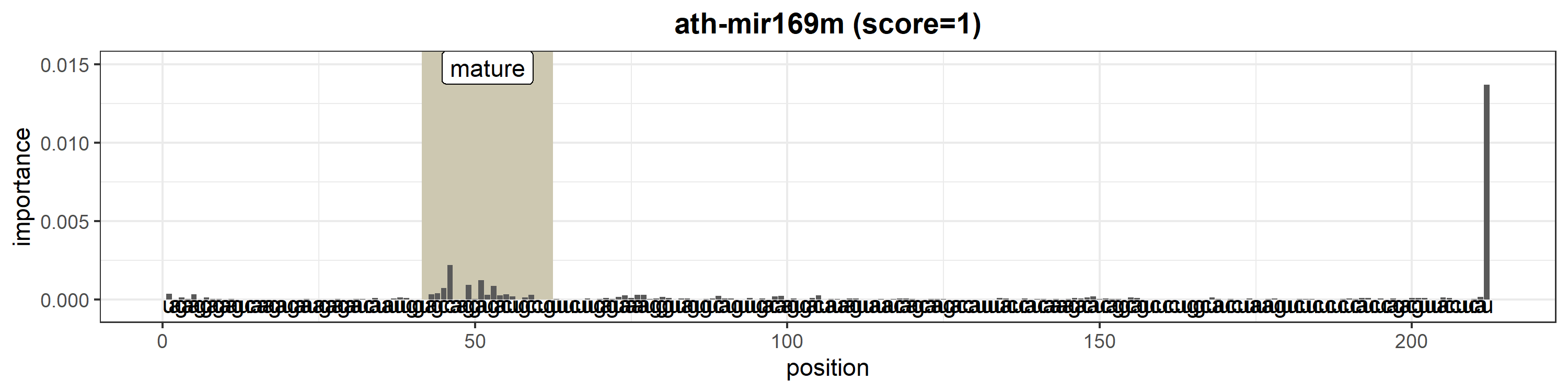

### ath-MIR170.png

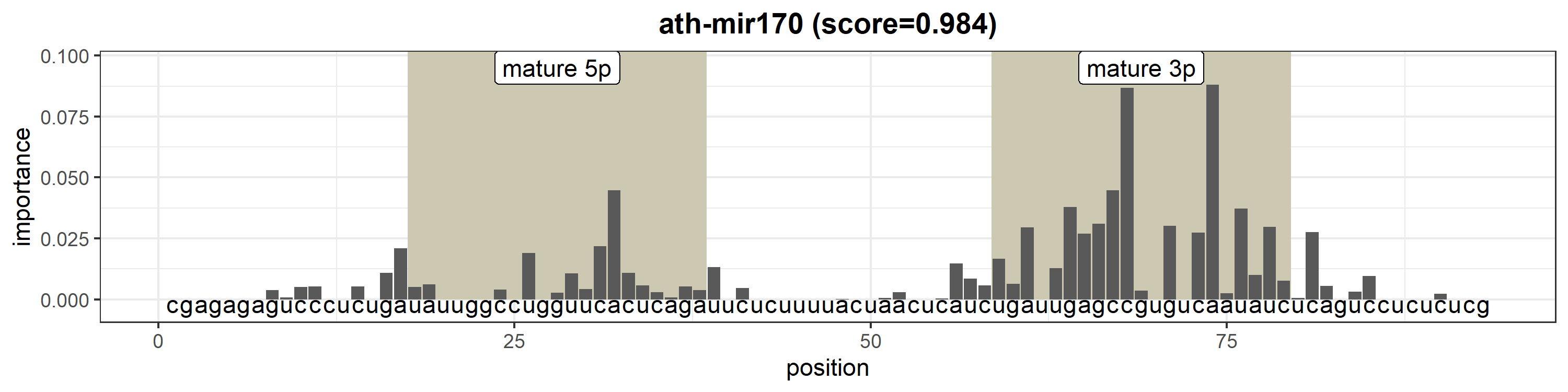

### ath-MIR171a.png

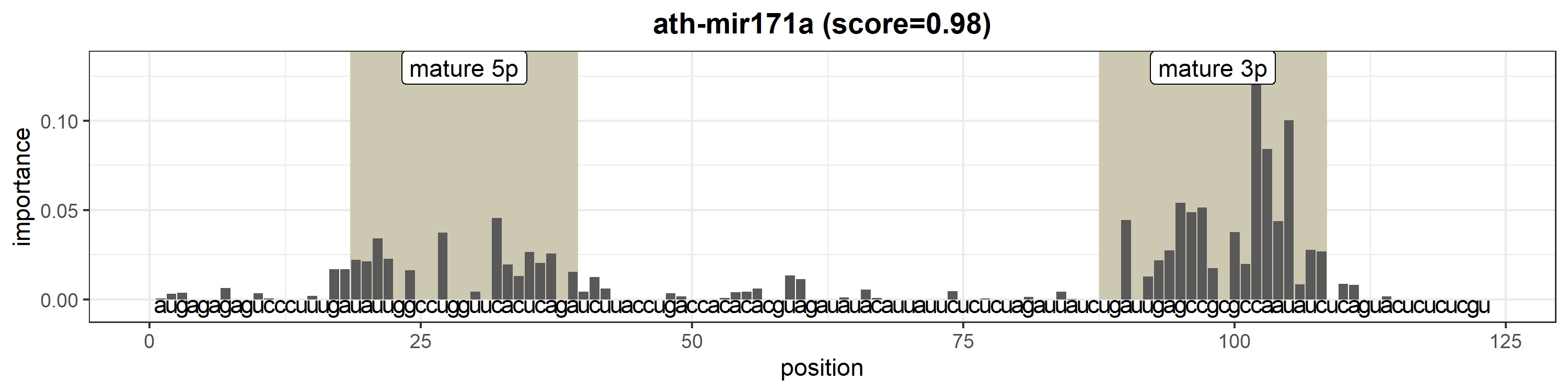

### ath-MIR171b.png

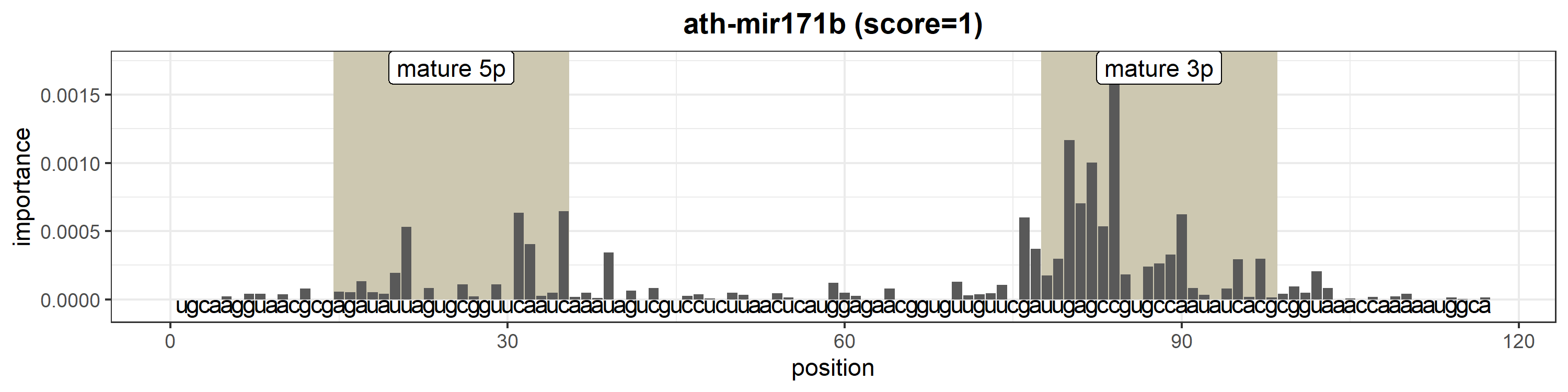

### ath-MIR171c.png

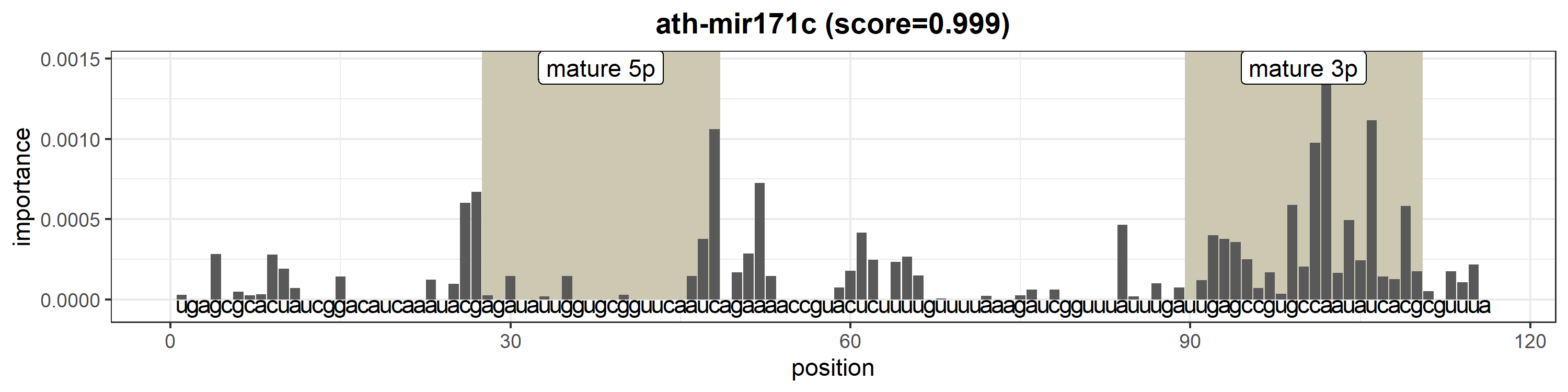

### ath-MIR172a.png

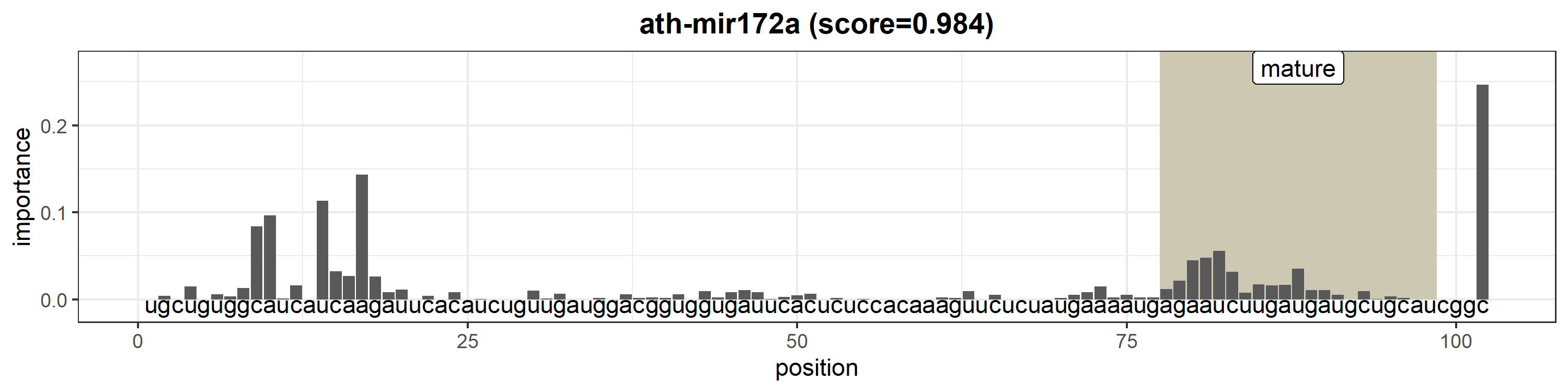

### ath-MIR172b.png

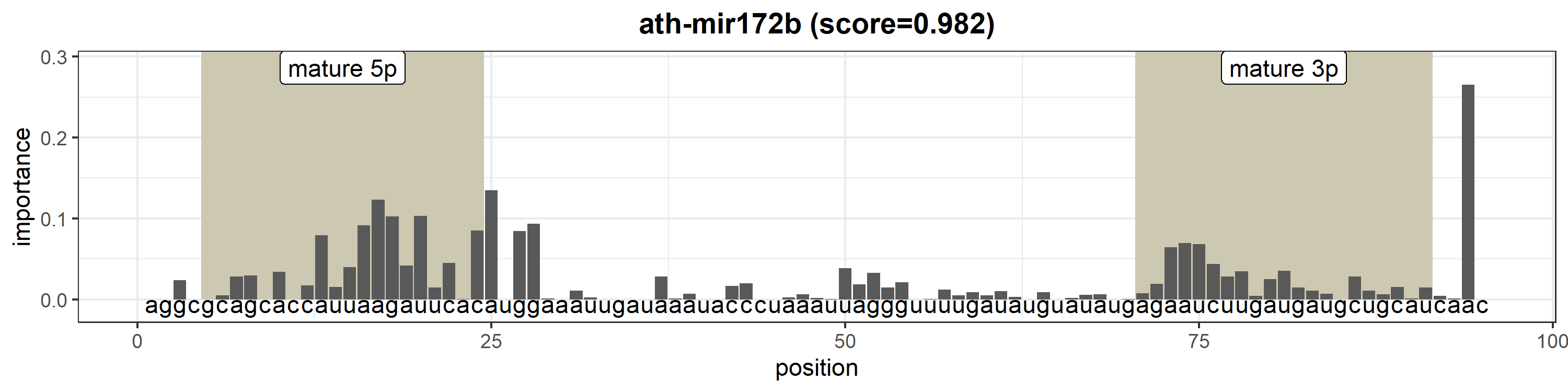

### ath-MIR172c.png

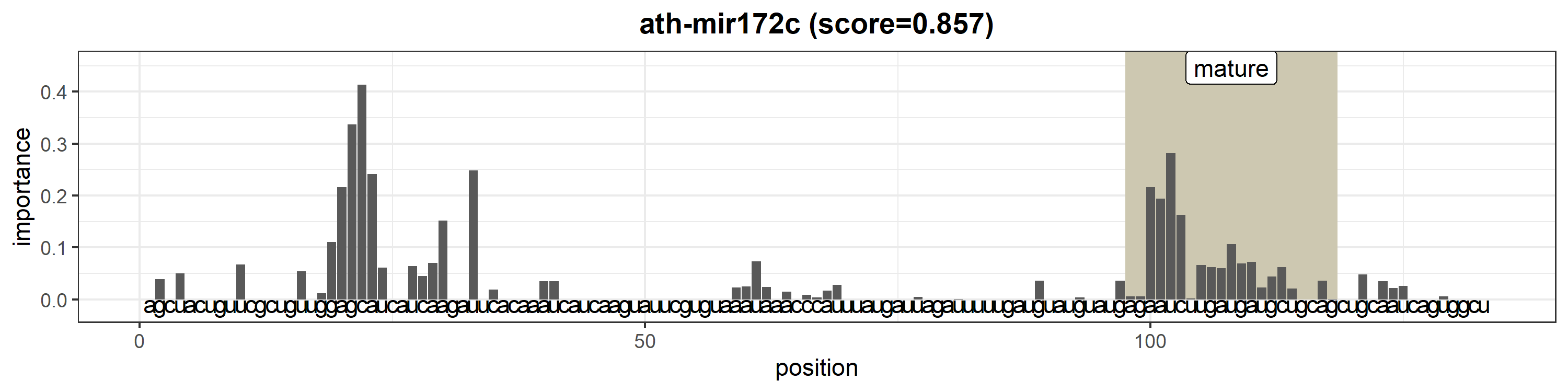

### ath-MIR172d.png

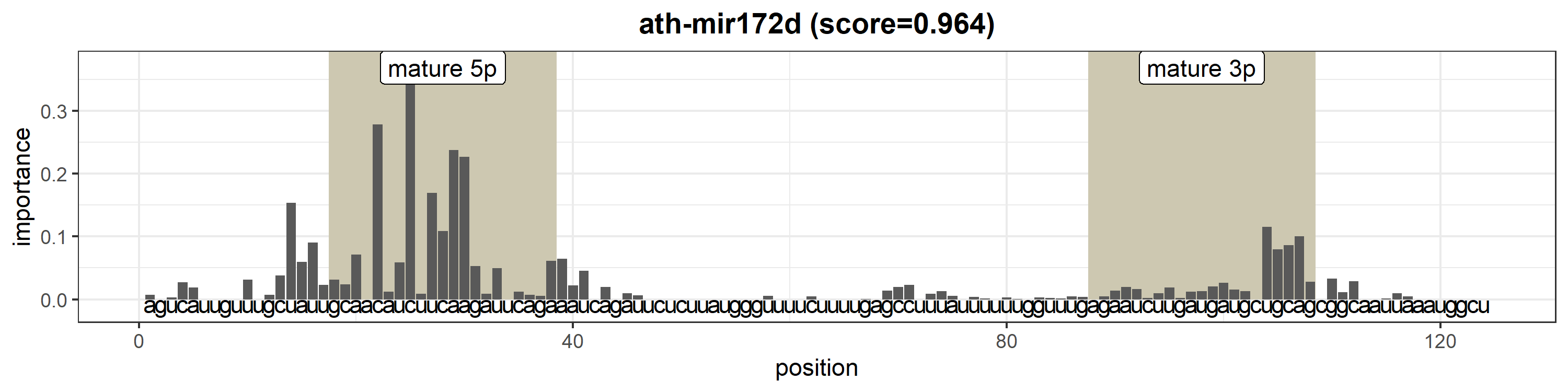

### ath-MIR172e.png

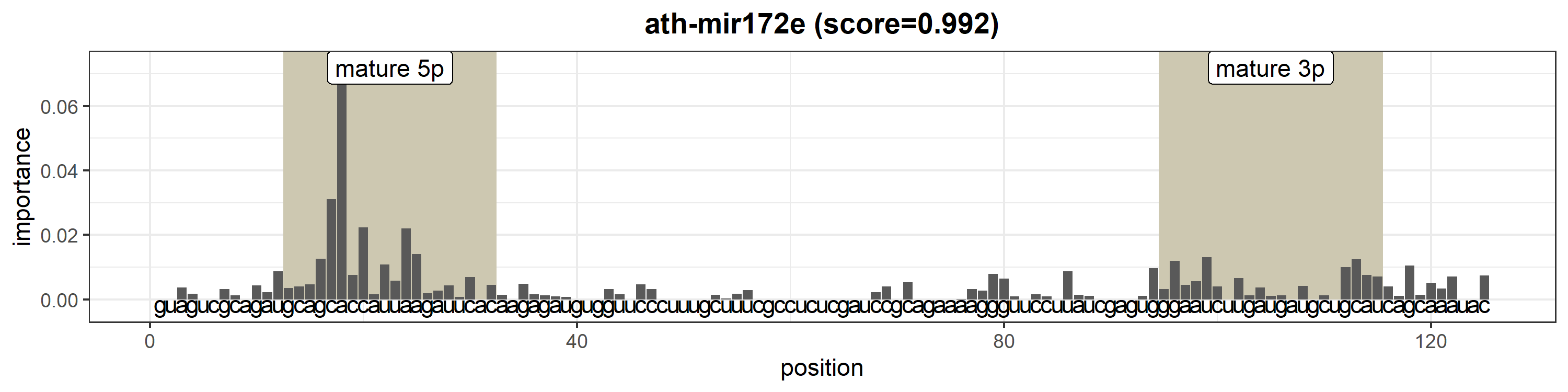

### ath-MIR173.png

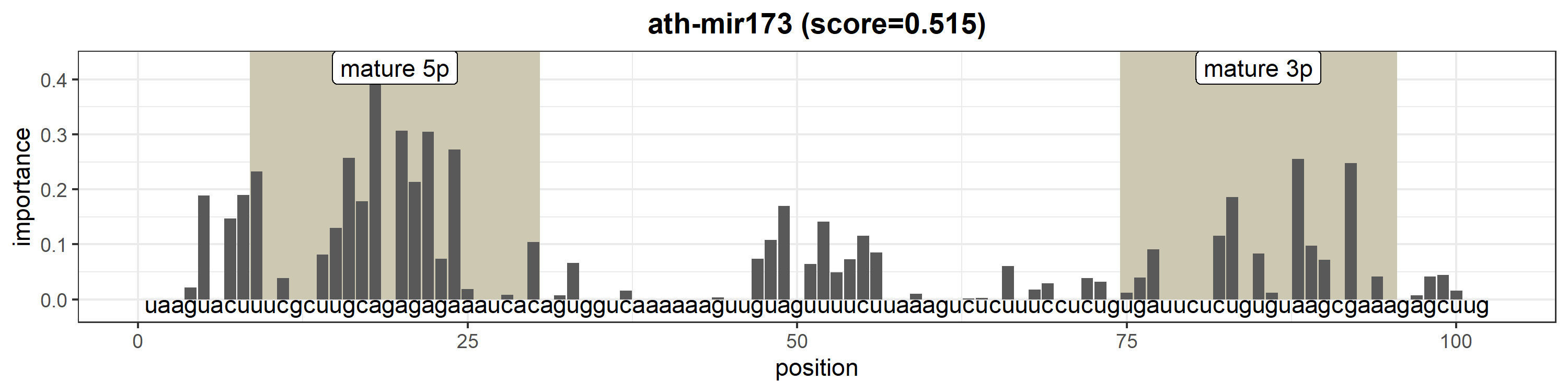

### ath-MIR1886.png

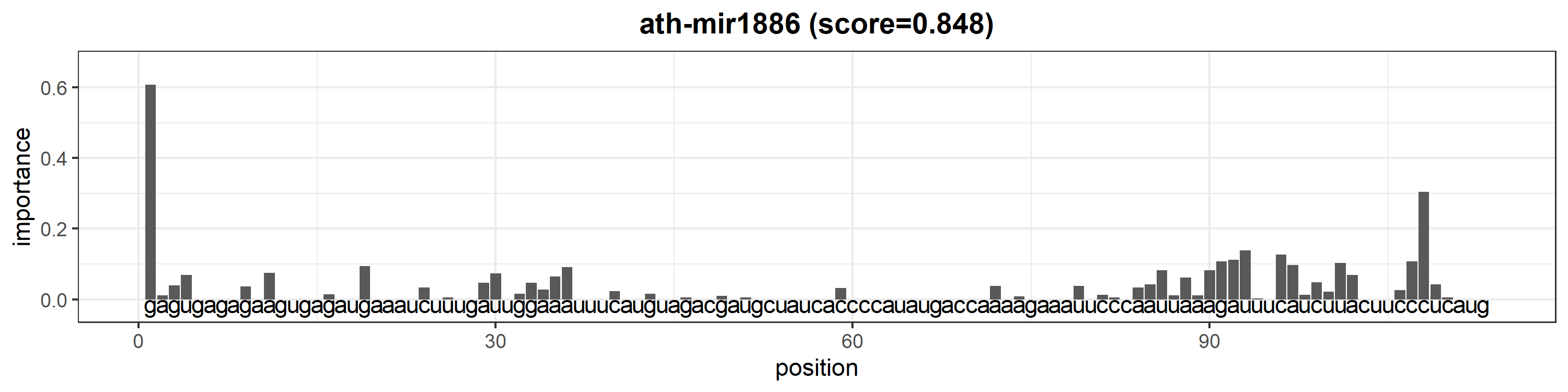

### ath-MIR1887.png

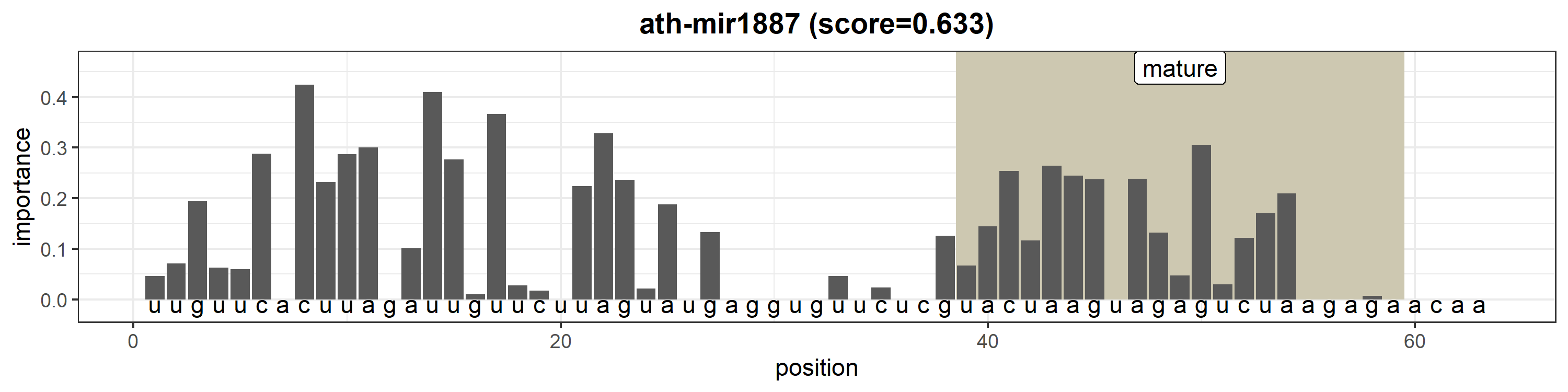

### ath-MIR1888a.png

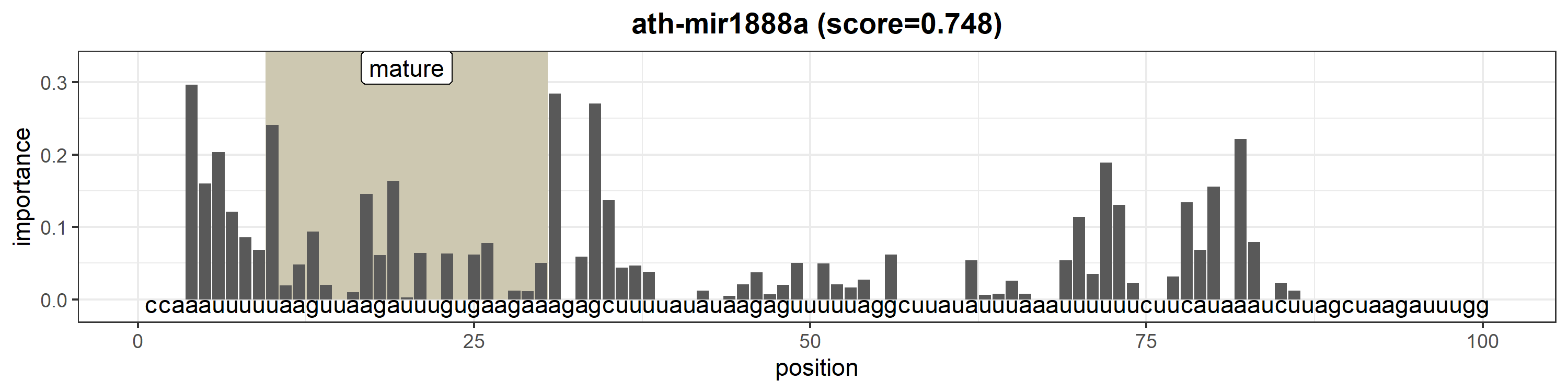

### ath-MIR1888b.png

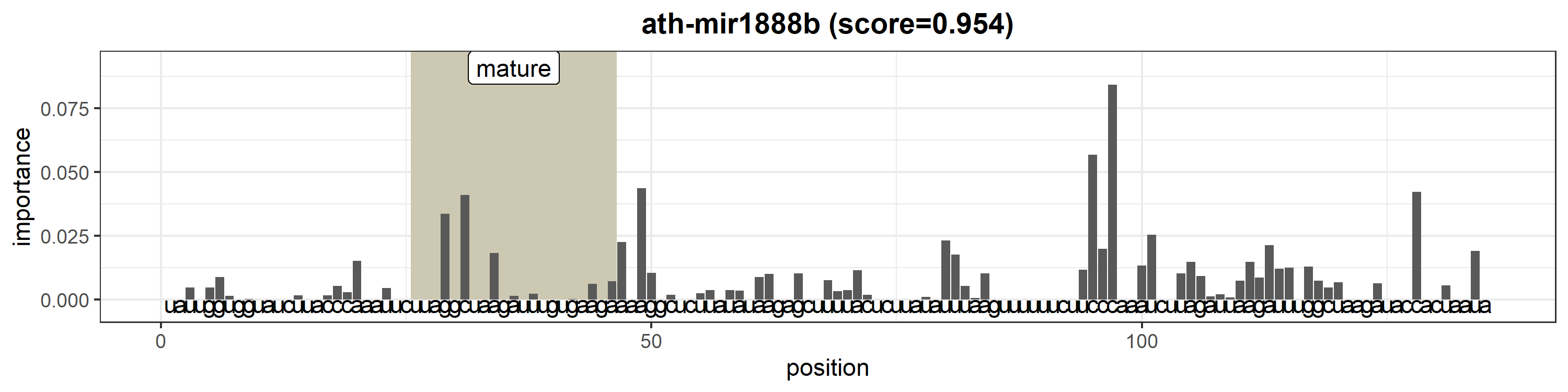

### ath-MIR2111a.png

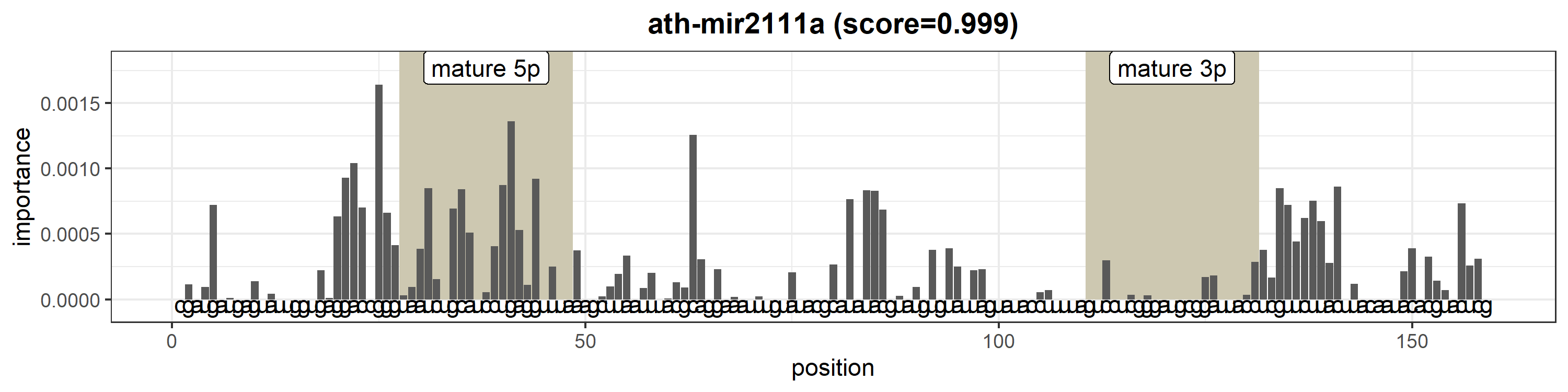

### ath-MIR2111b.png

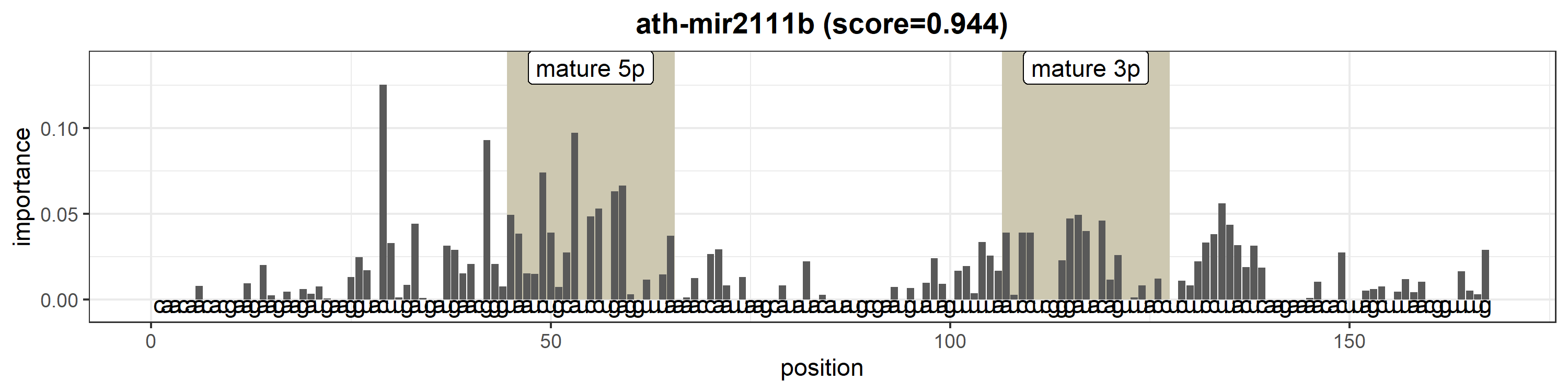

### ath-MIR2112.png

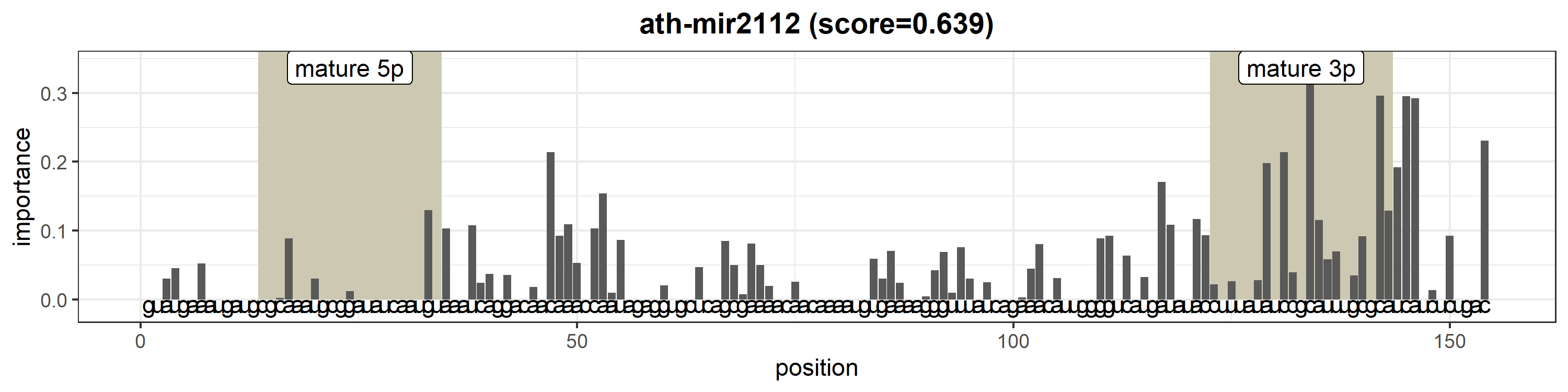

### ath-MIR2933a.png

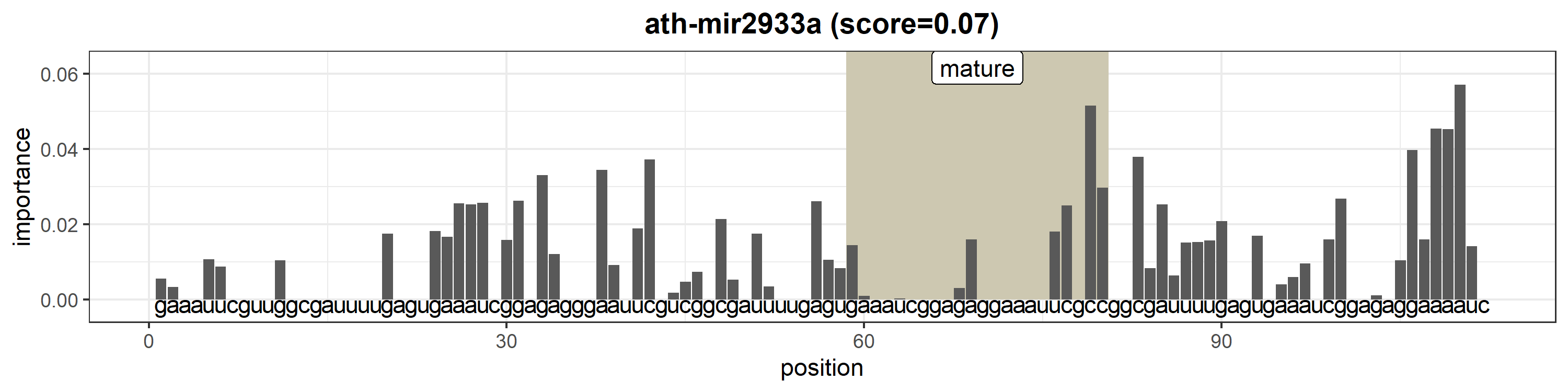

### ath-MIR2933b.png

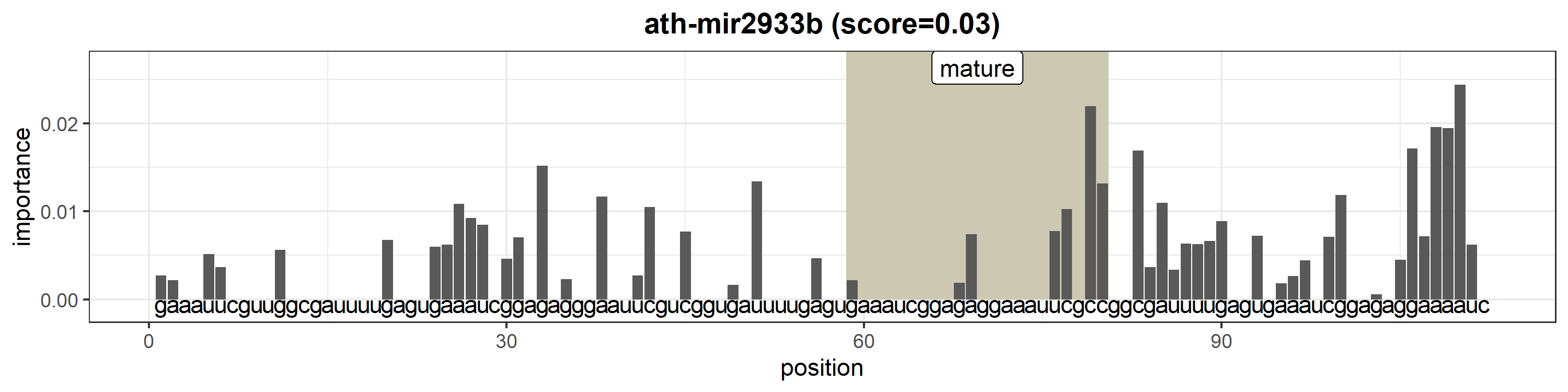

### ath-MIR2934.png

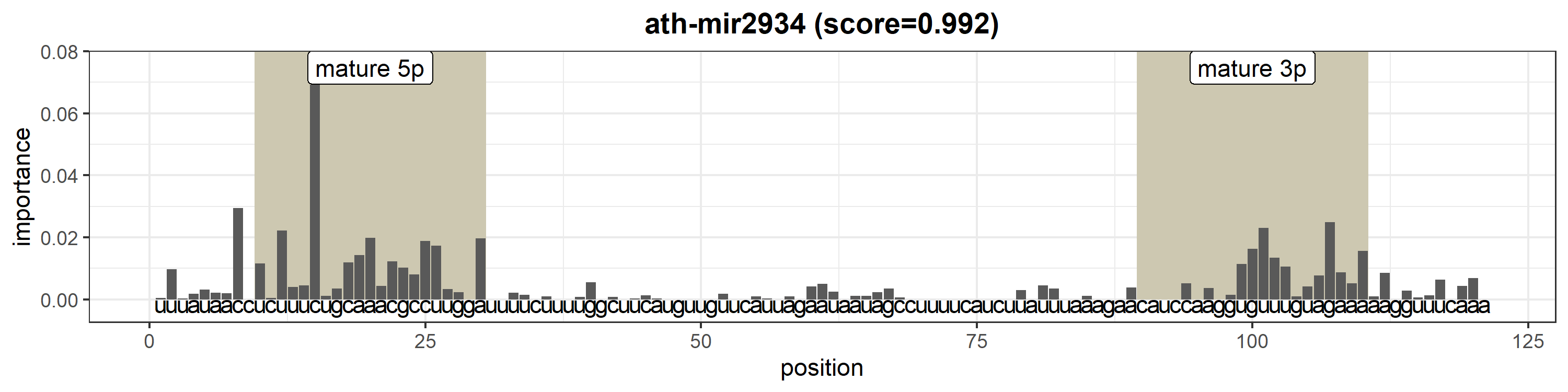

### ath-MIR10515.png

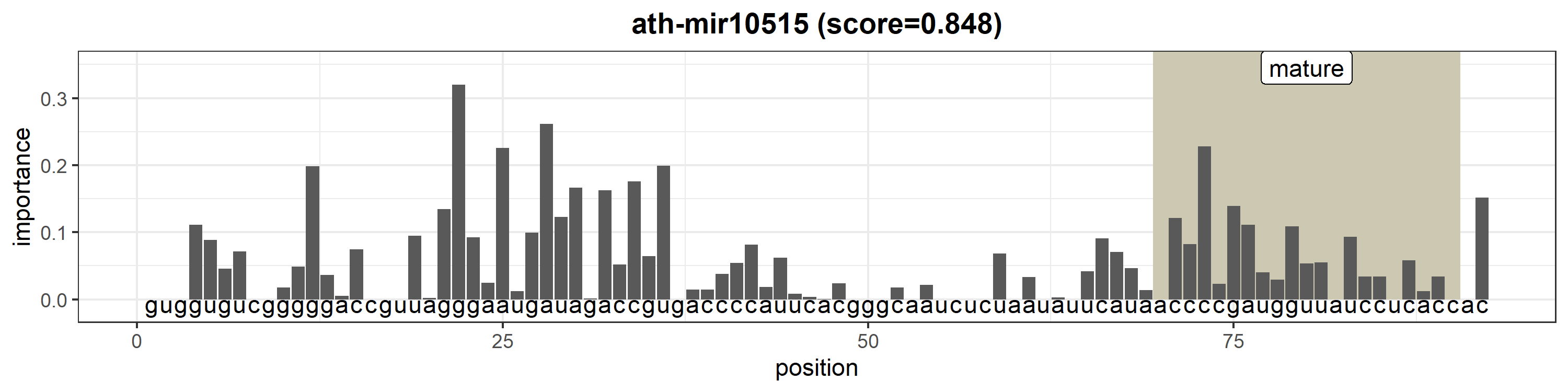
