## Supplementary figures and images for "High precision in microRNA prediction: a novel genome-wide approach based on convolutional deep residual networks"

### aga-bantam.png

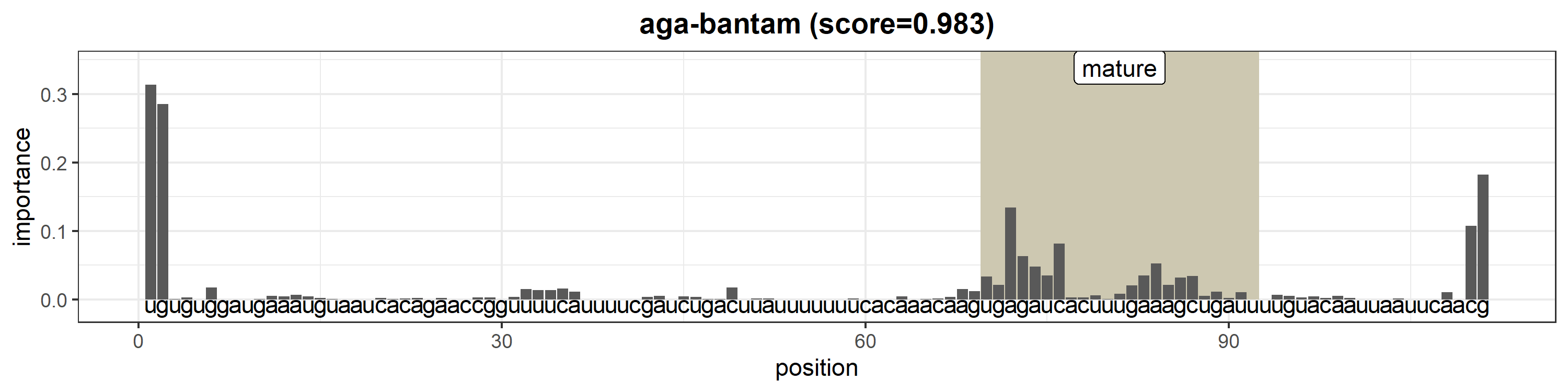

### aga-let-7.png

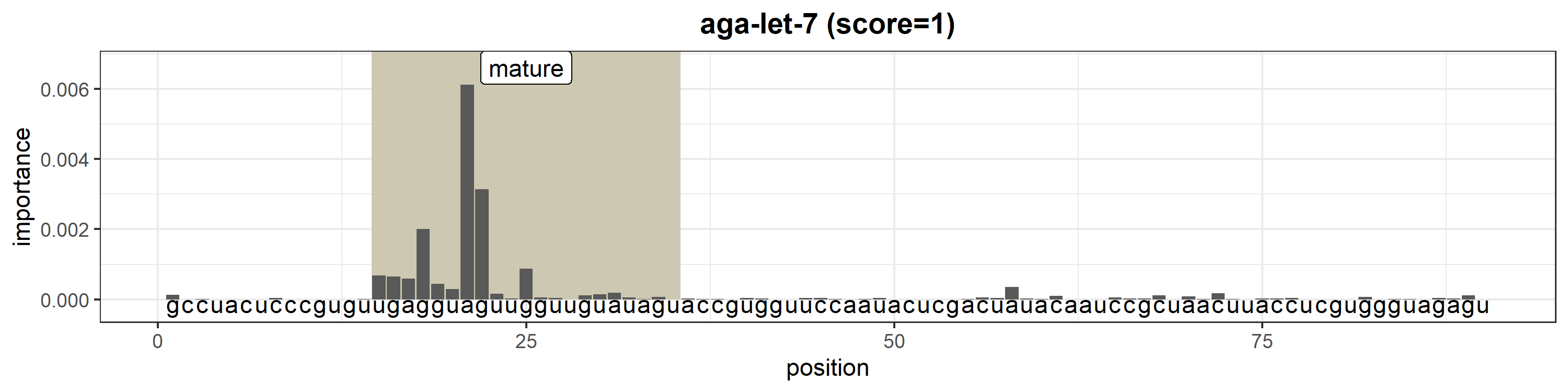

### aga-mir-1.png

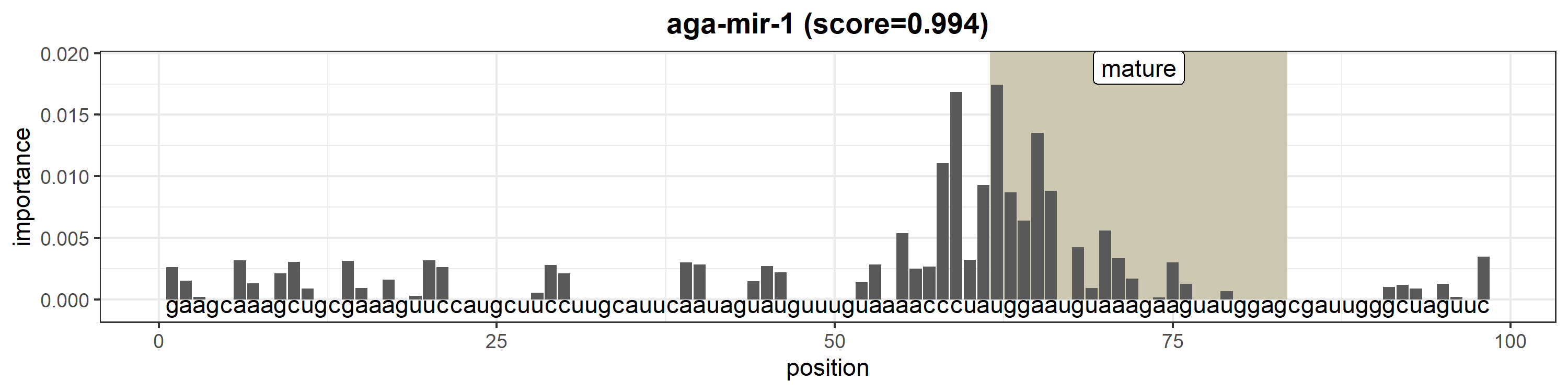

### aga-mir-2-1.png

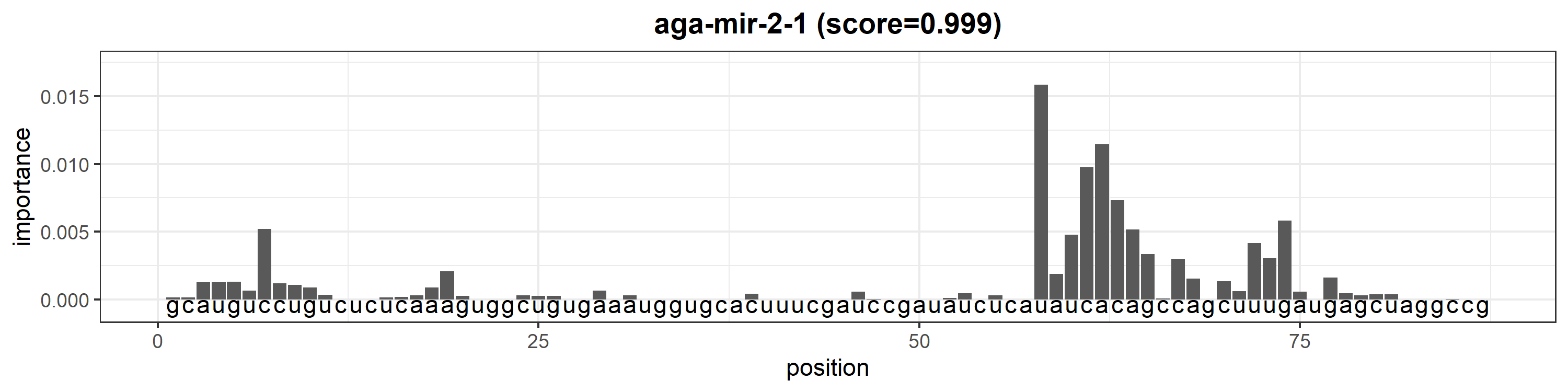

### aga-mir-2-2.png

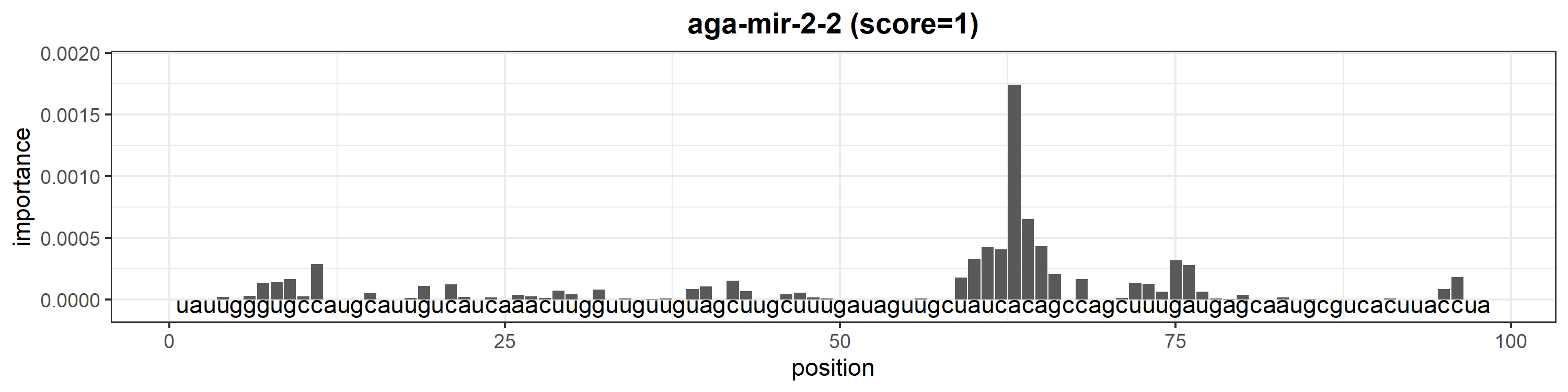

### aga-mir-2b.png

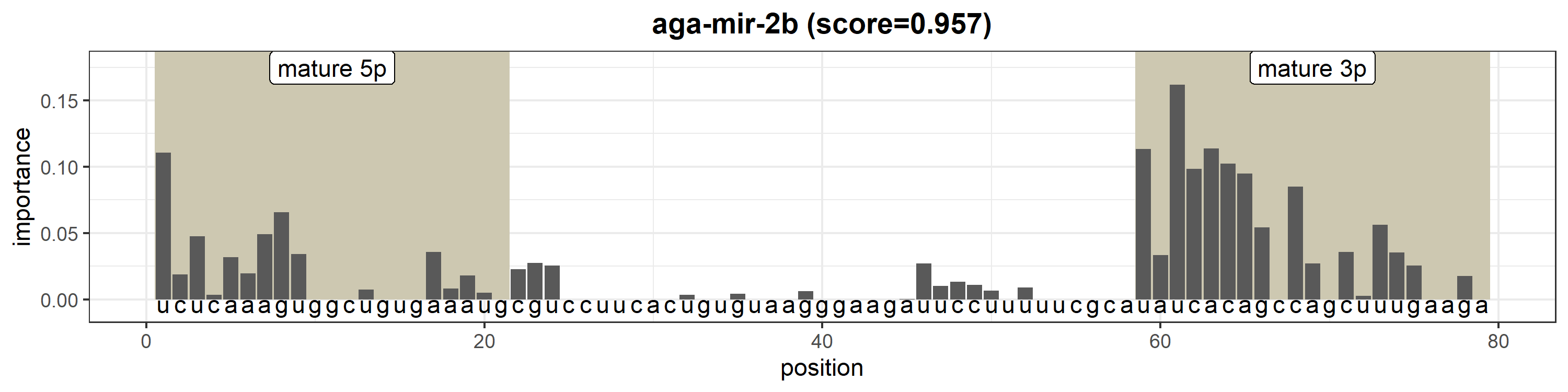

### aga-mir-2c.png

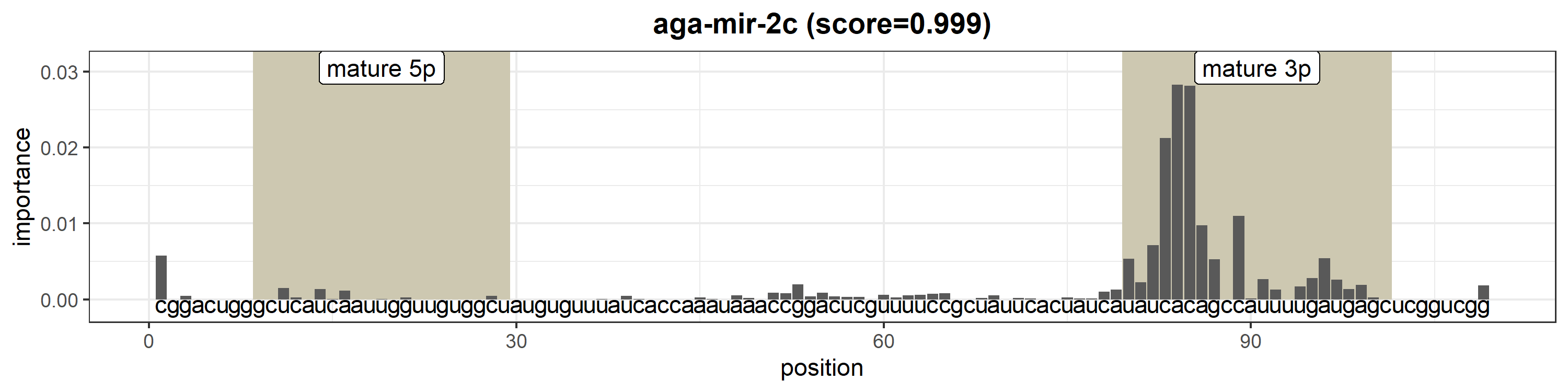

### aga-mir-10.png

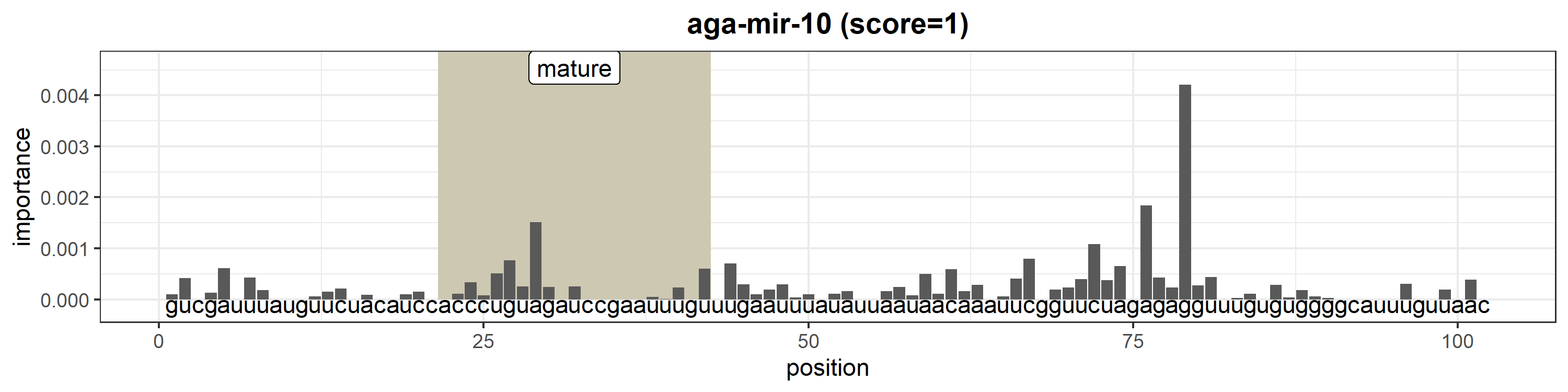

### aga-mir-11.png

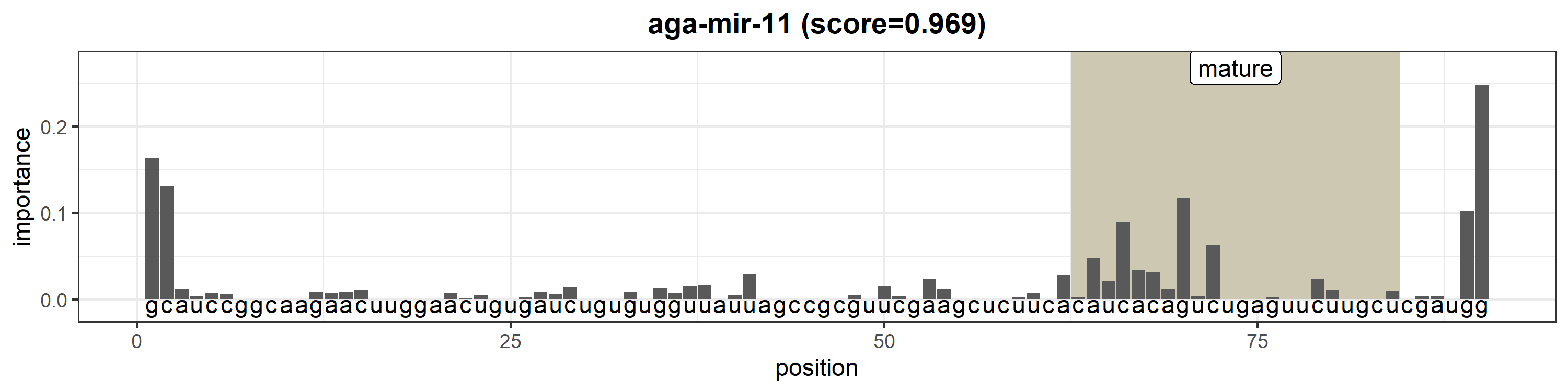

### aga-mir-12.png

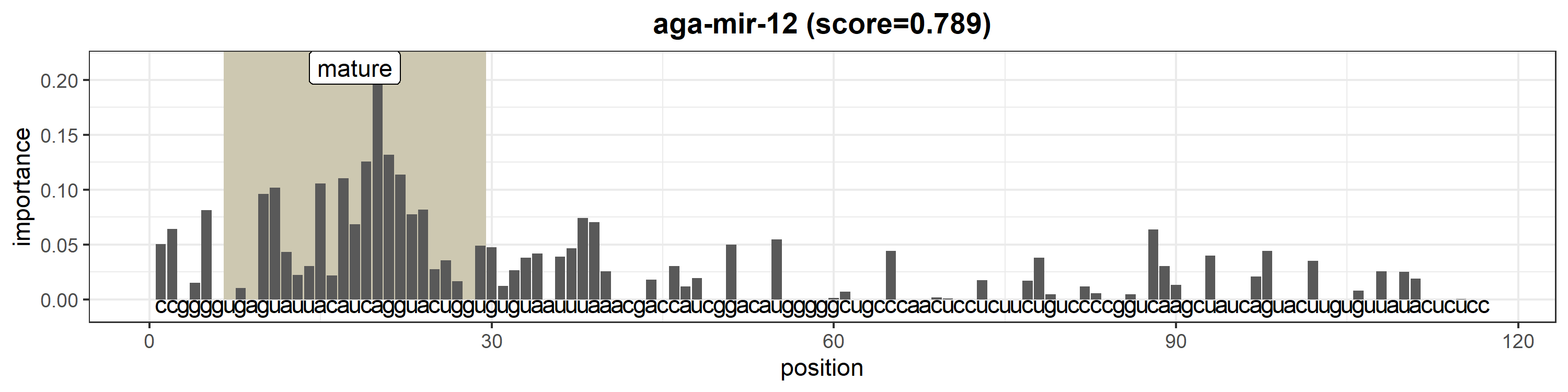

### aga-mir-13b.png

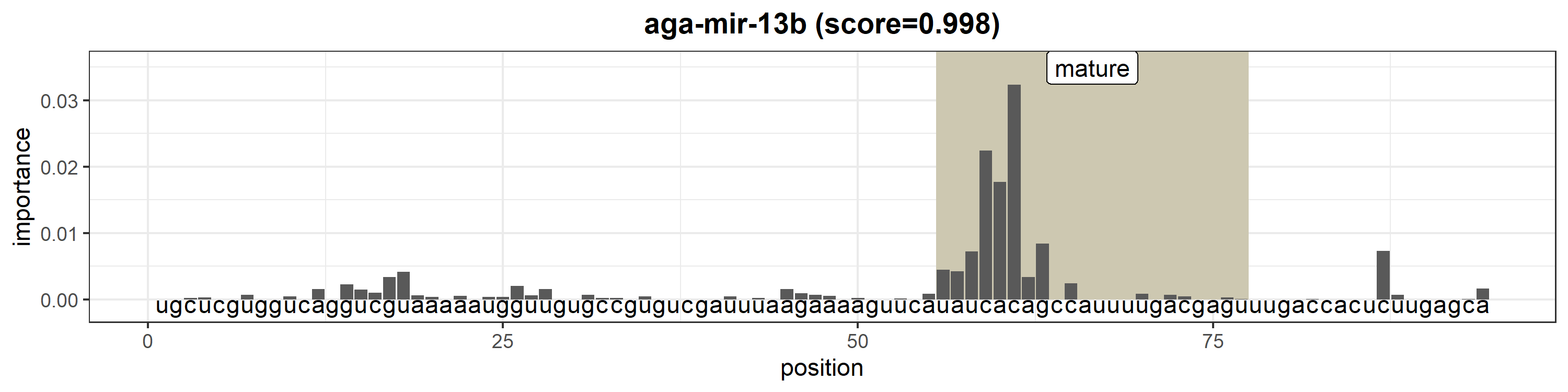

### aga-mir-14.png

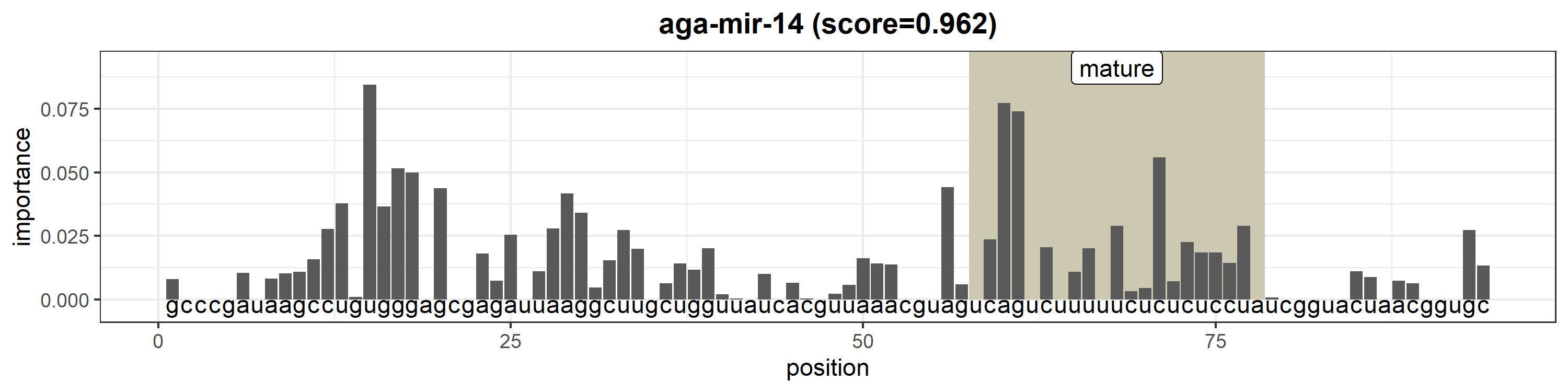

### aga-mir-31.png

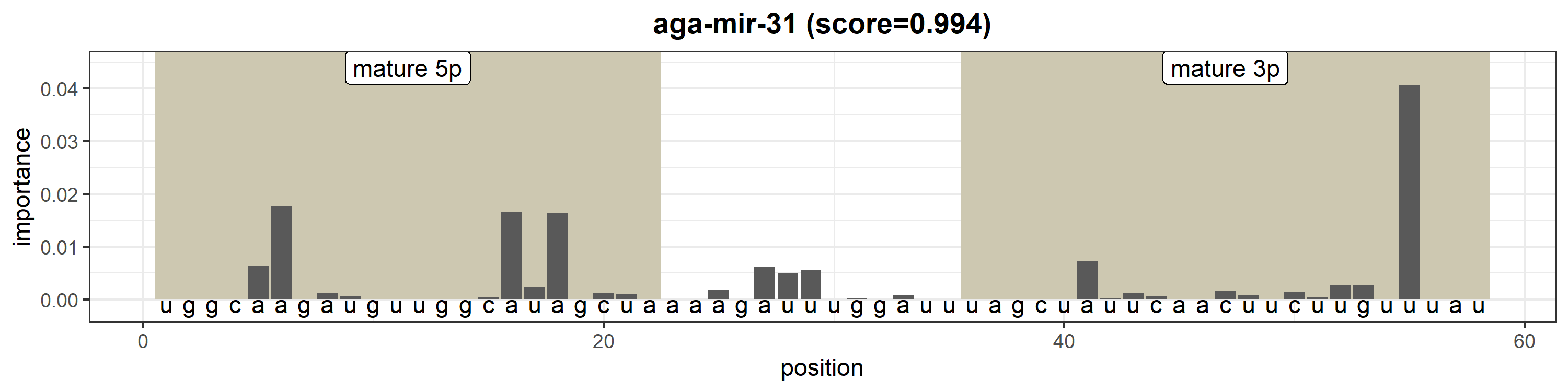

### aga-mir-100.png

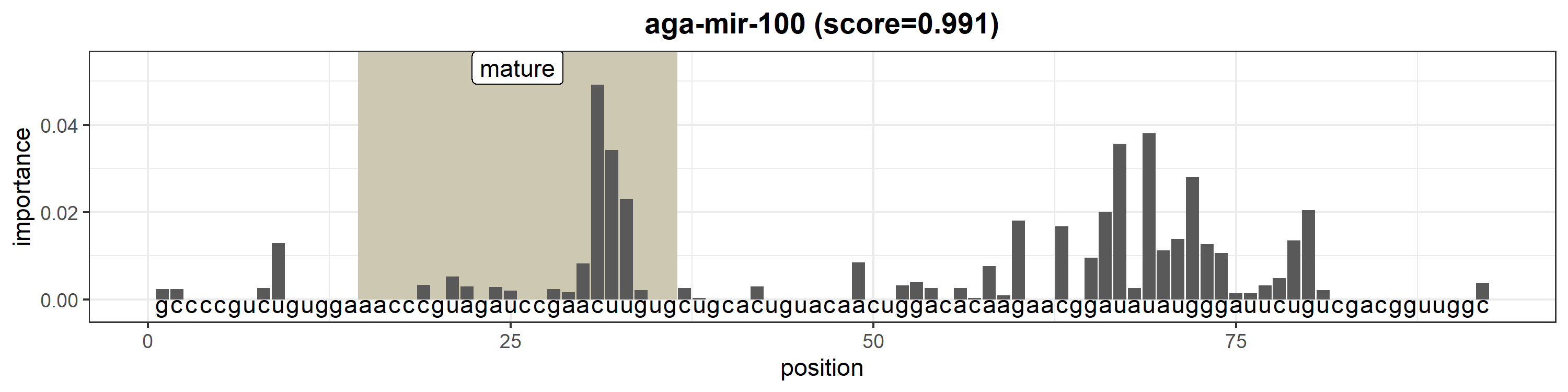

### aga-mir-124.png

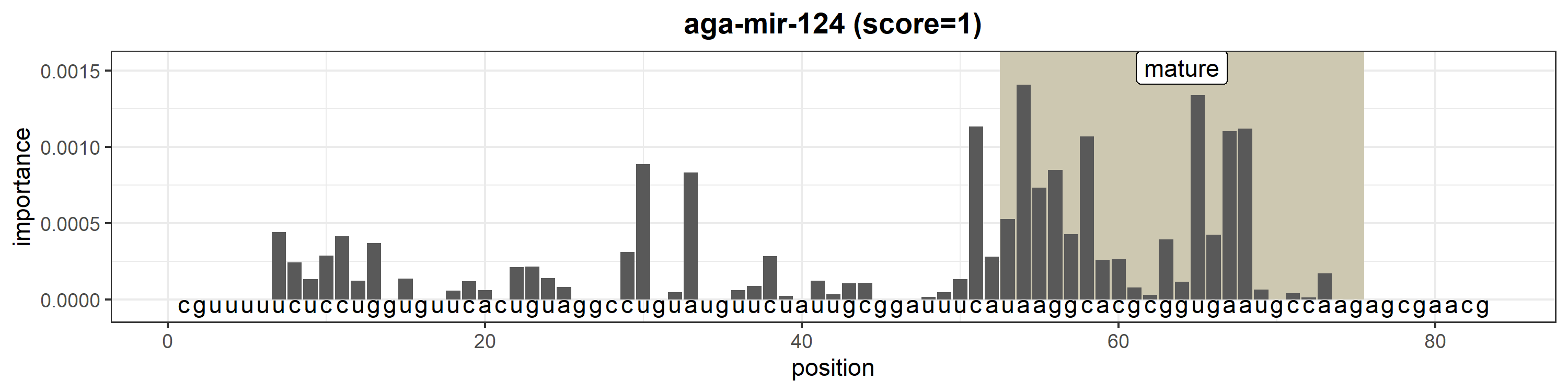

### aga-mir-125.png

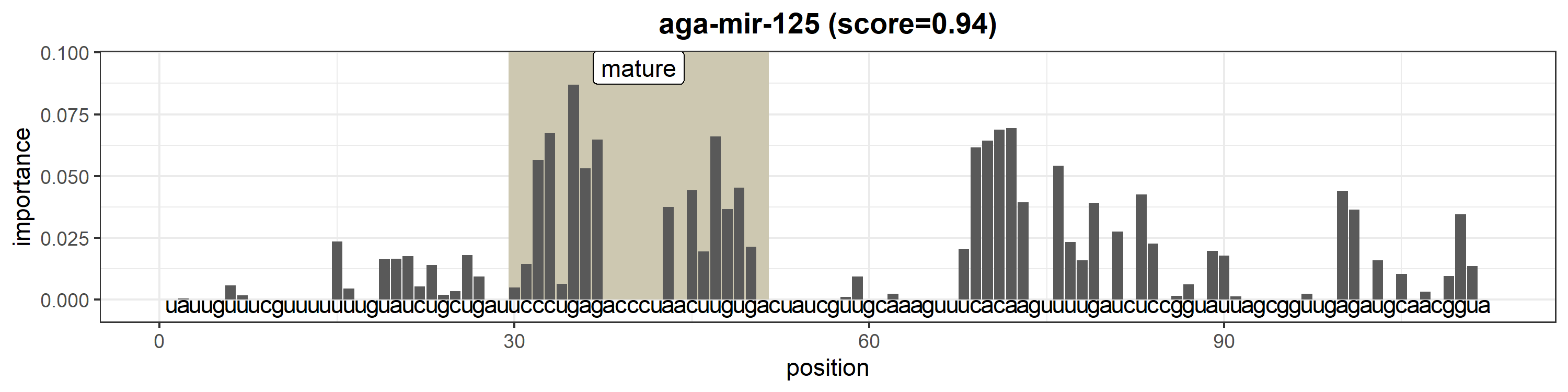

### aga-mir-133.png

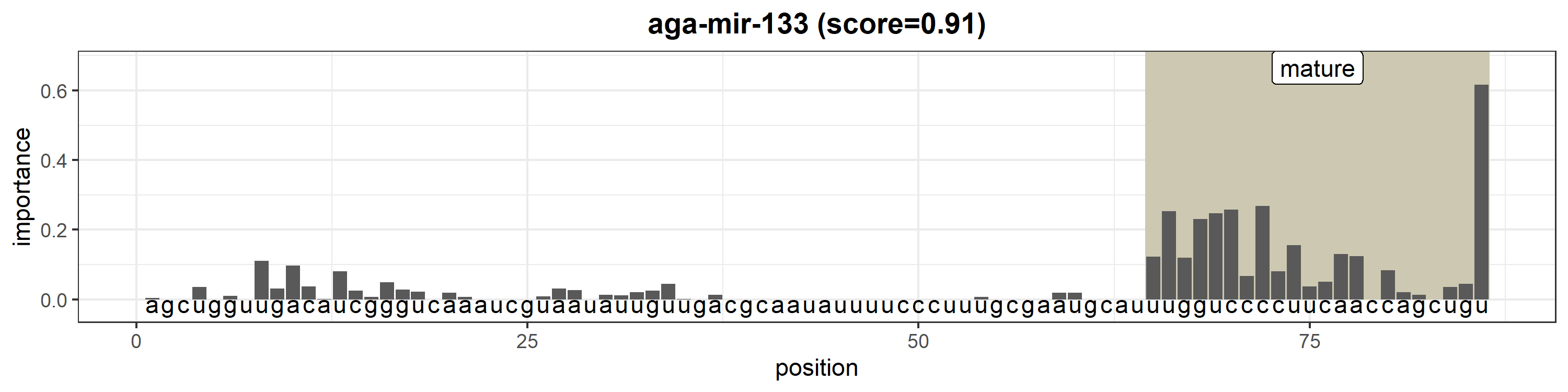

### aga-mir-137.png

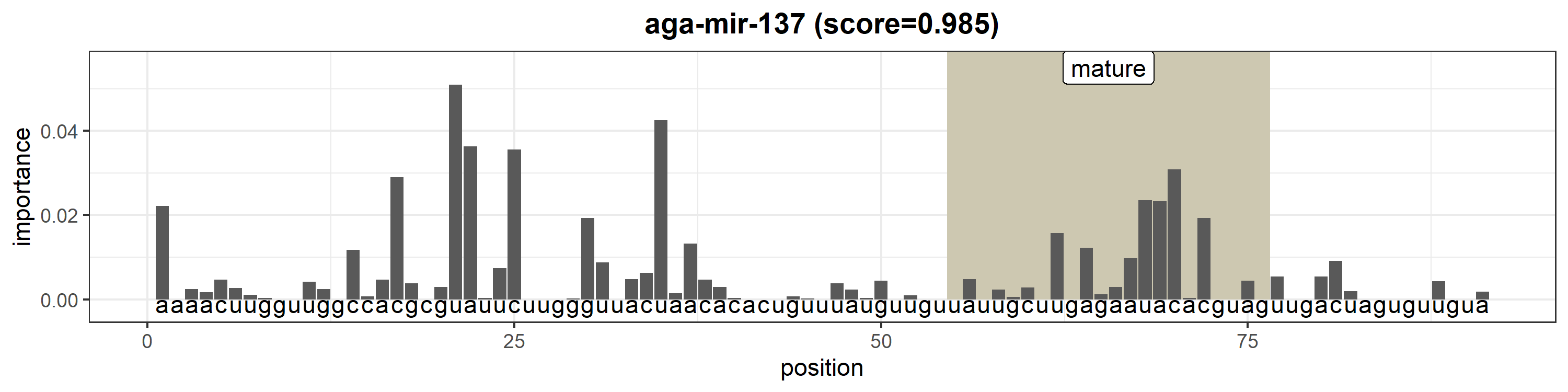

### aga-mir-184.png

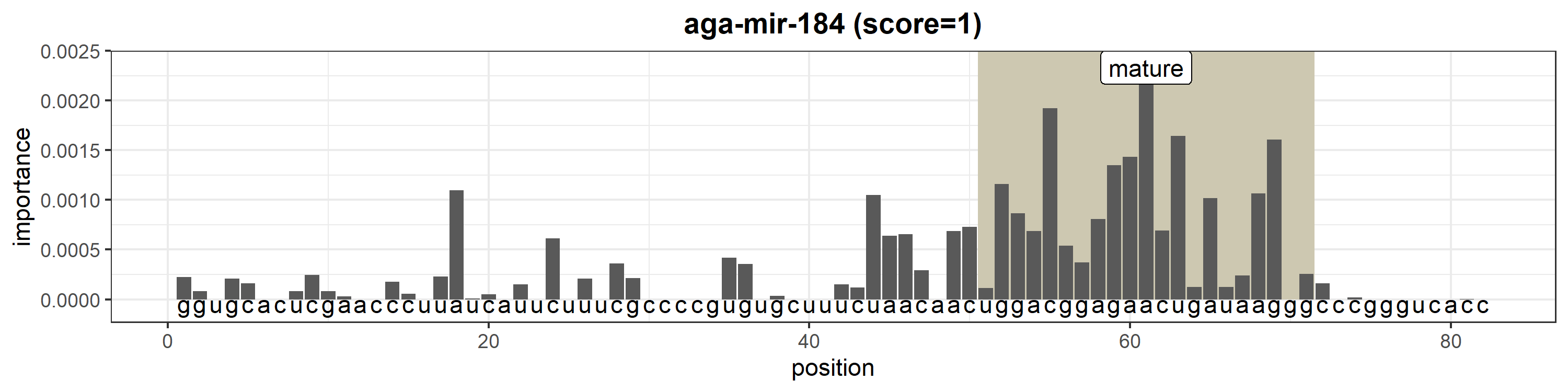

### aga-mir-190.png

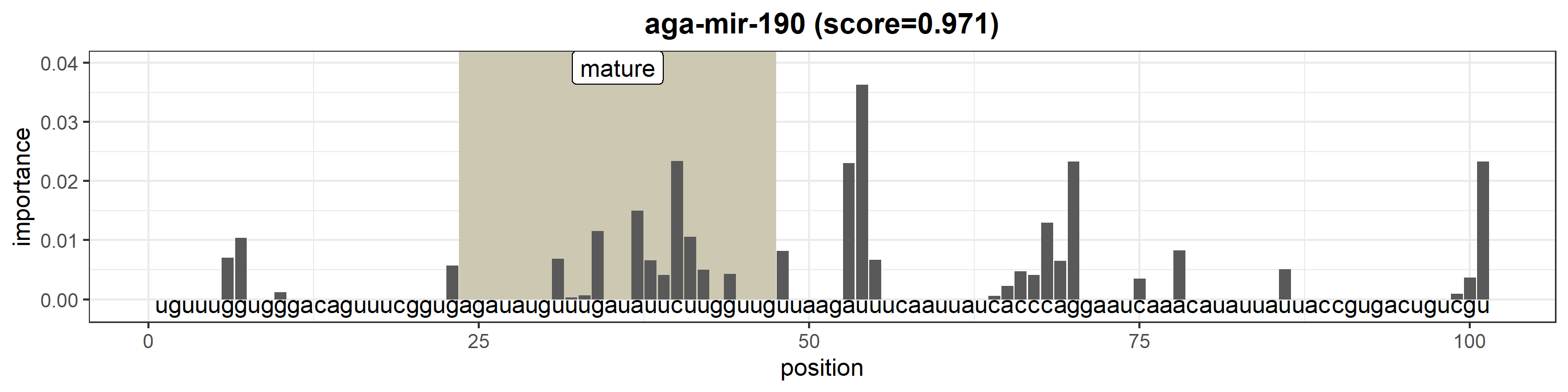

### aga-mir-210.png

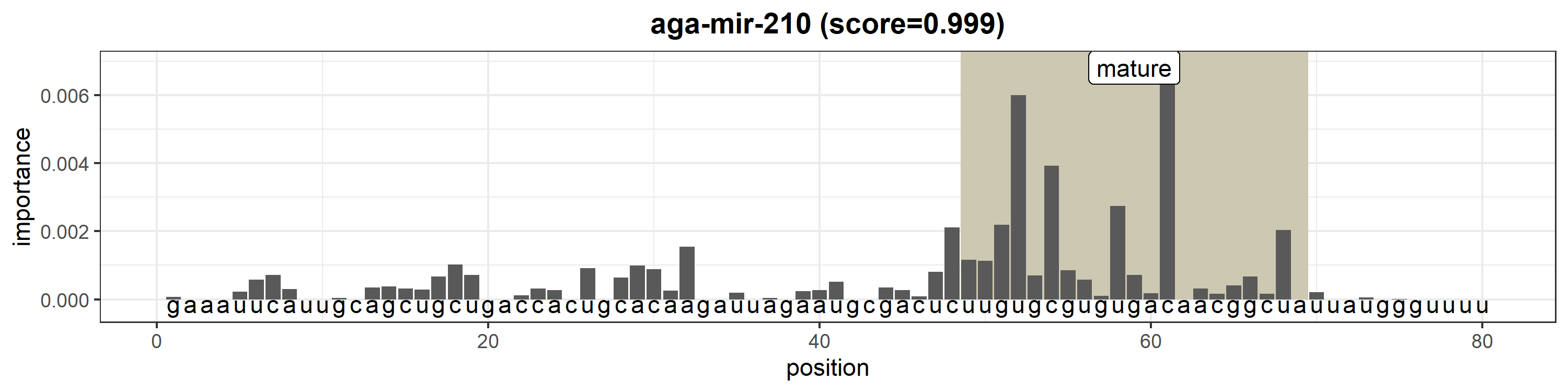

### aga-mir-219.png

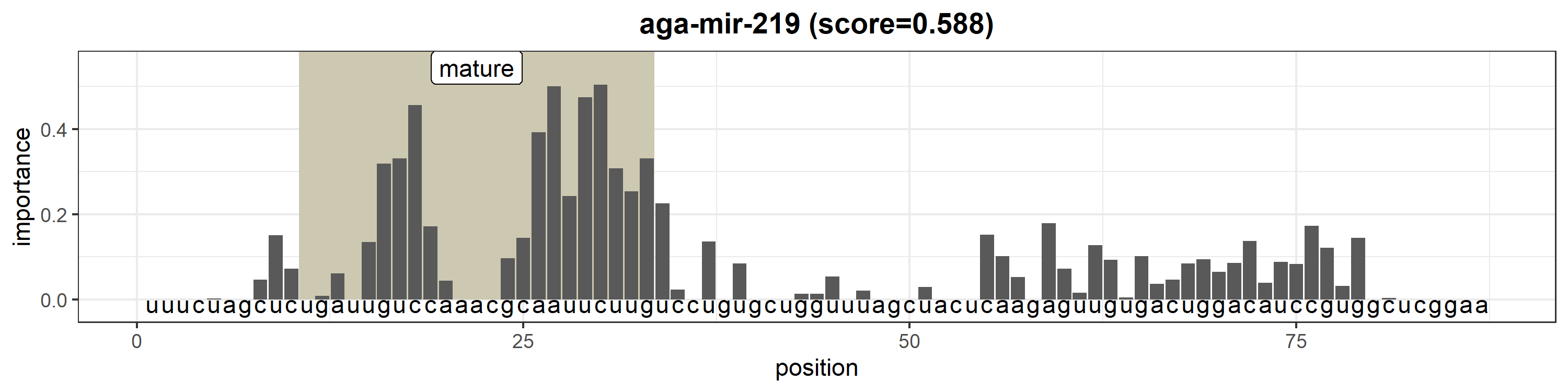

### aga-mir-263a.png

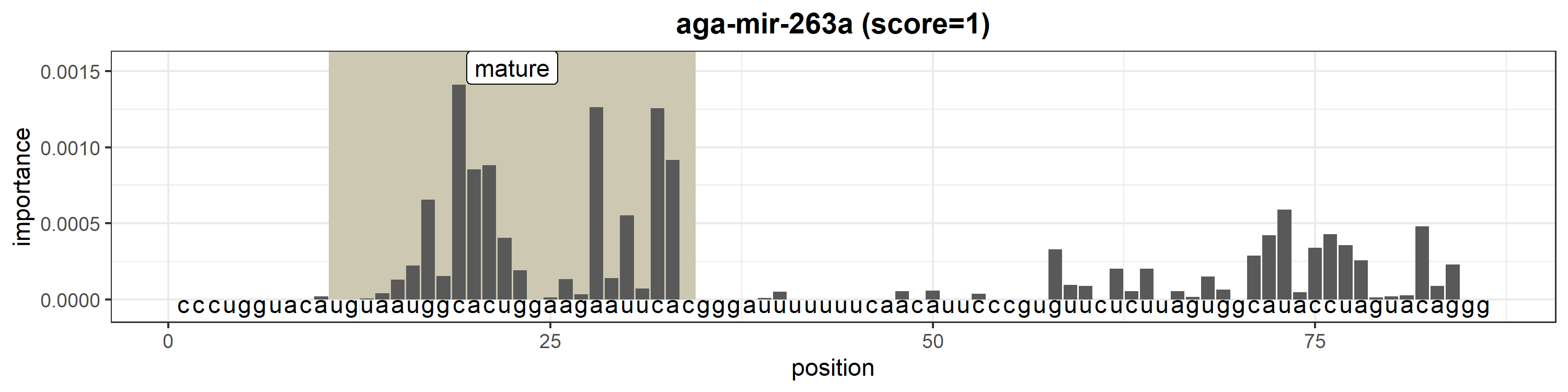

### aga-mir-263b.png

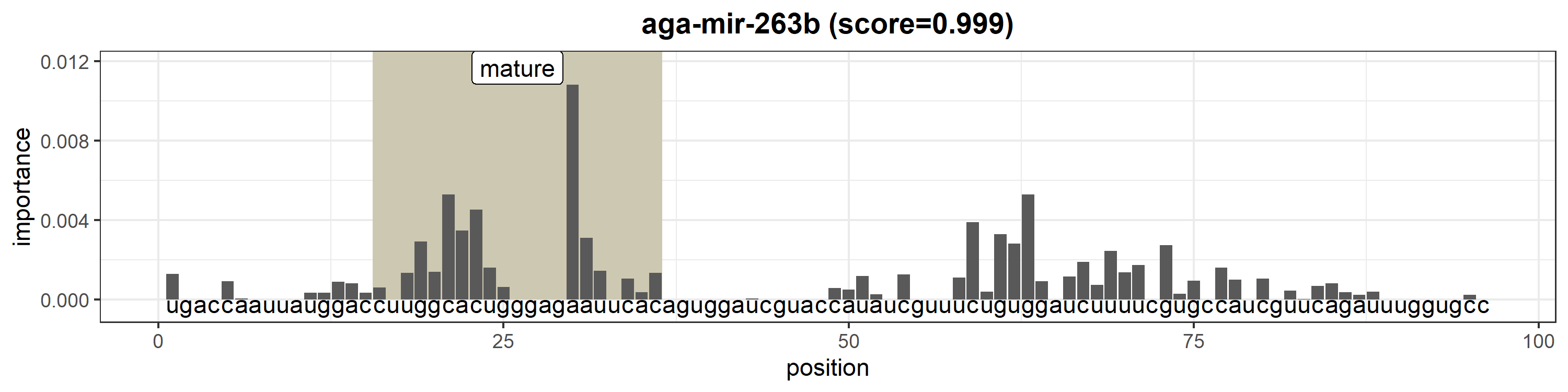

### aga-mir-275.png

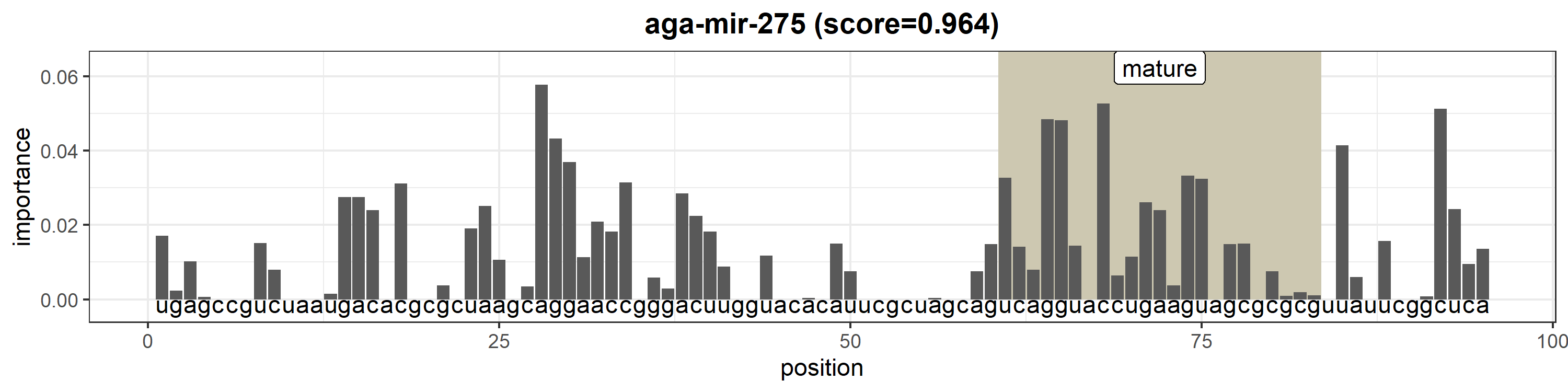

### aga-mir-276.png

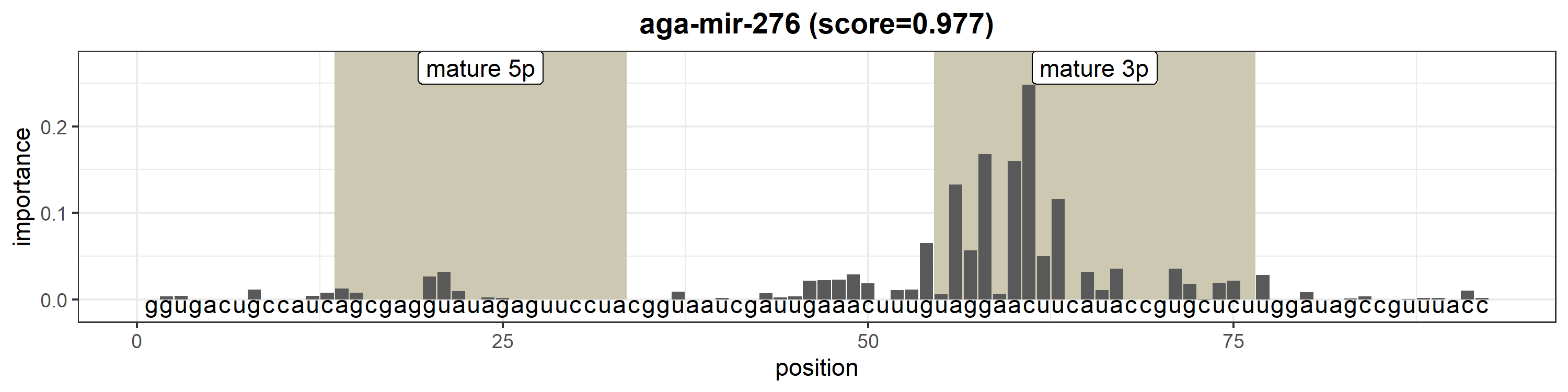

### aga-mir-277.png

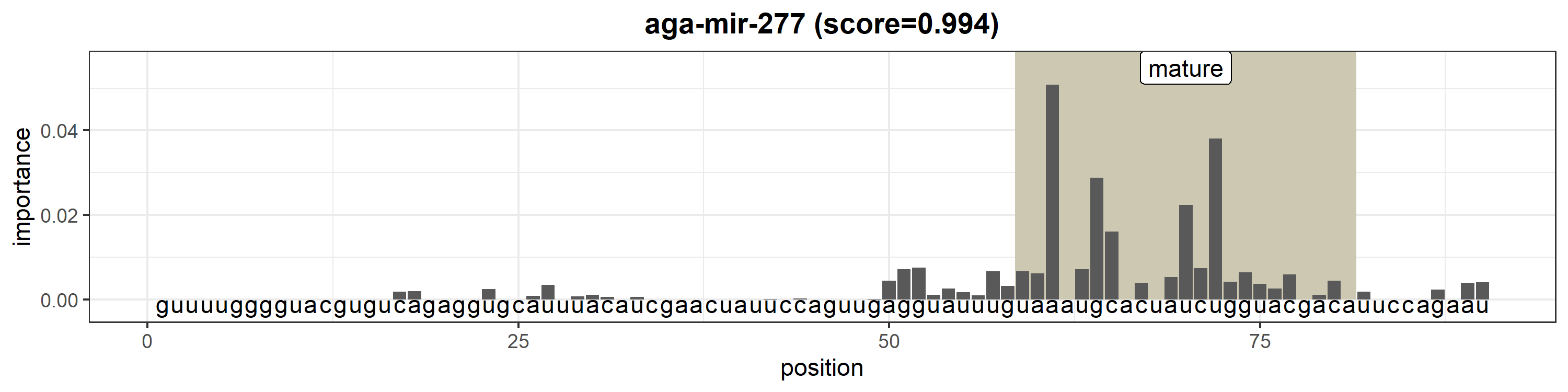

### aga-mir-278.png

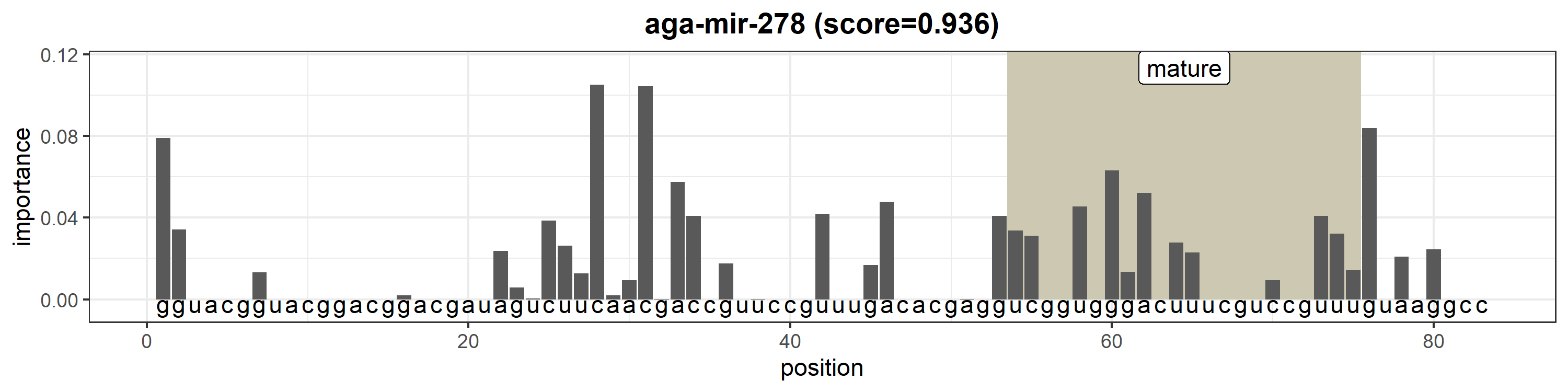

### aga-mir-279.png

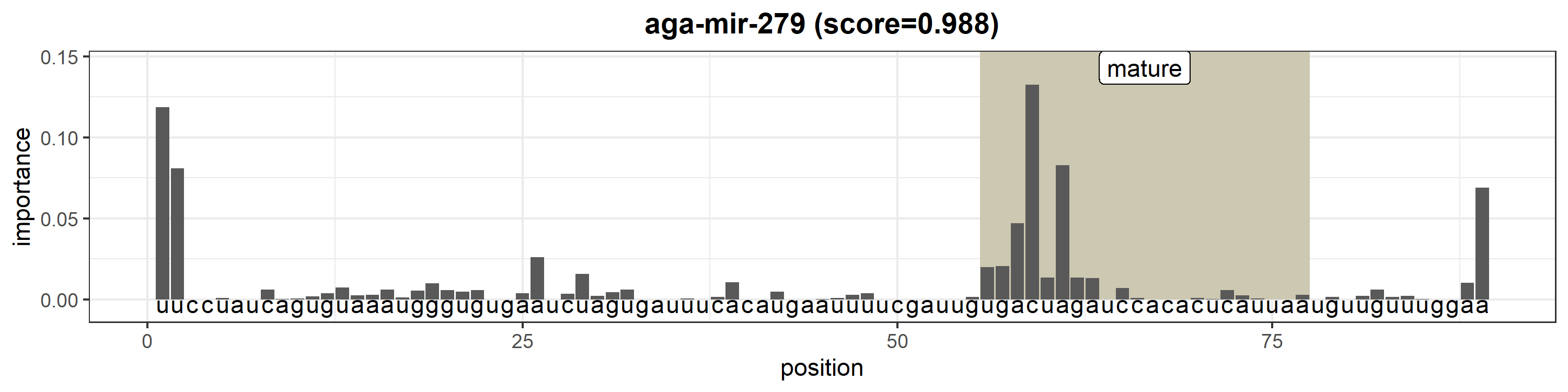

### aga-mir-281.png

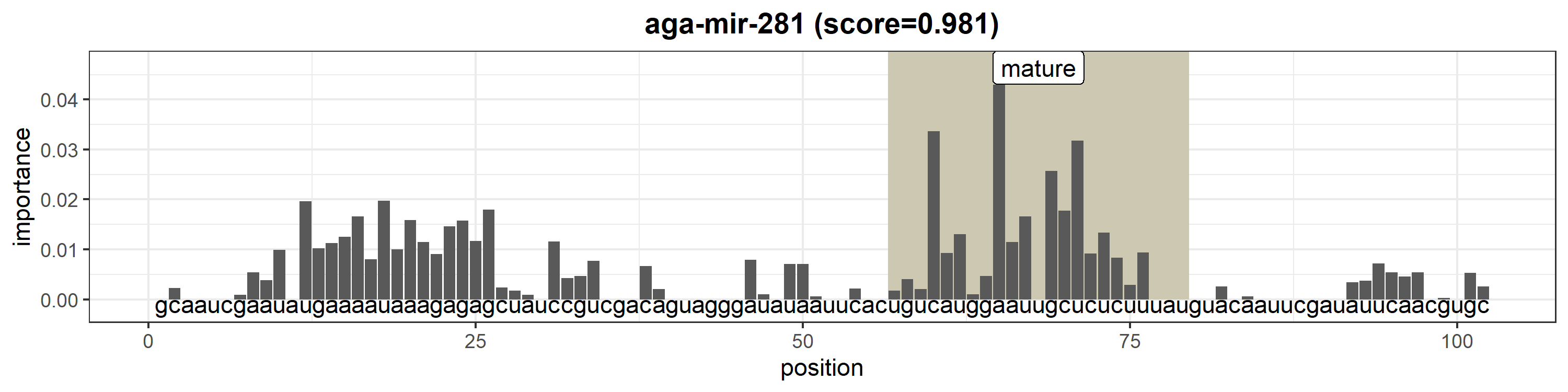

### aga-mir-282.png

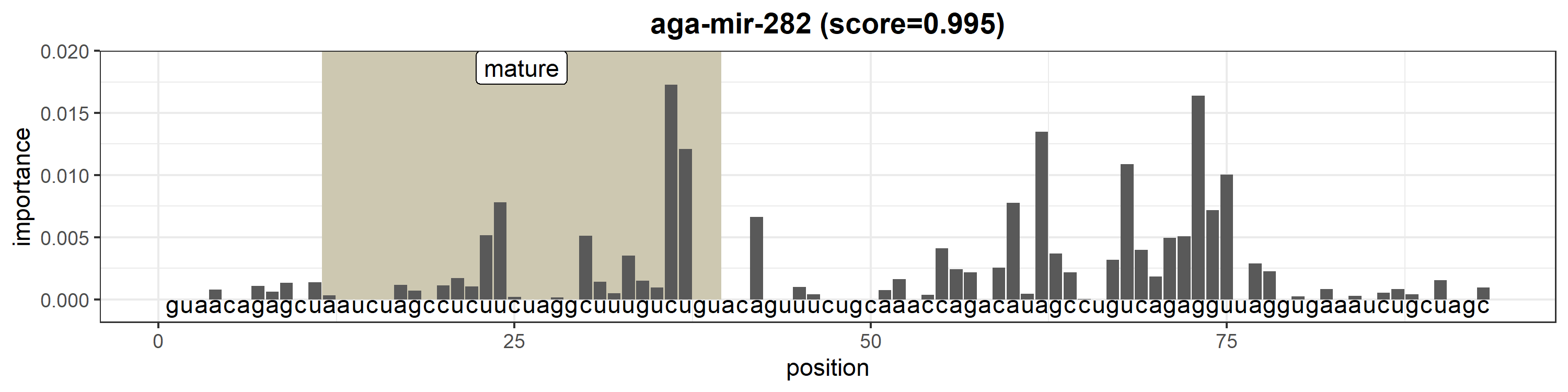

### aga-mir-283.png

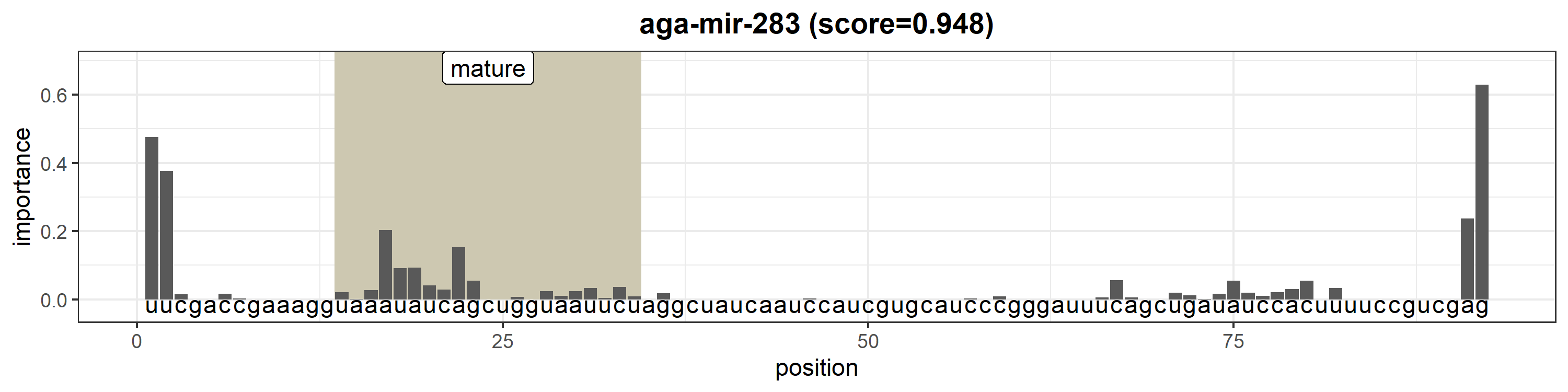

### aga-mir-285.png

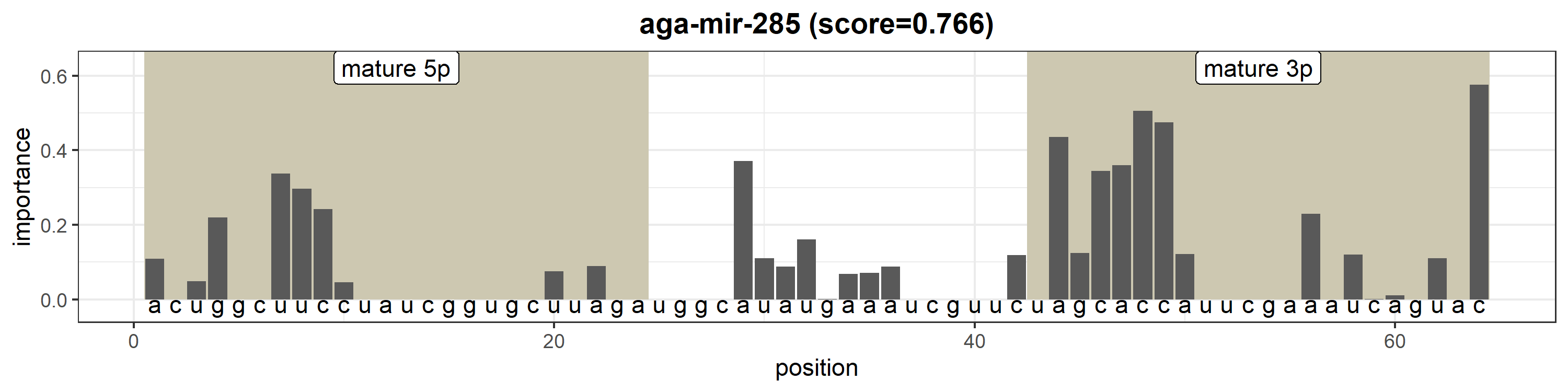

### aga-mir-286.png

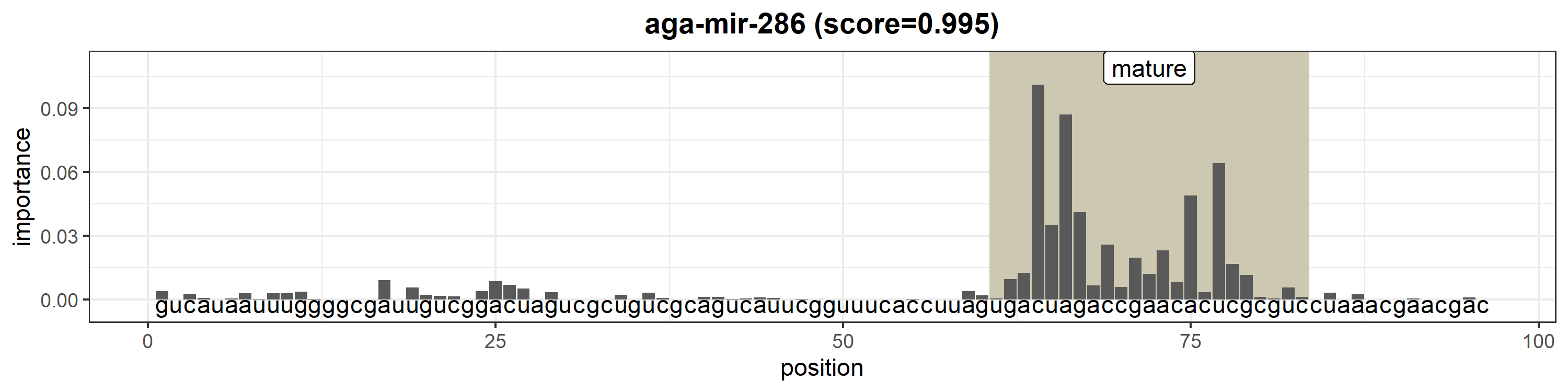

### aga-mir-286b.png

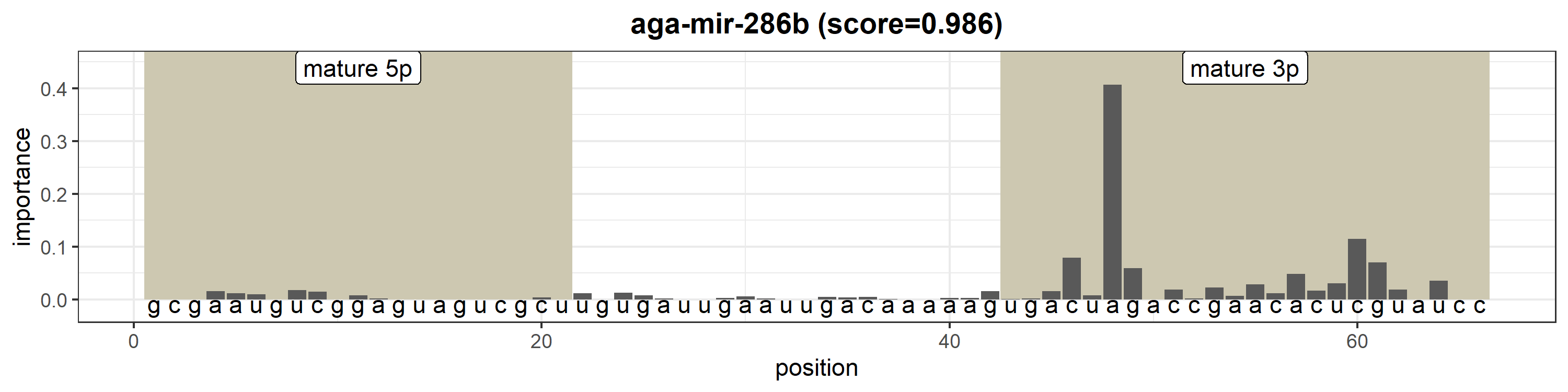

### aga-mir-305.png

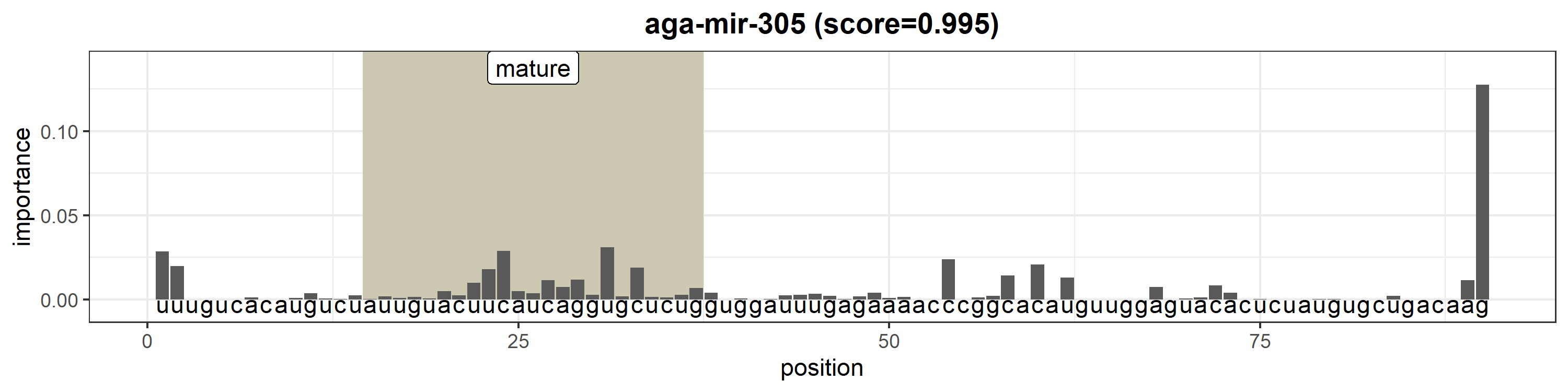

### aga-mir-306.png

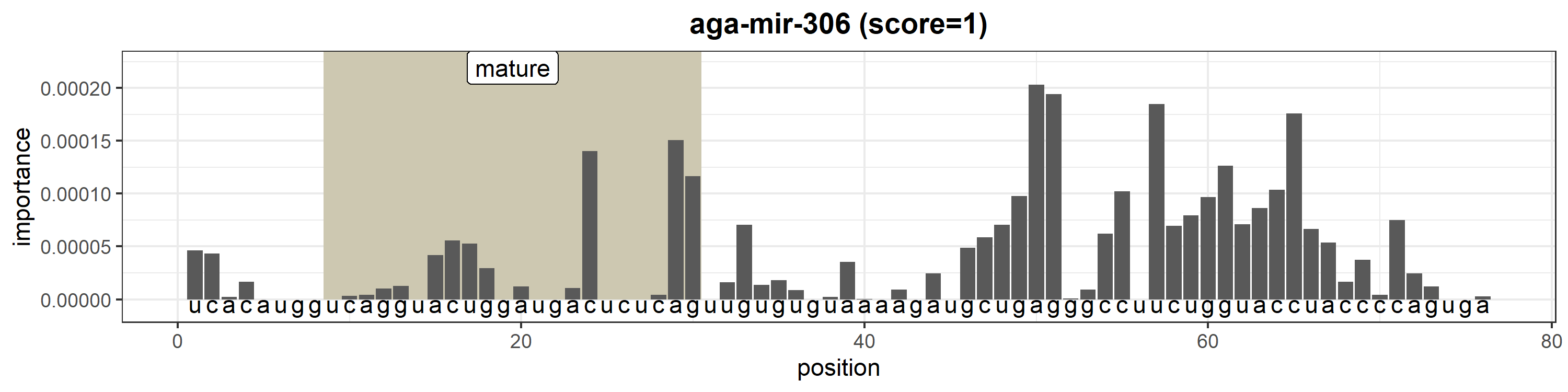

### aga-mir-307.png

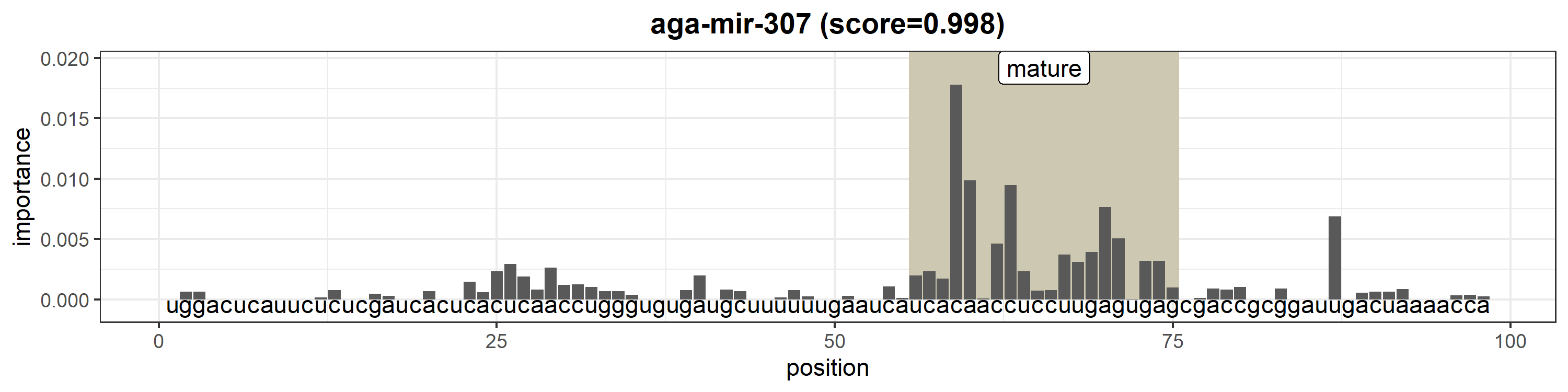

### aga-mir-308.png

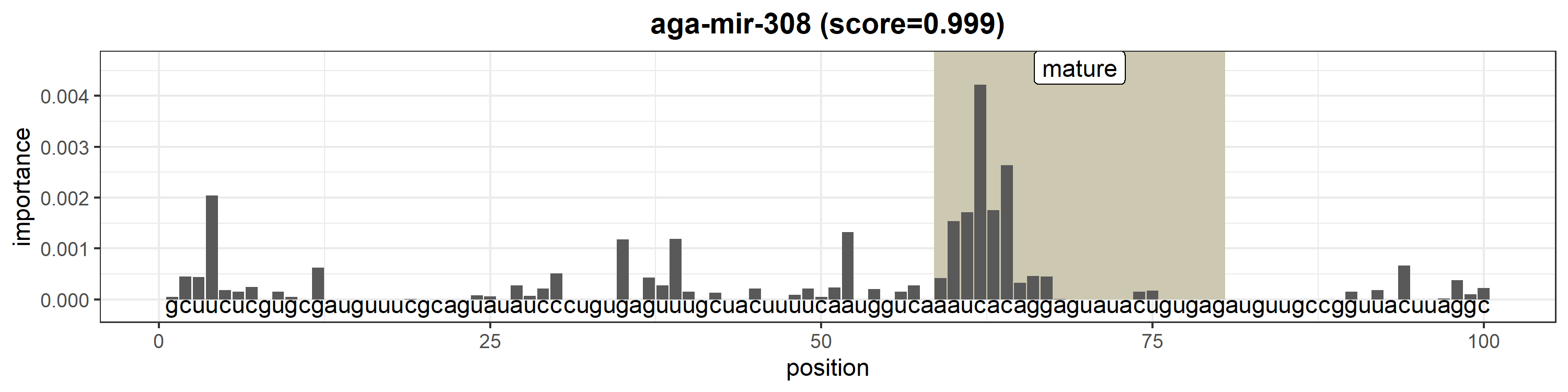

### aga-mir-309.png

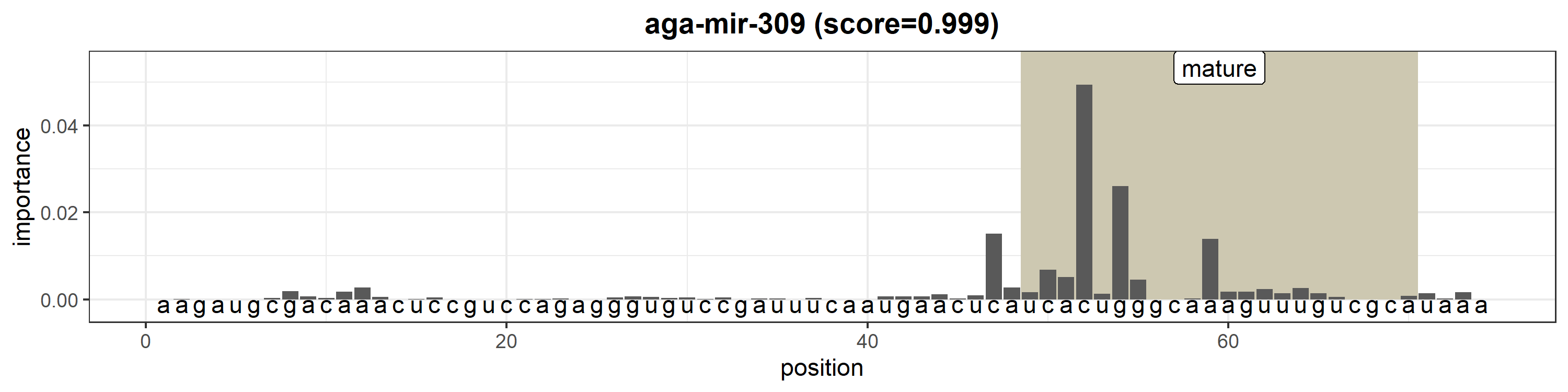

### aga-mir-1000.png

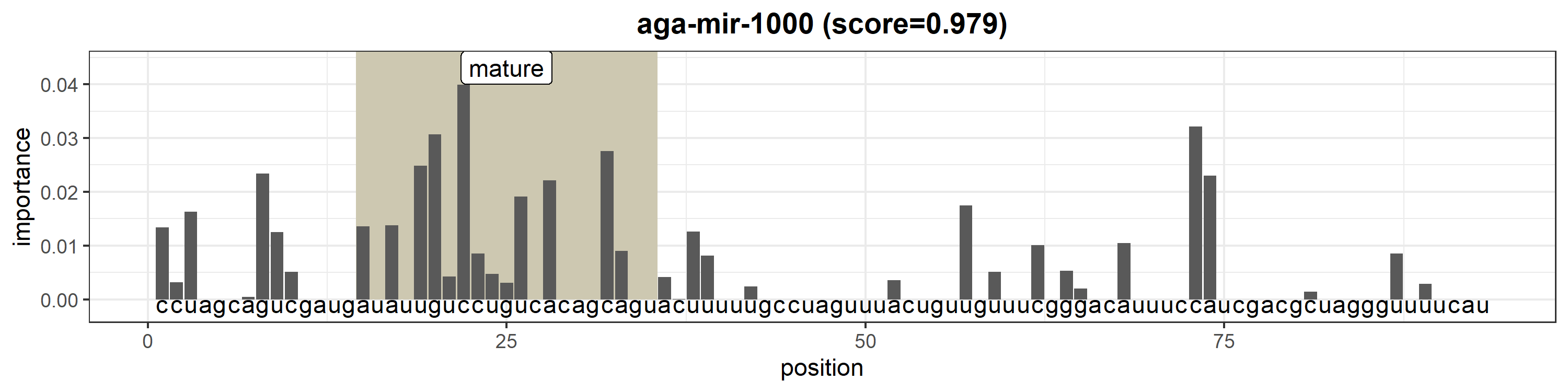

### aga-mir-1175.png

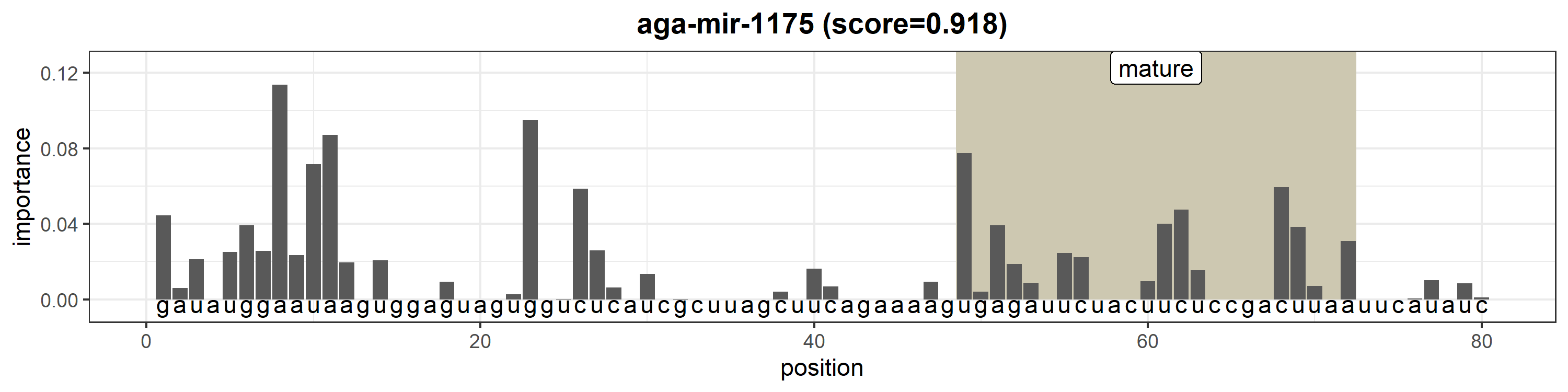

### aga-mir-1889.png

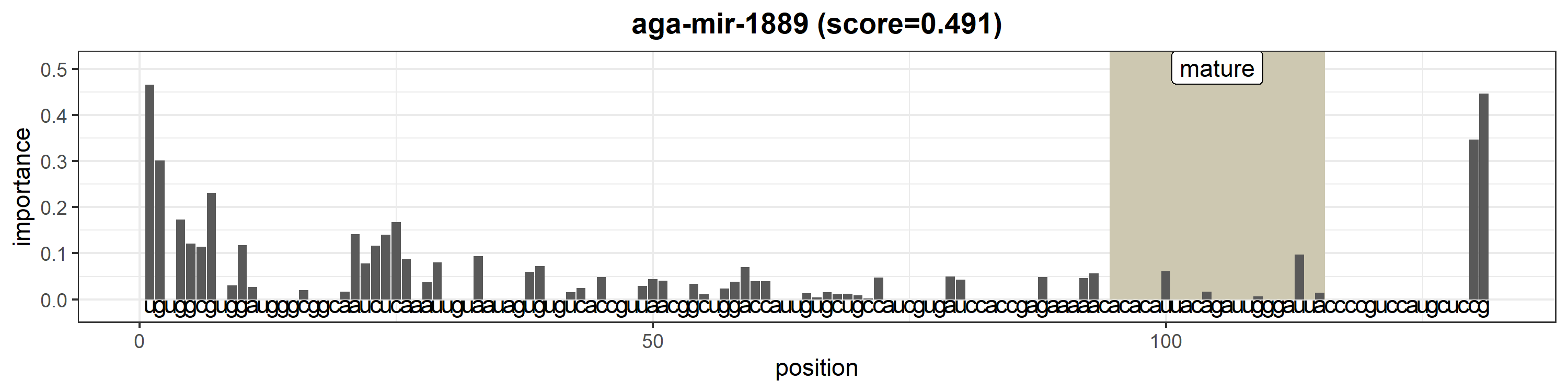

### aga-mir-1890.png

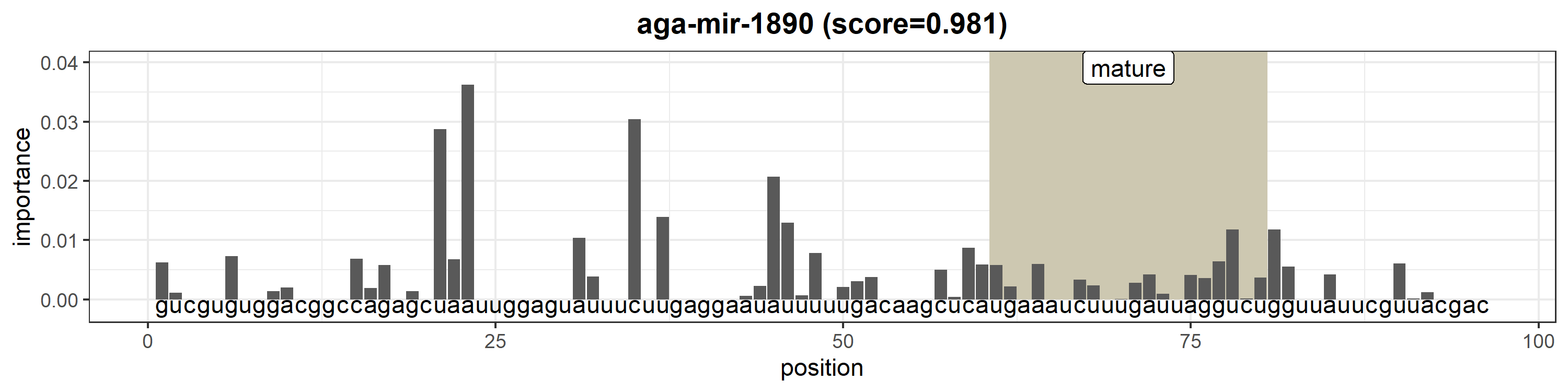

### aga-mir-1891.png

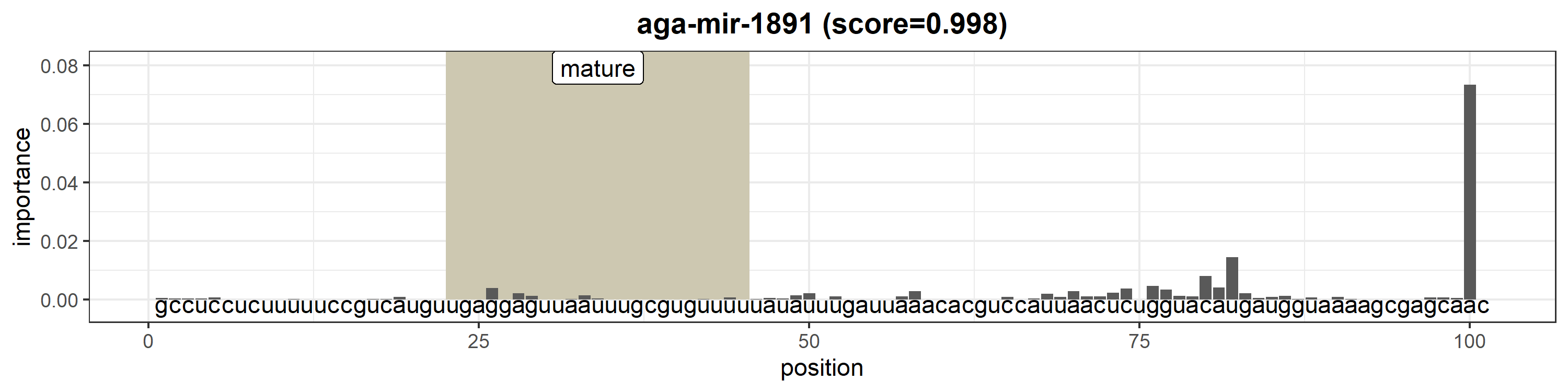

### aga-mir-2796.png

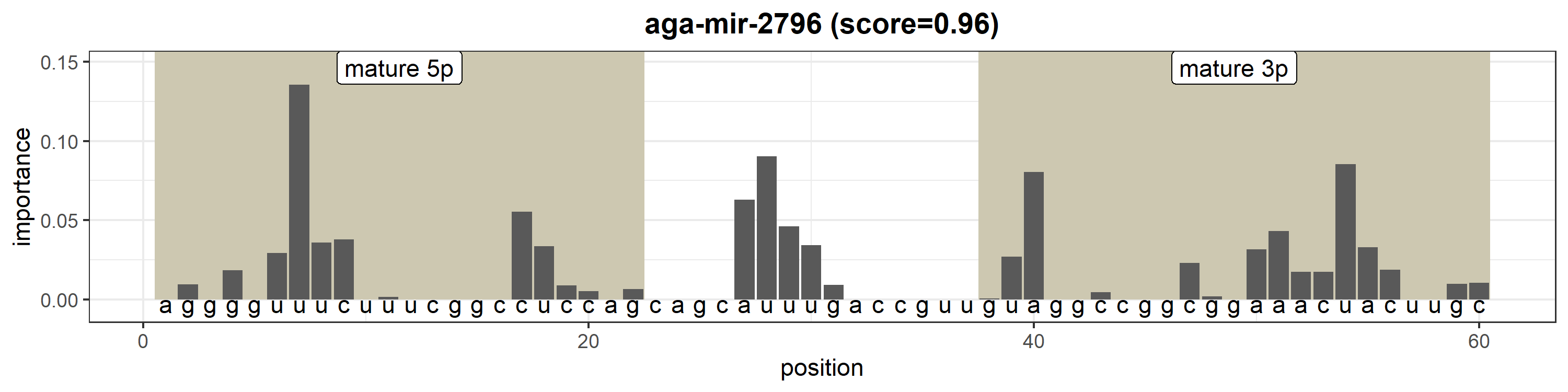

### aga-mir-2944a-1.png

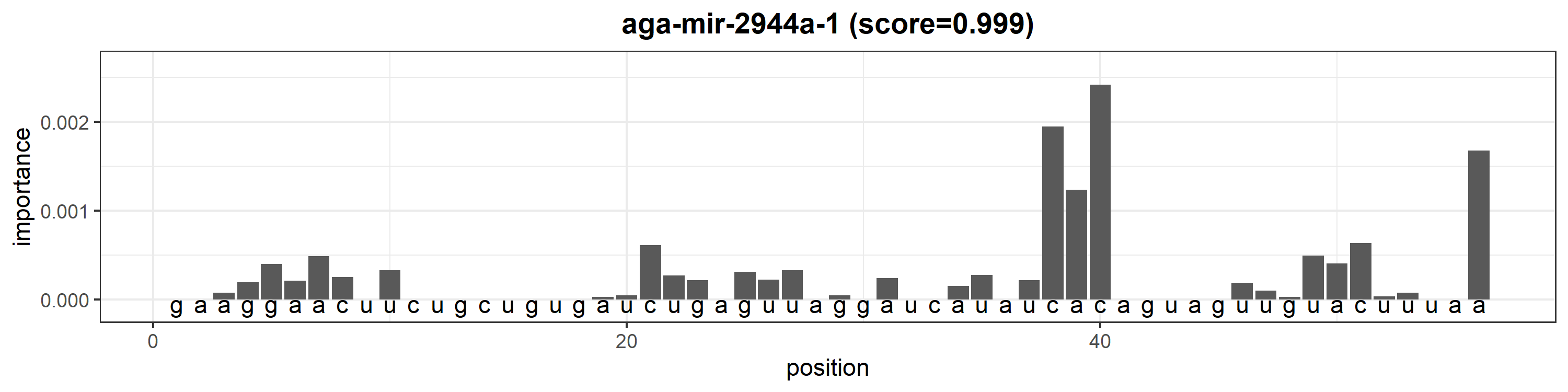

### aga-mir-2944a-2.png

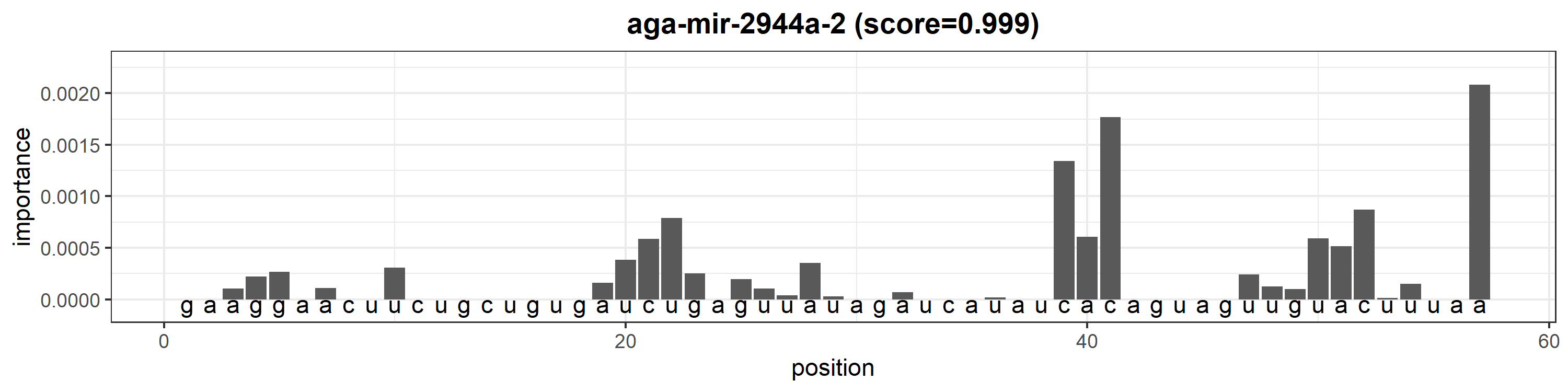

### aga-mir-2944b.png

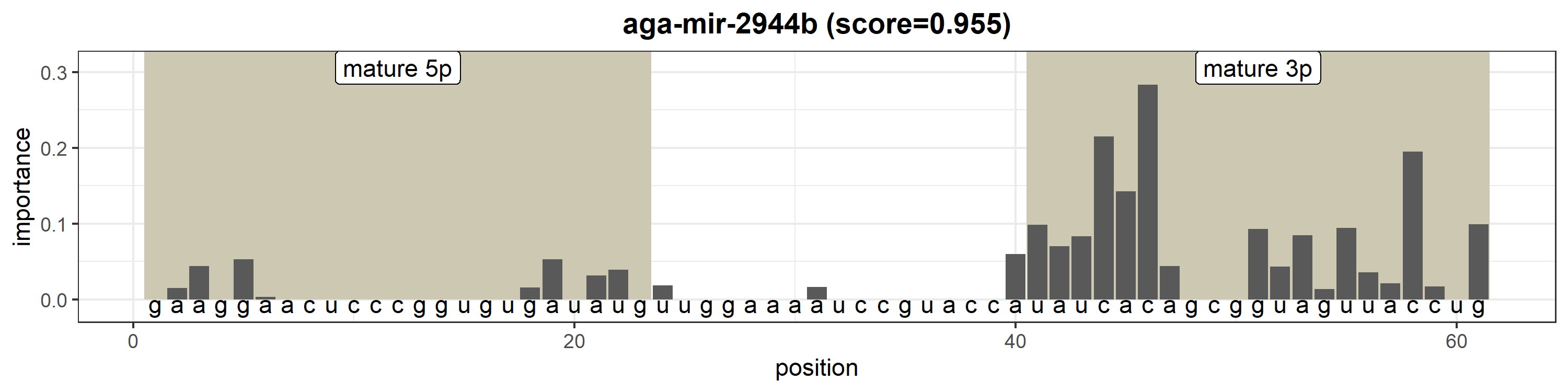

### aga-mir-2945.png

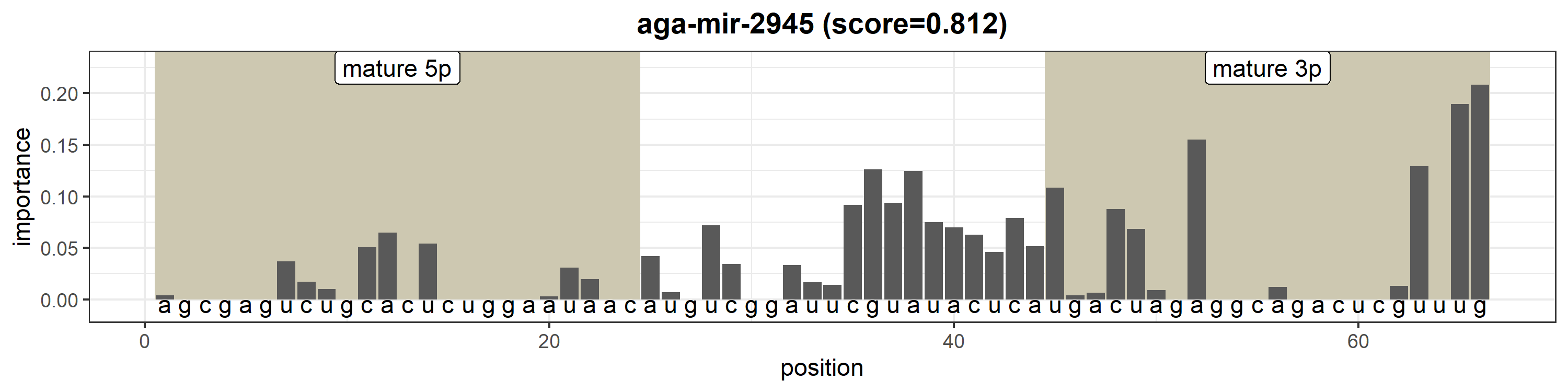

### aga-mir-10355.png

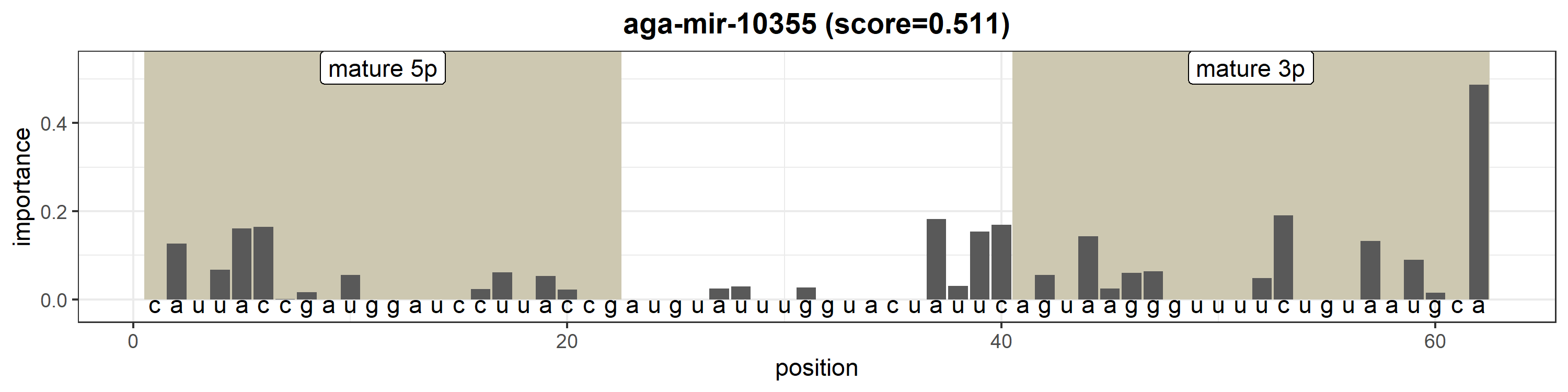

### aga-mir-10356-1.png

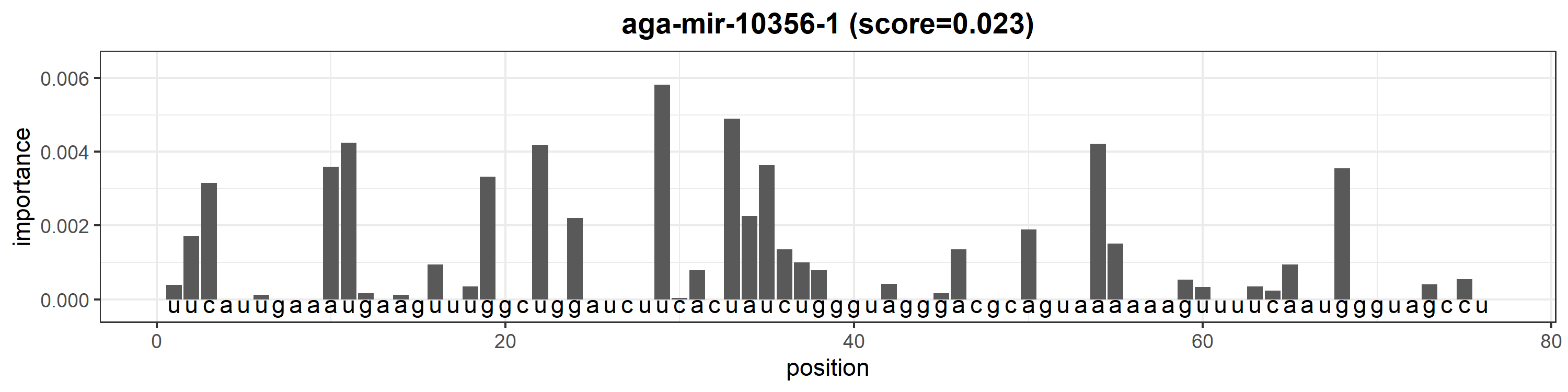

### aga-mir-10356-2.png

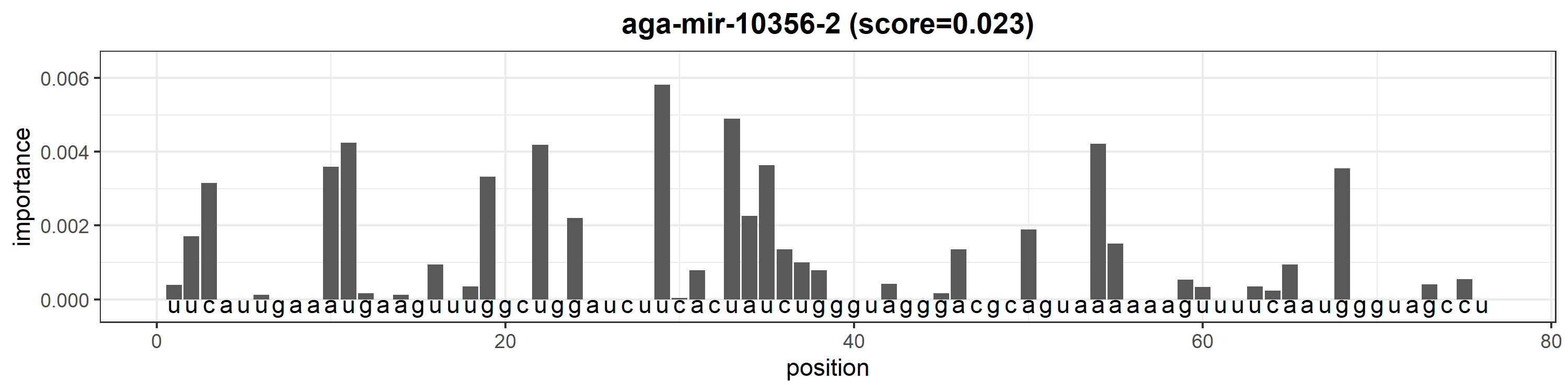

### aga-mir-10357.png

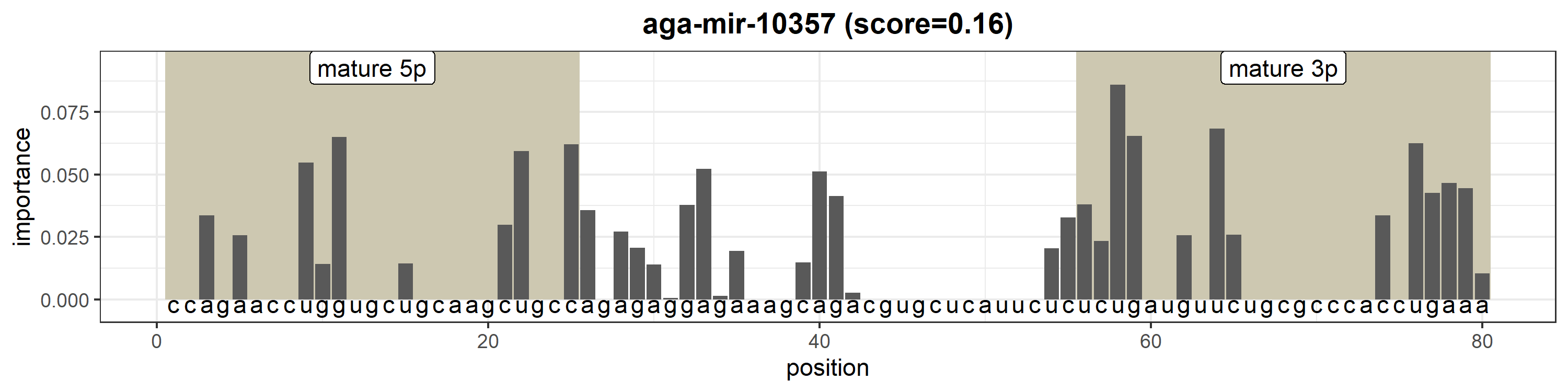

### aga-mir-10358-1.png

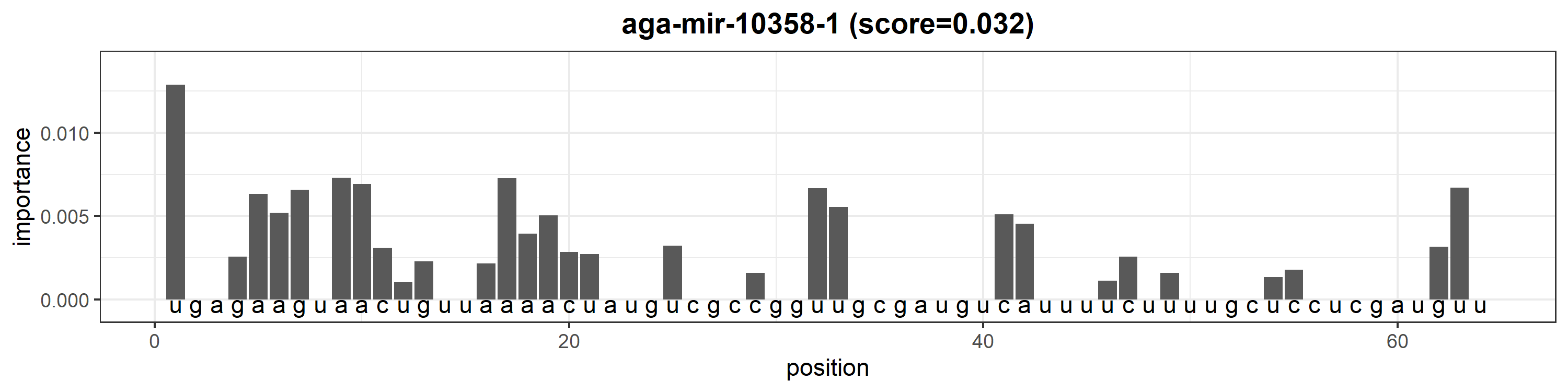

### aga-mir-10358-2.png

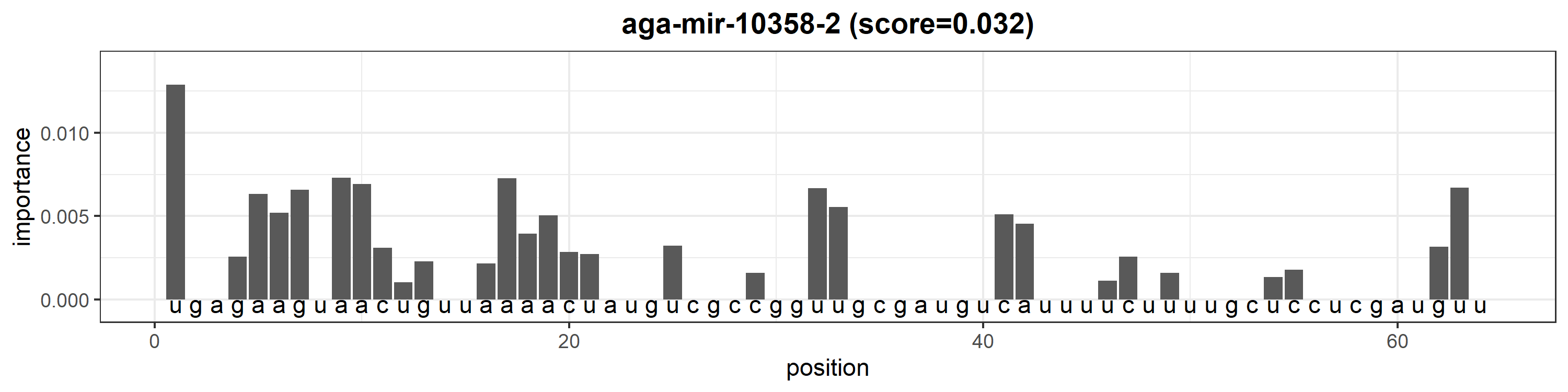

### aga-mir-10358-3.png

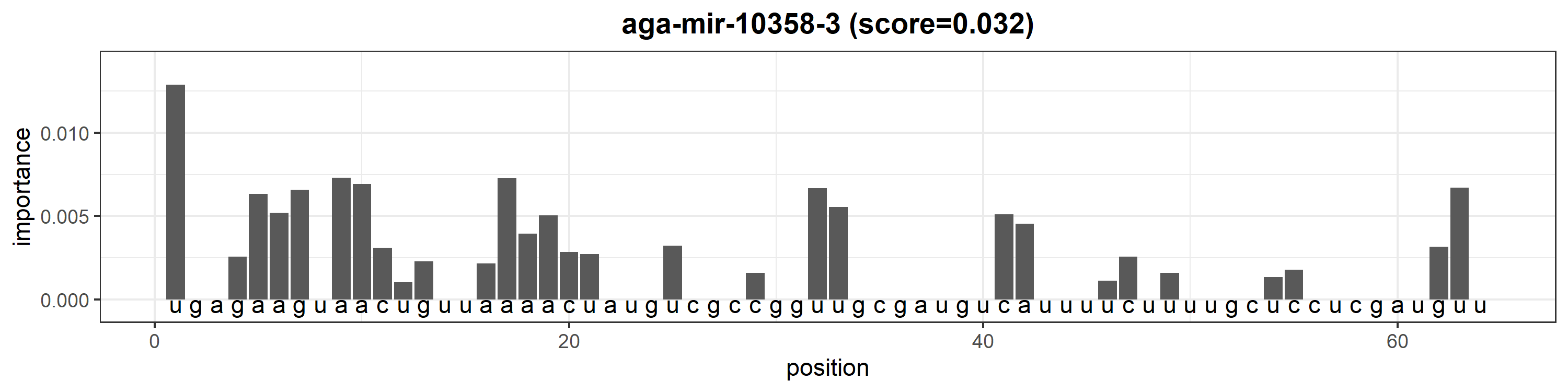

### aga-mir-10358-4.png

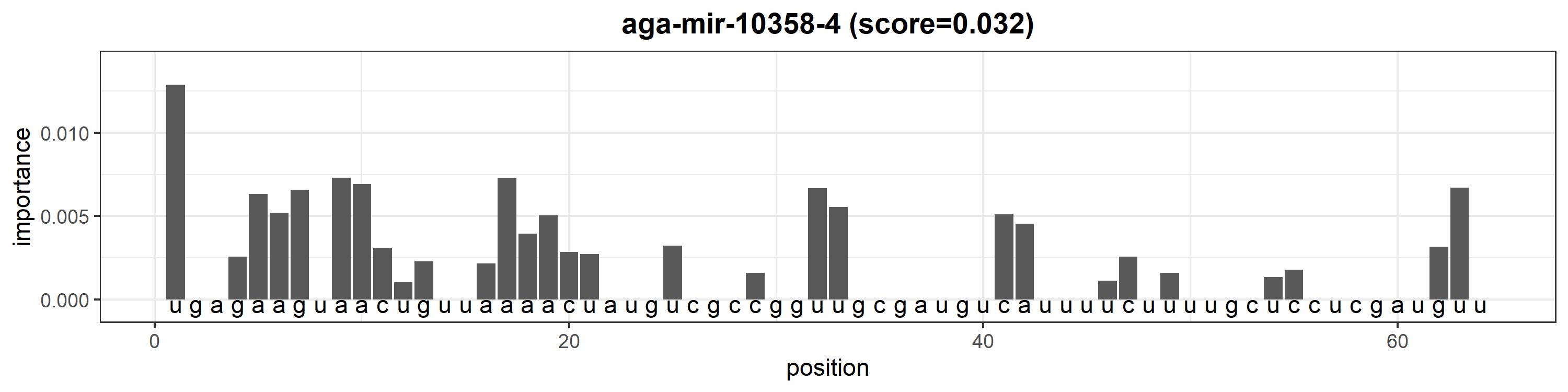

### aga-mir-10358-5.png

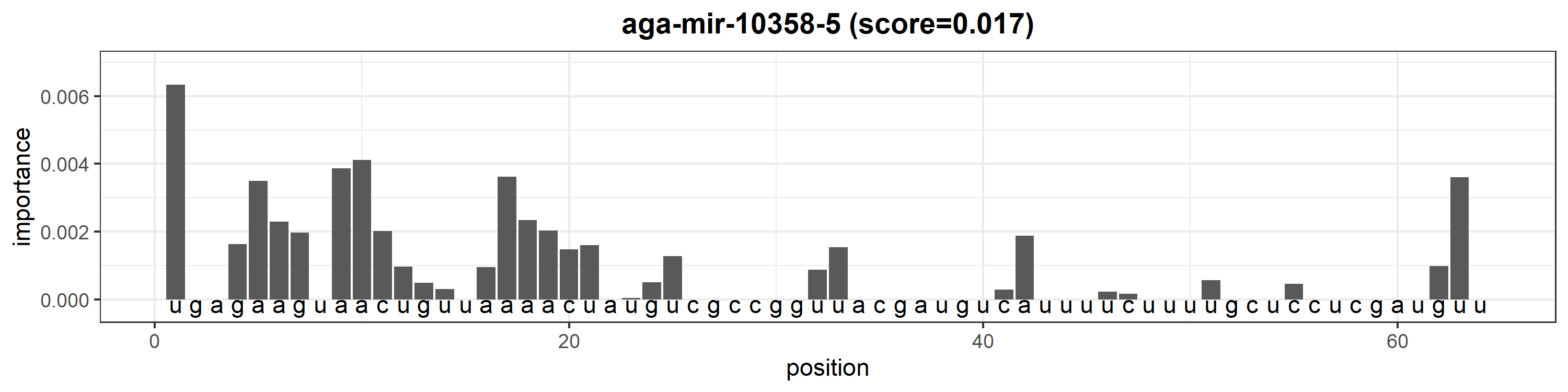

### aga-mir-10359.png

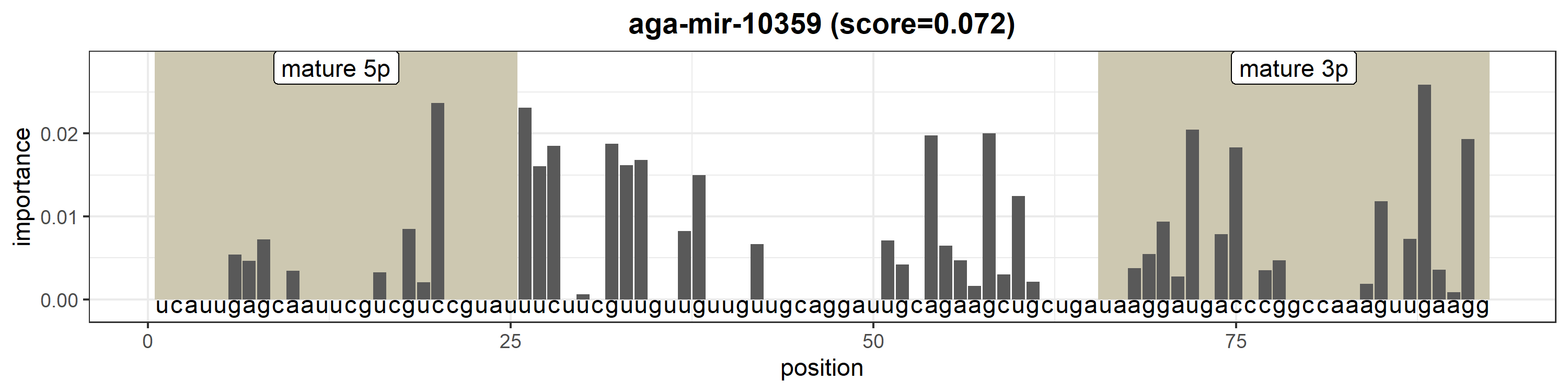

### aga-mir-10360.png

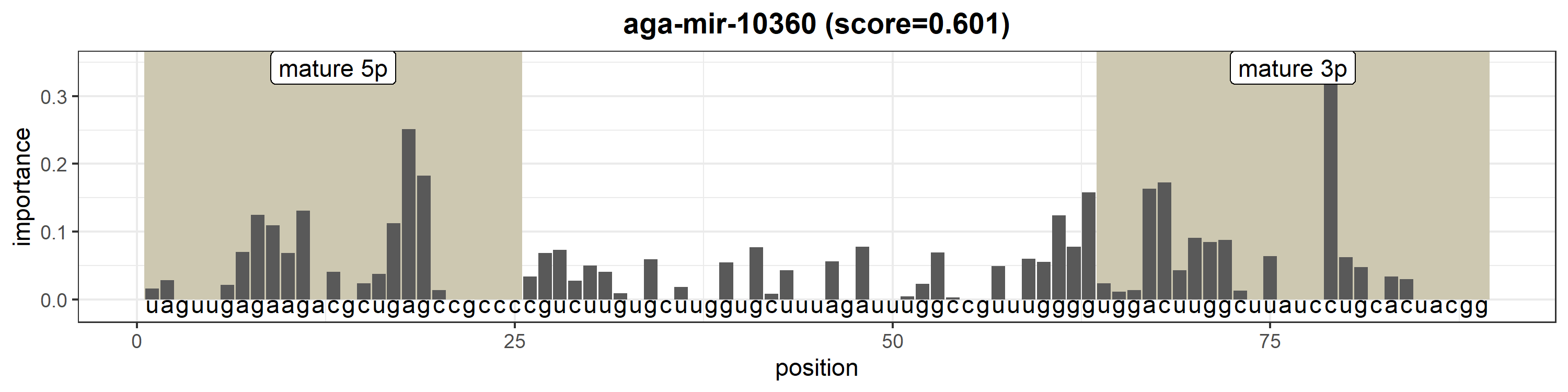

### aga-mir-10361a-1.png

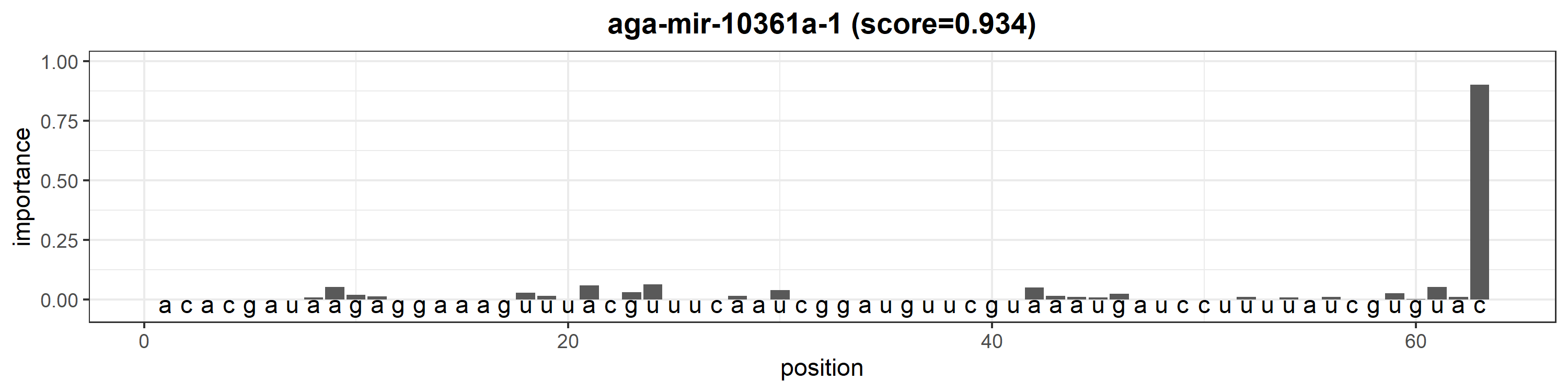

### aga-mir-10361a-2.png

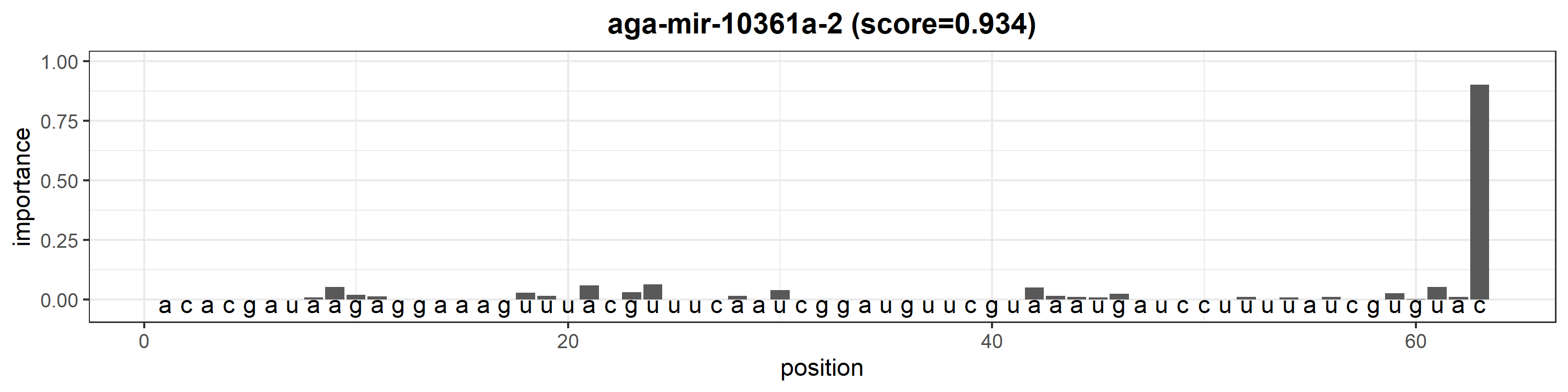

### aga-mir-10361a-3.png

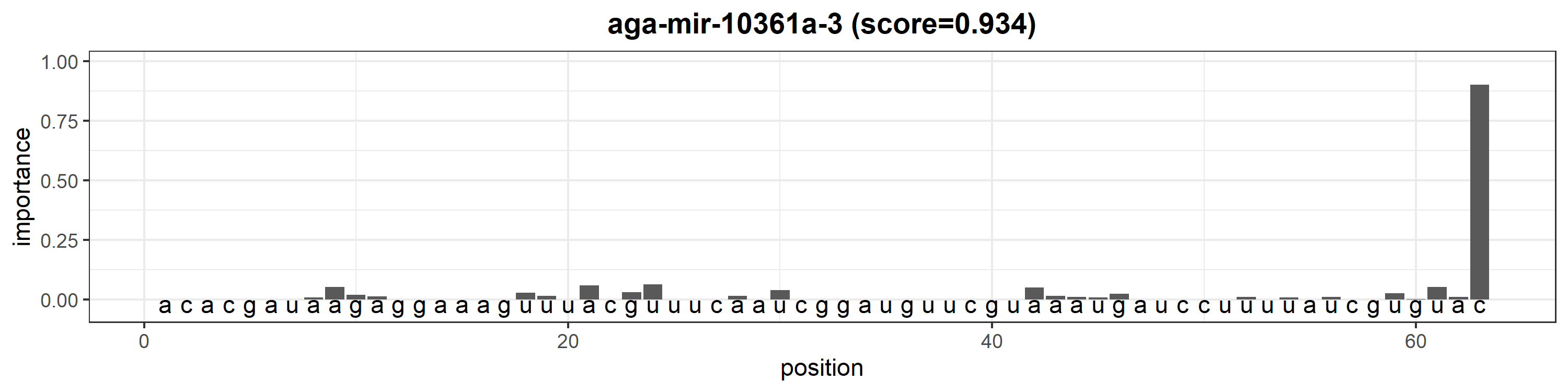

### aga-mir-10361b.png

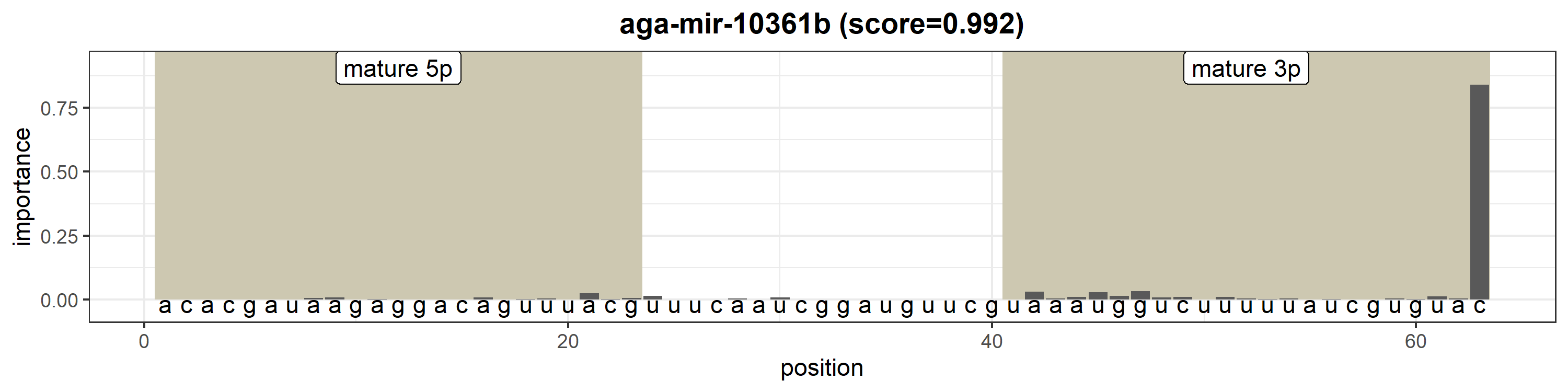

### aga-mir-10362.png

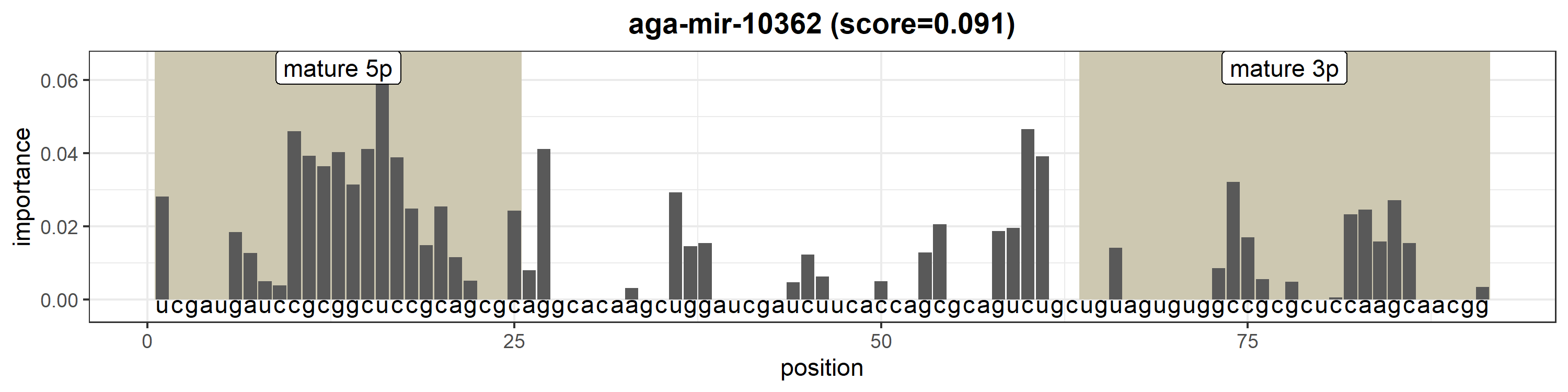

### aga-mir-10363.png

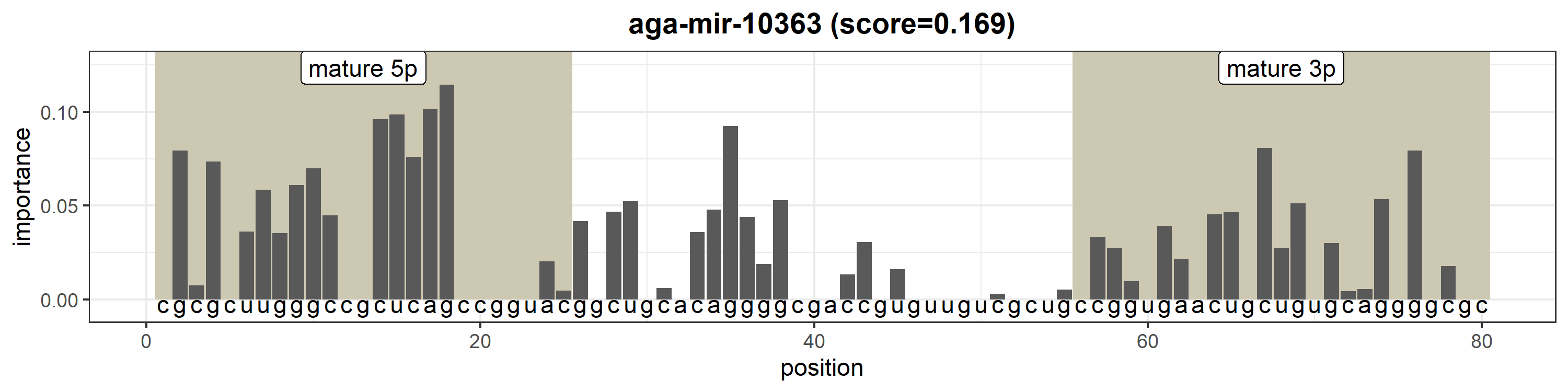

### aga-mir-10364.png

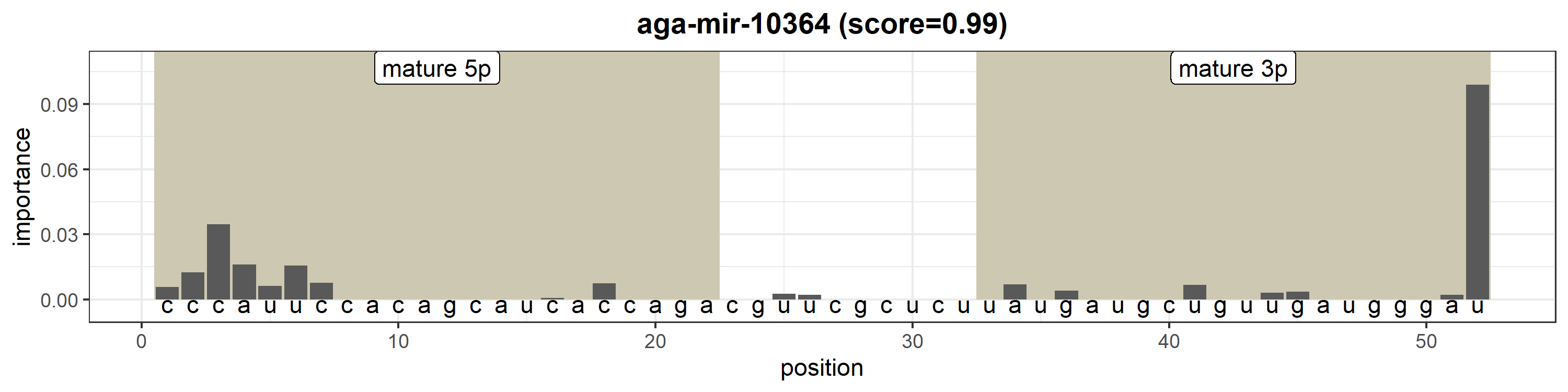

### aga-mir-10365.png

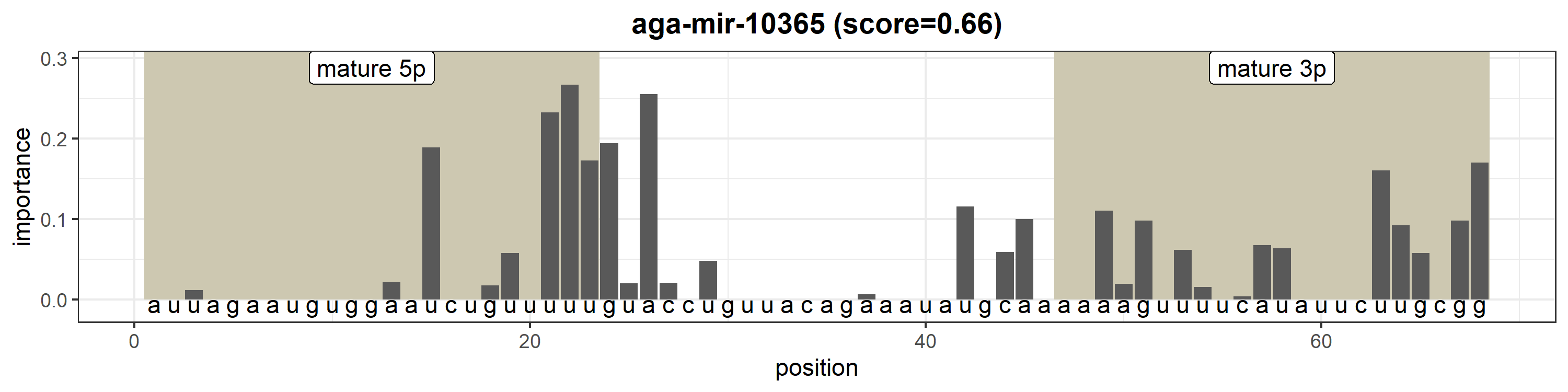

### aga-mir-10366a-1.png

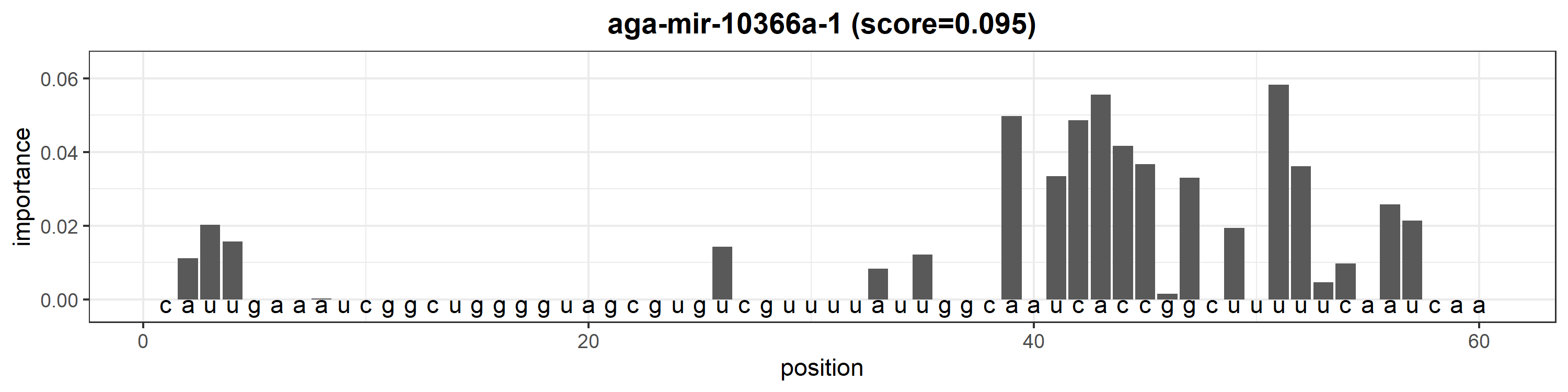

### aga-mir-10366a-2.png

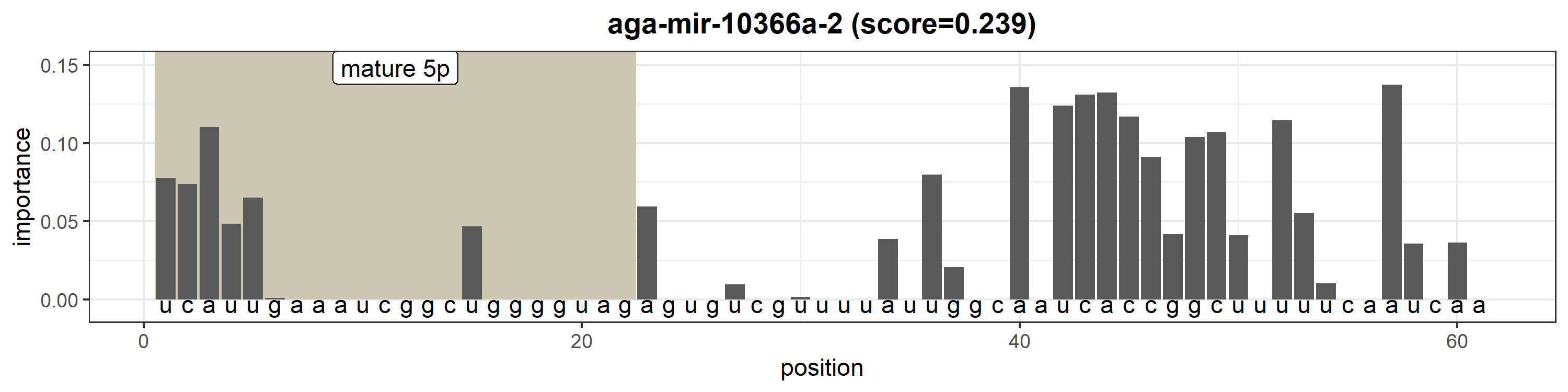

### aga-mir-10366b.png

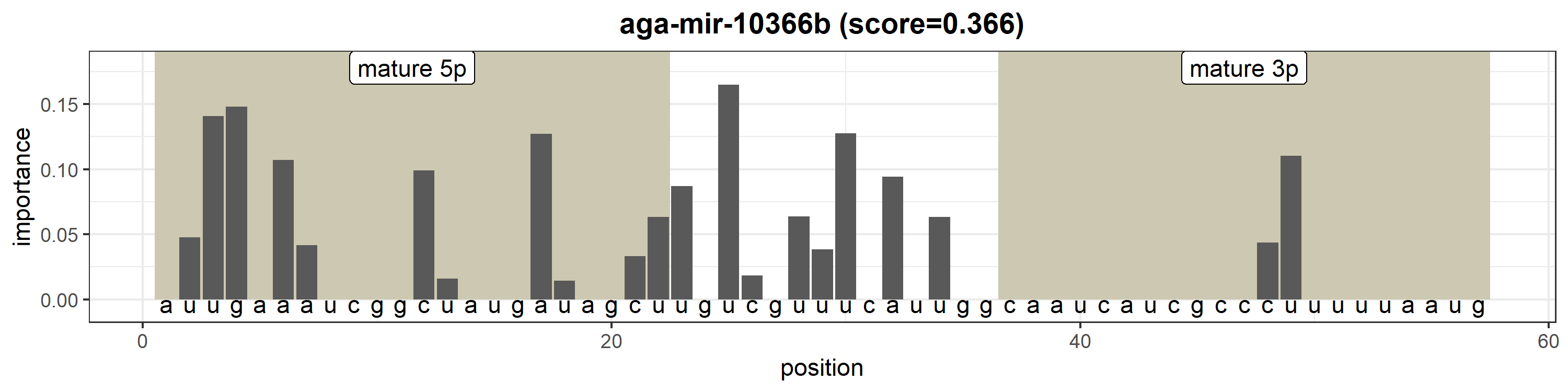

### aga-mir-10367.png

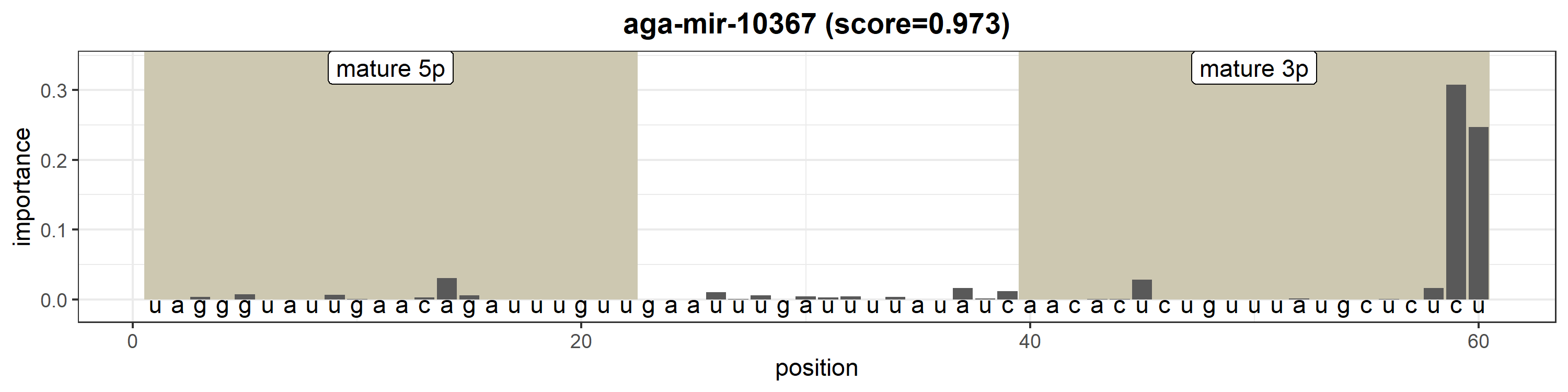

### aga-mir-10368.png

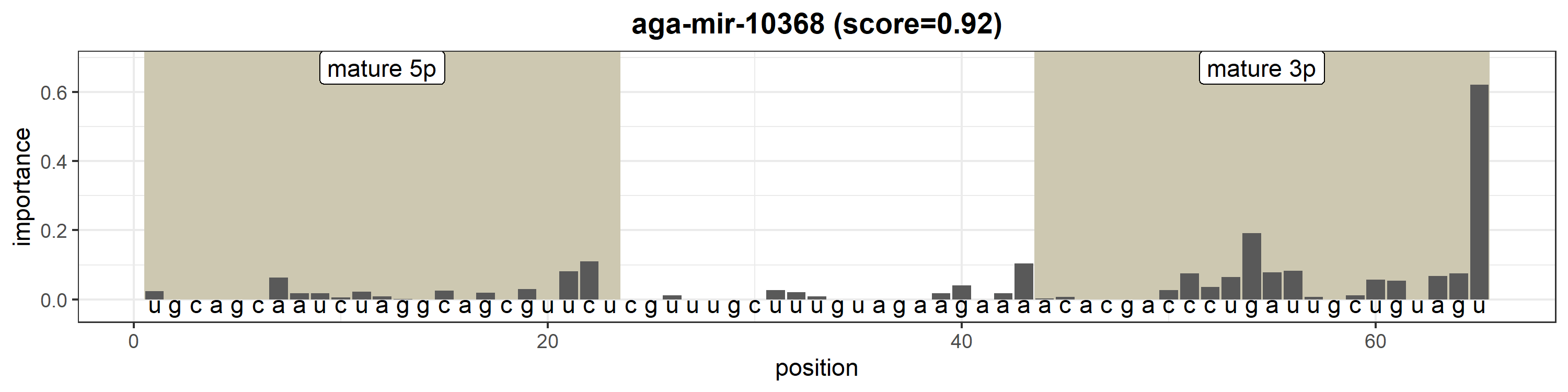

### aga-mir-10369.png

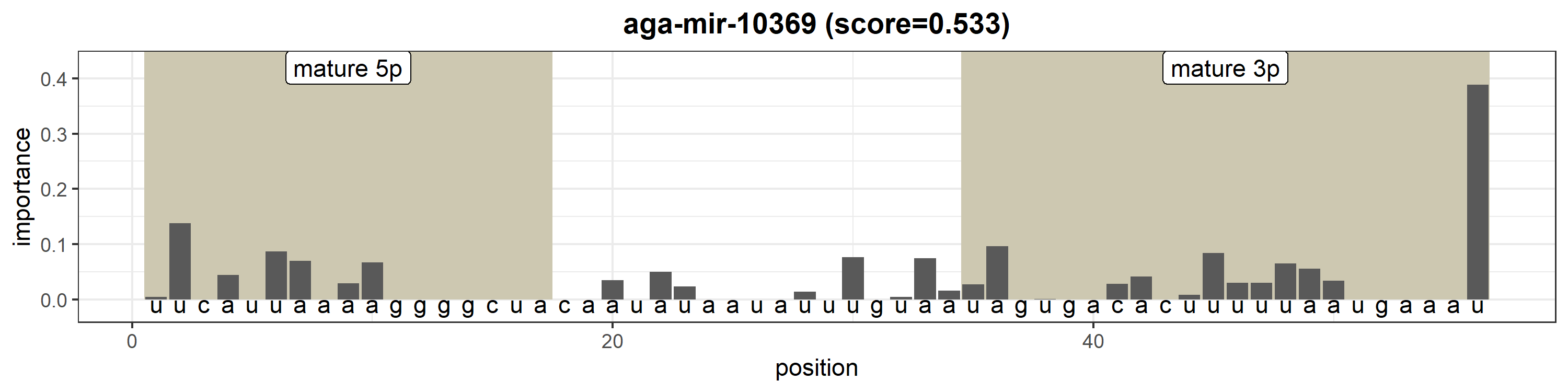

### aga-mir-10370-1.png

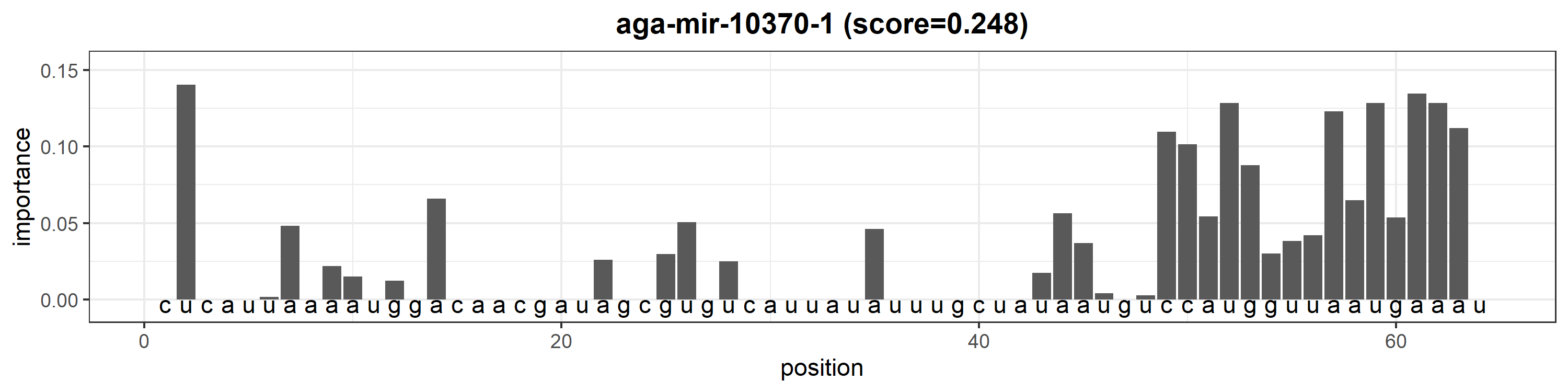

### aga-mir-10370-2.png

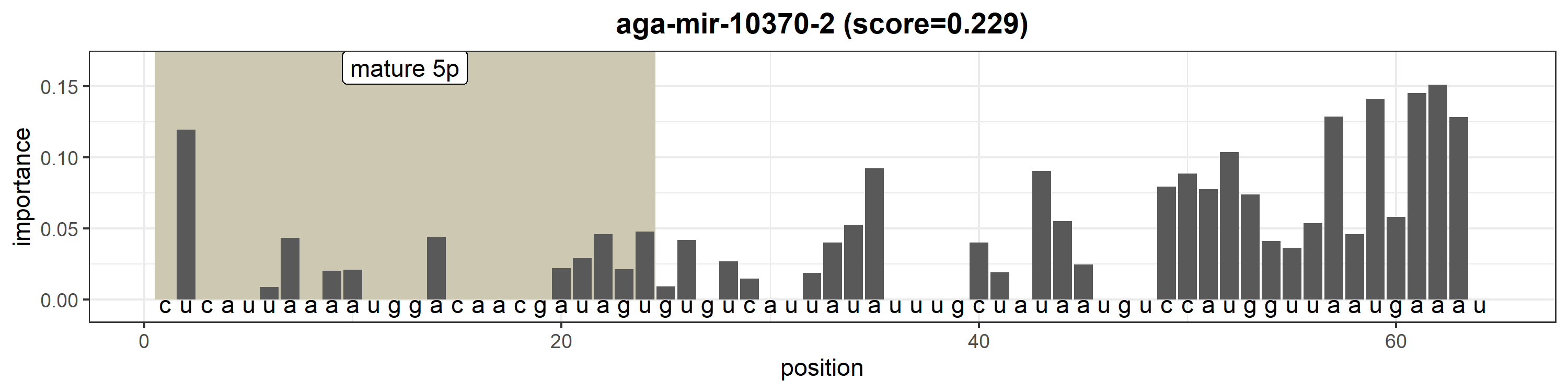

### aga-mir-10371.png

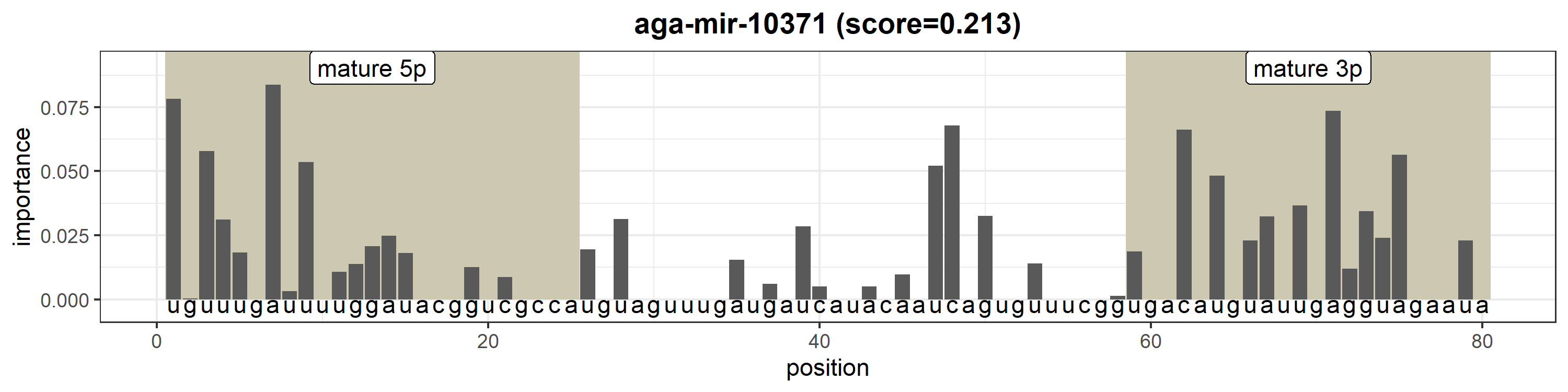

### aga-mir-10372a.png

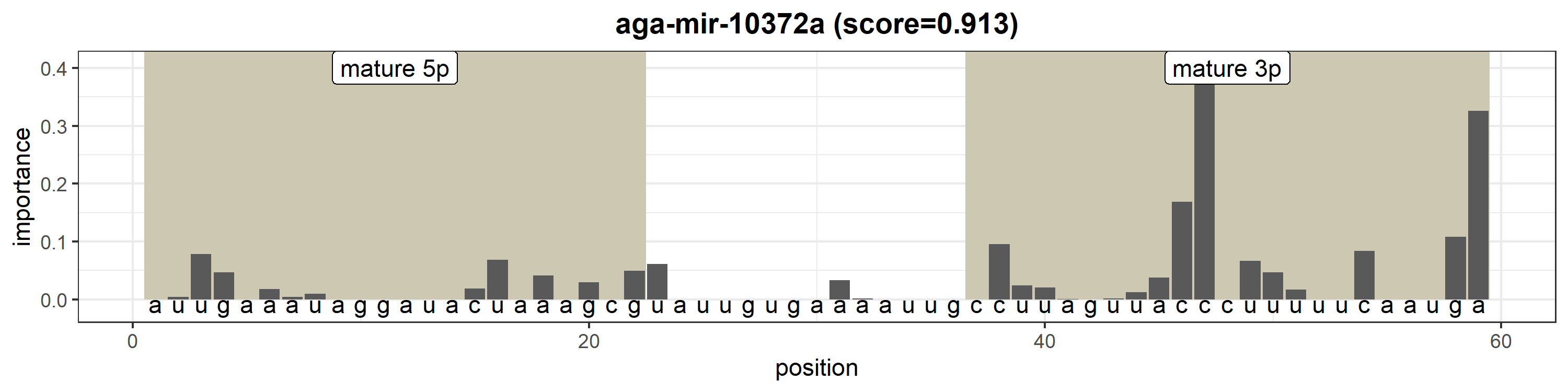

### aga-mir-10372b.png

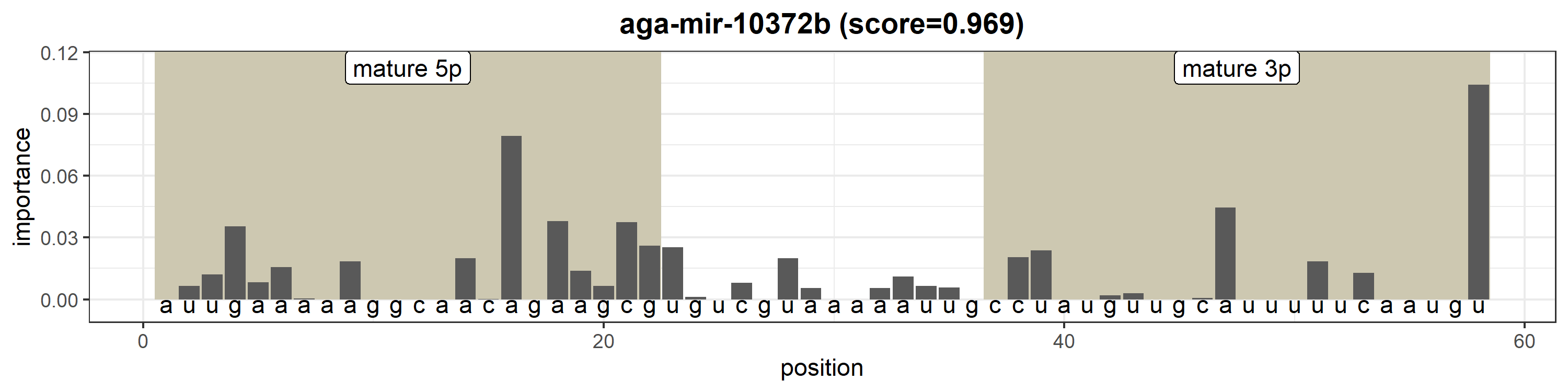

### aga-mir-10373.png

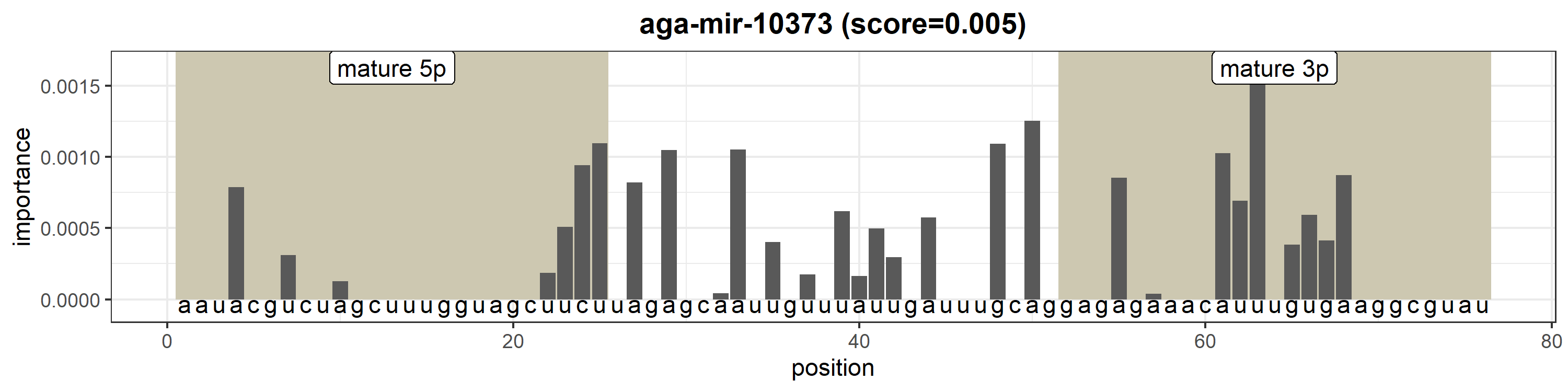

### aga-mir-10374.png

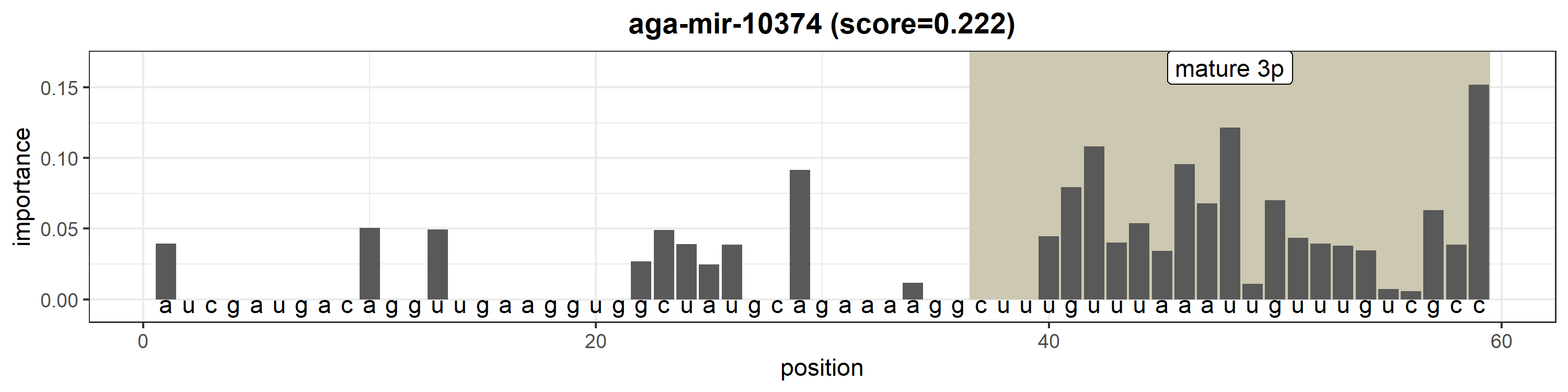

### aga-mir-10375a.png

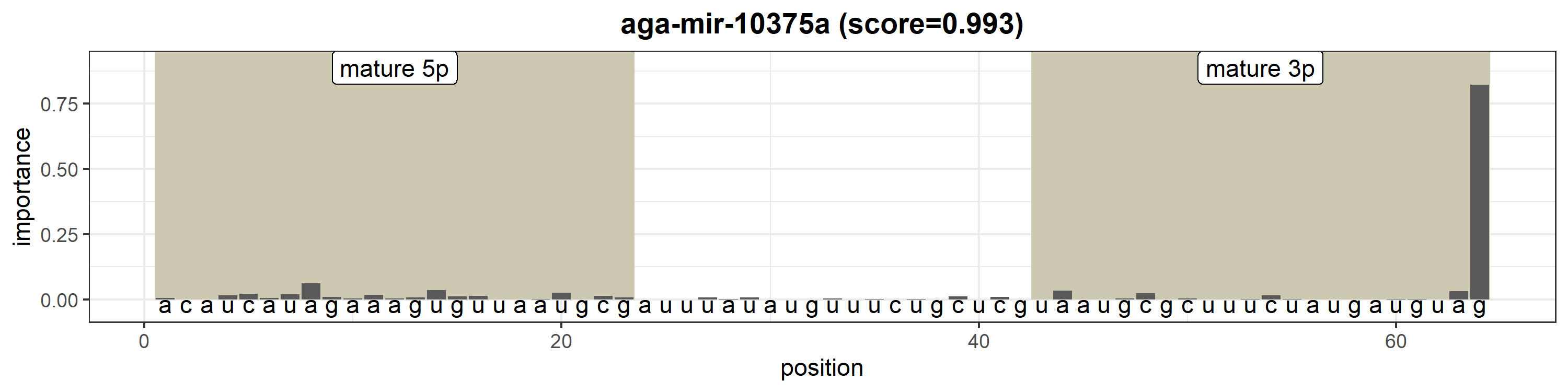

### aga-mir-10375b-1.png

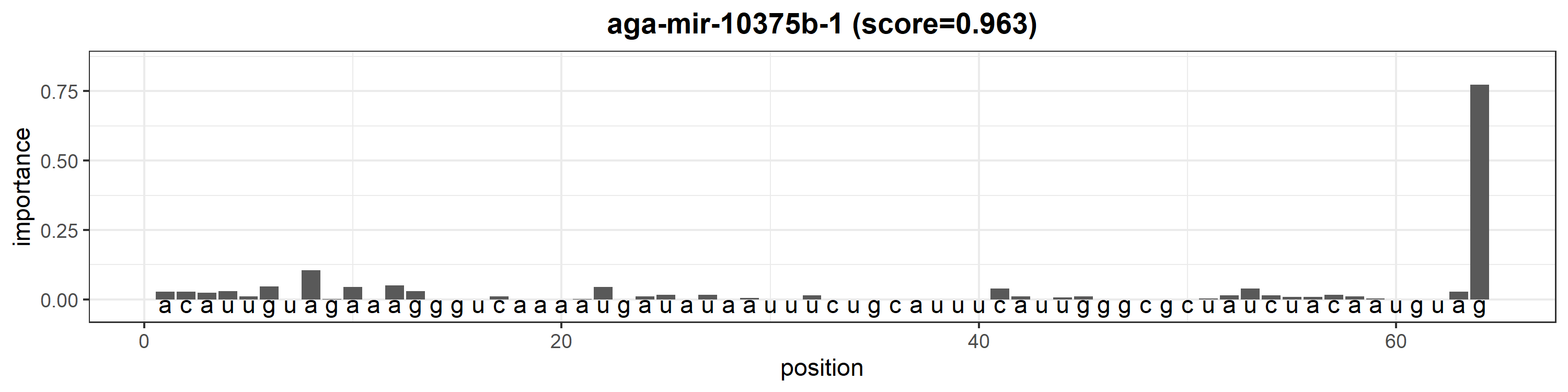

### aga-mir-10375b-2.png

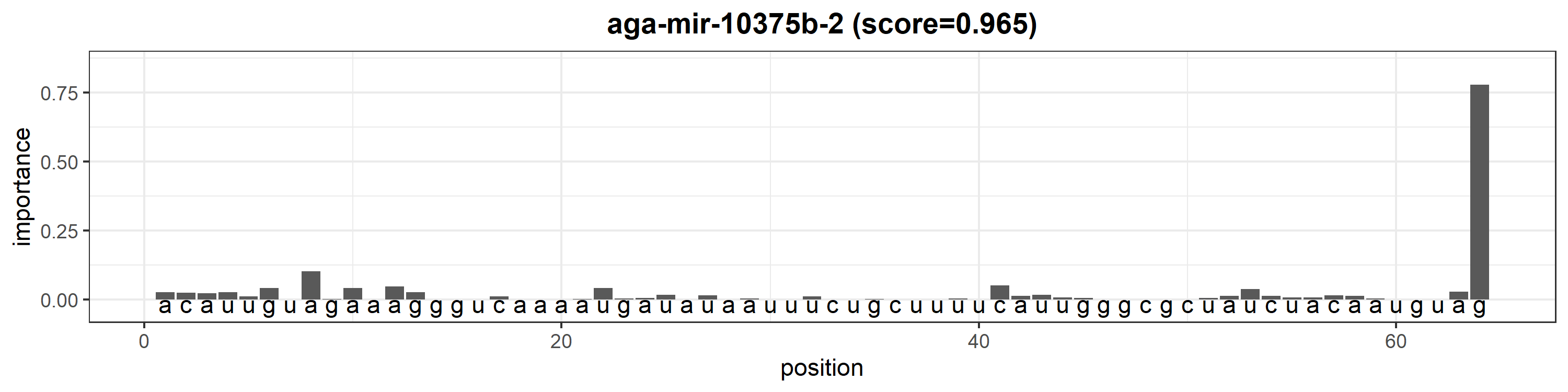

### aga-mir-10375b-3.png

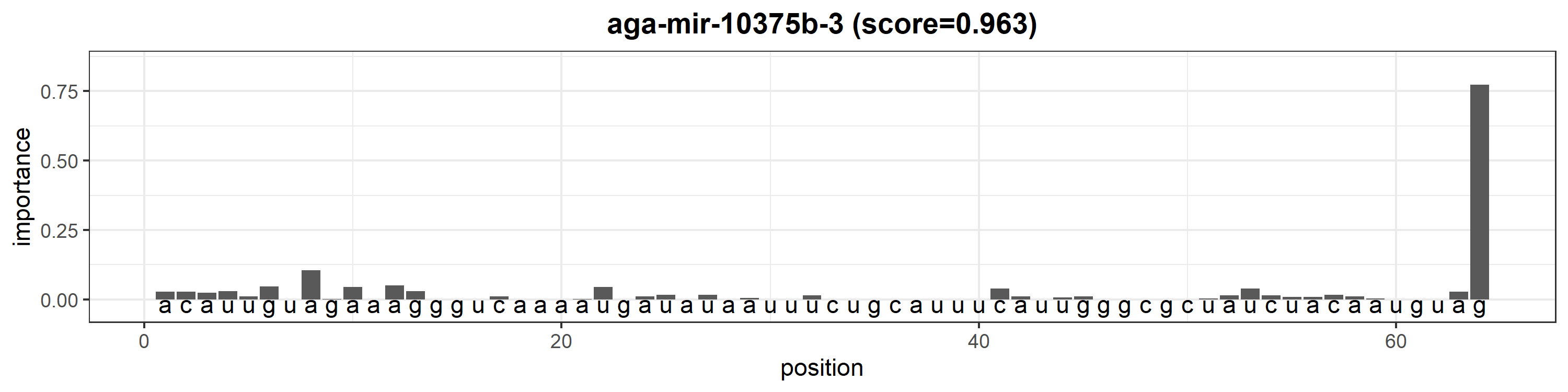

### aga-mir-10376.png

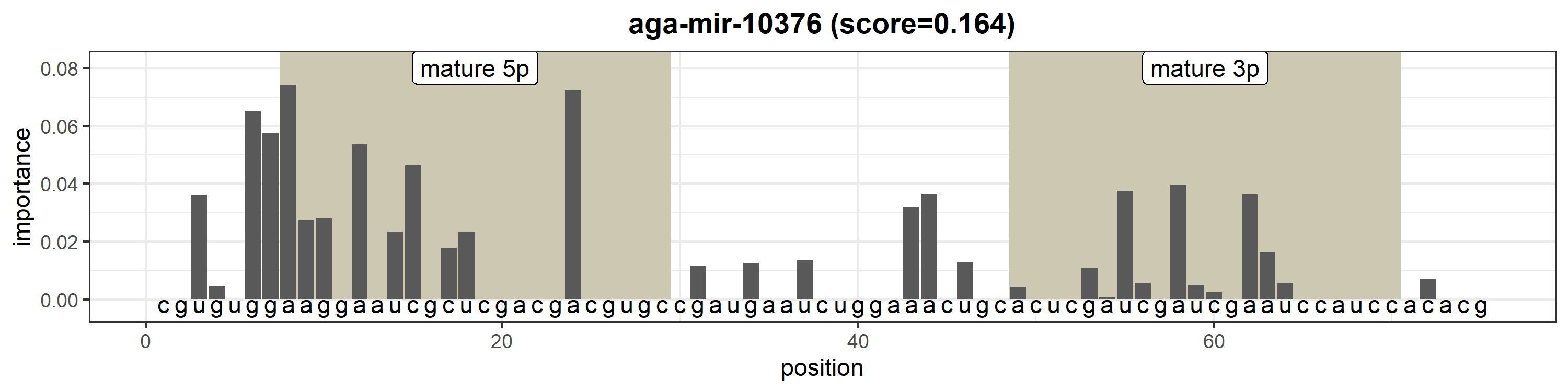

### aga-mir-10377a.png

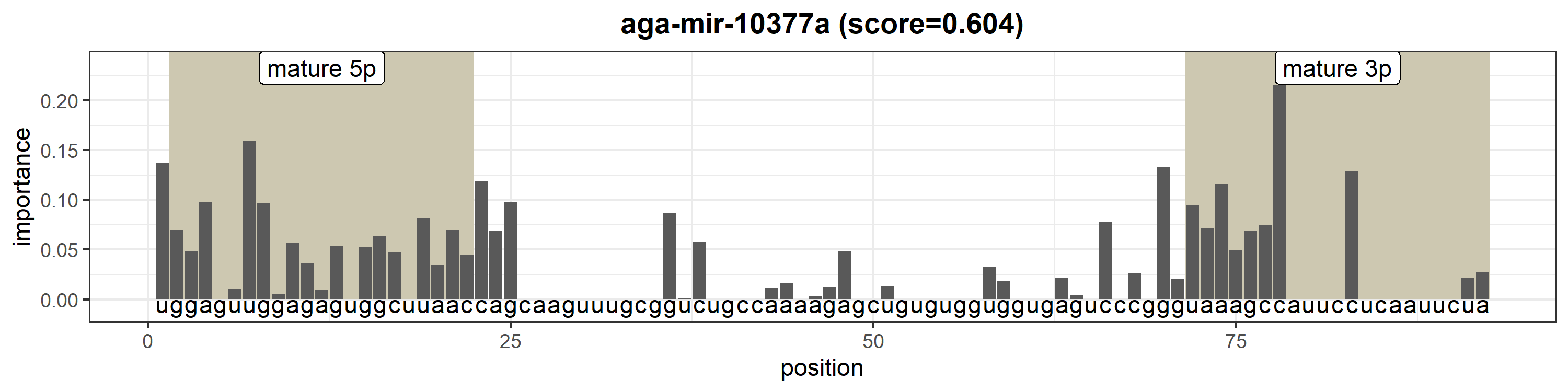

### aga-mir-10377b.png

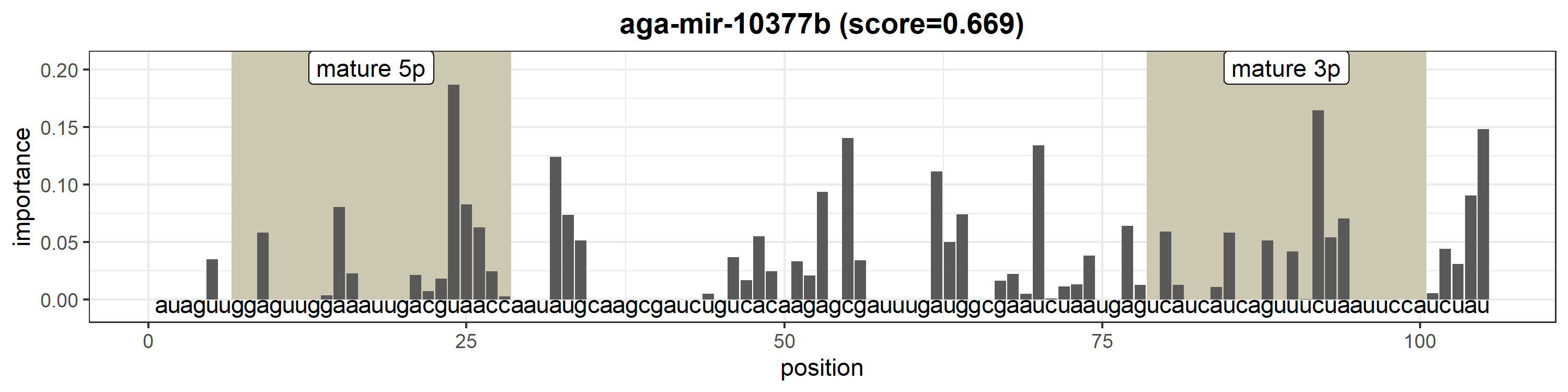

### aga-mir-10378.png

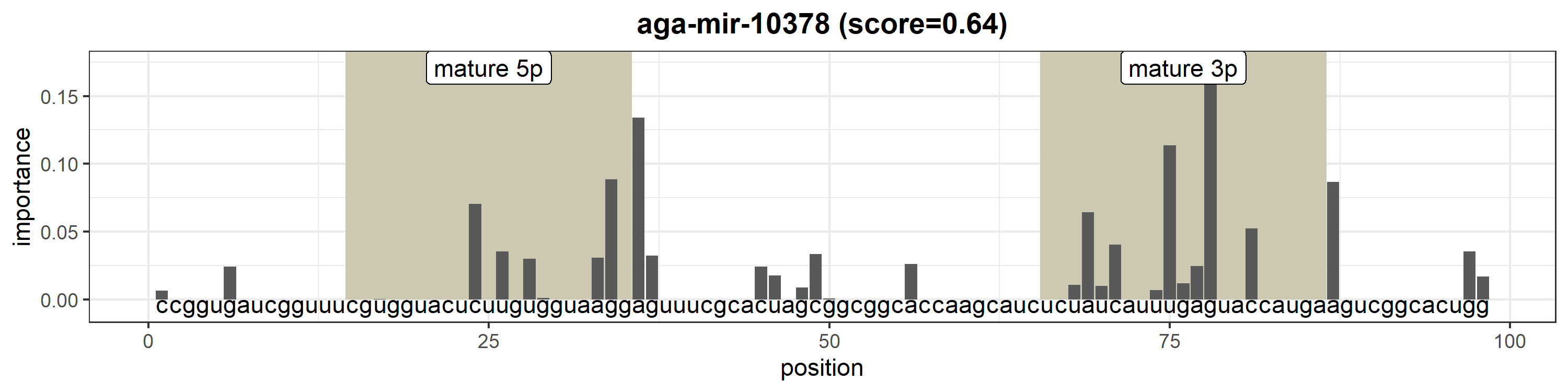

### aga-mir-10379.png

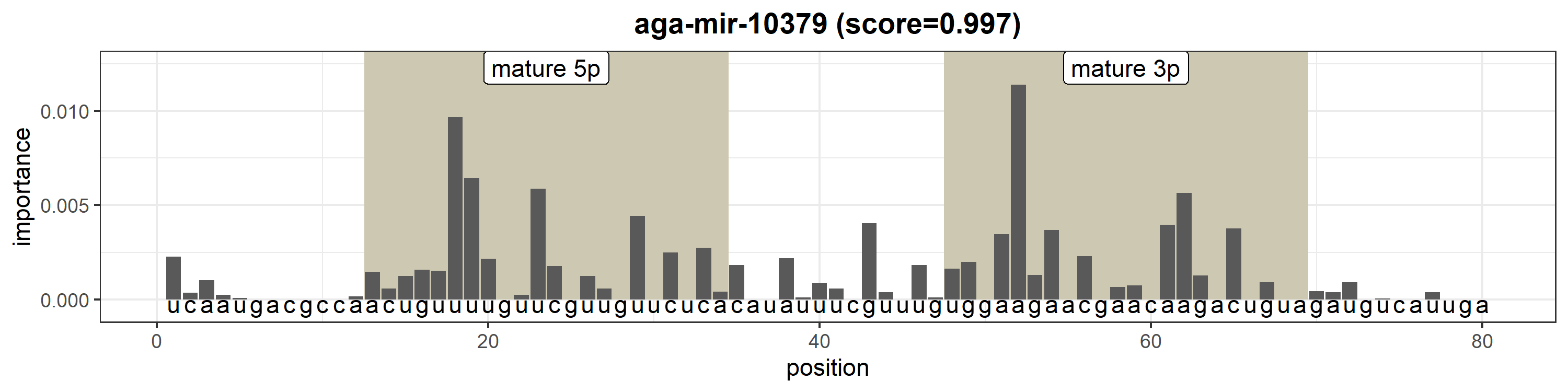

### aga-mir-10380.png

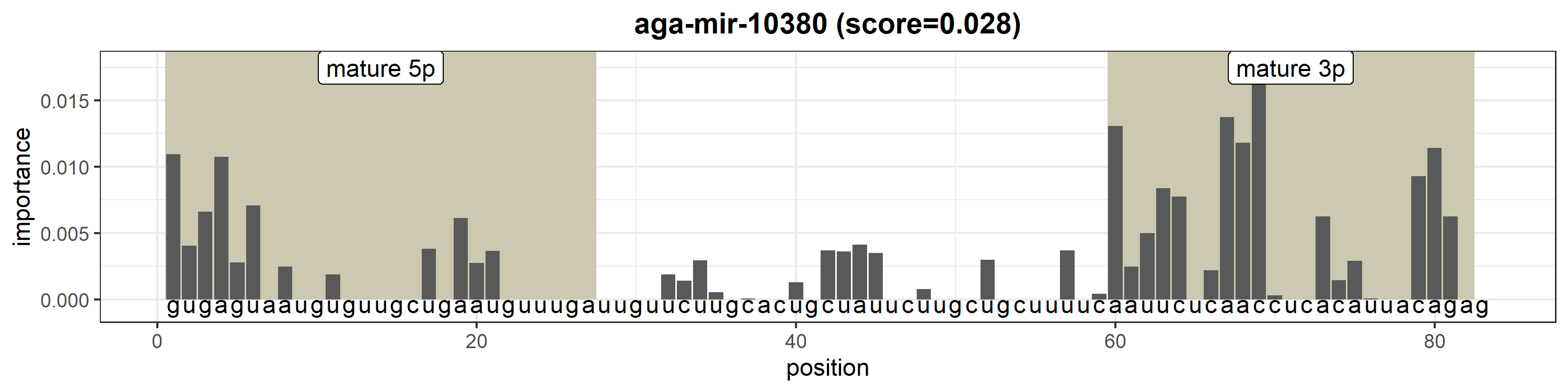

### aga-mir-10381.png

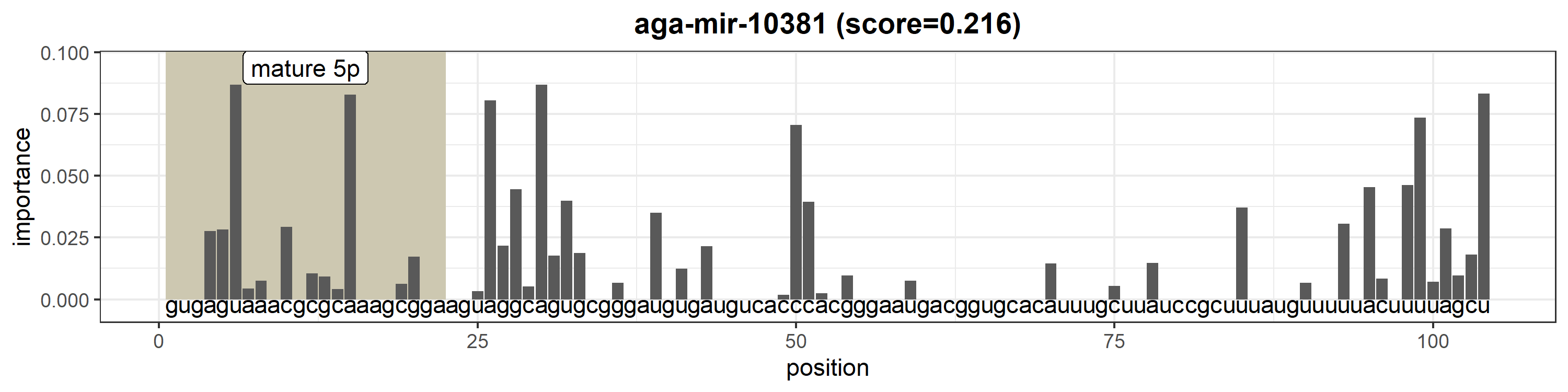

### aga-mir-12413.png

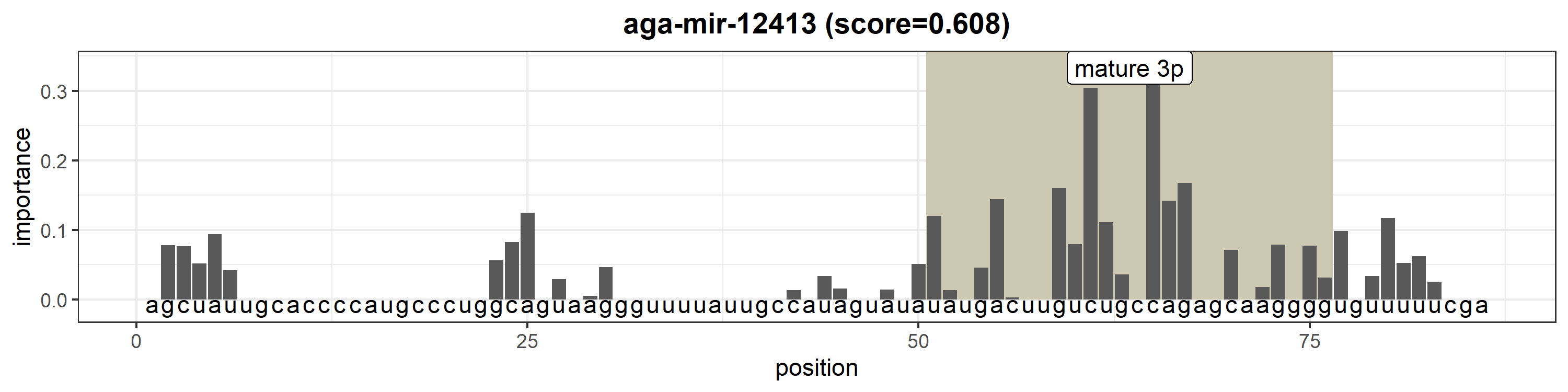

### aga-mir-12414.png

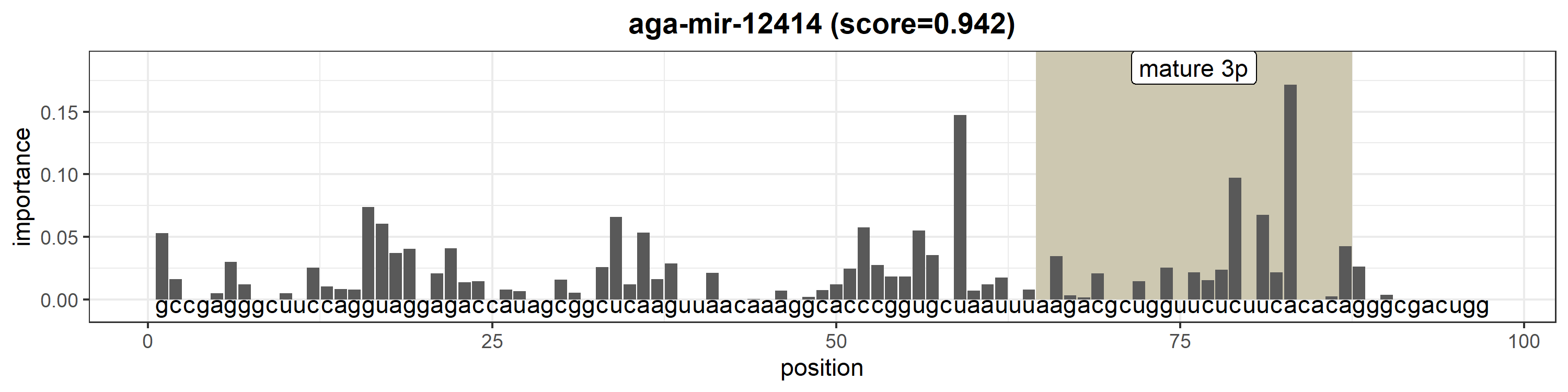

### aga-mir-12415.png

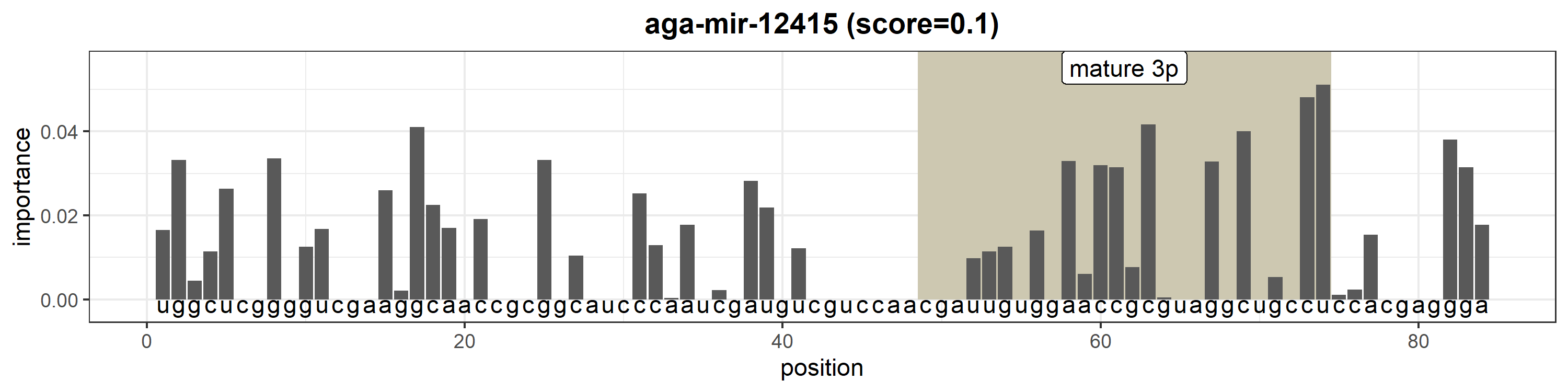

### aga-mir-12416.png

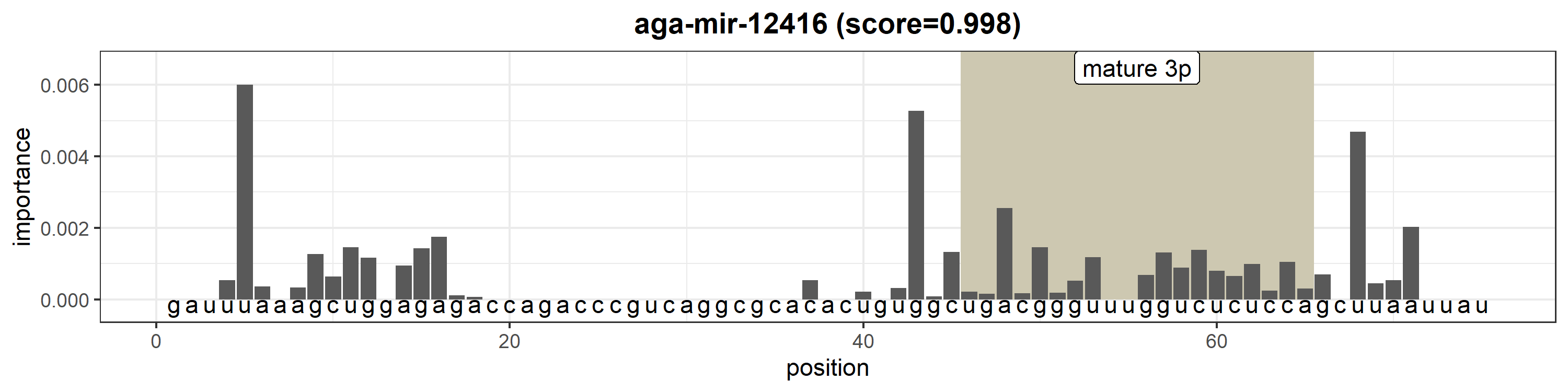

### aga-mir-12417.png

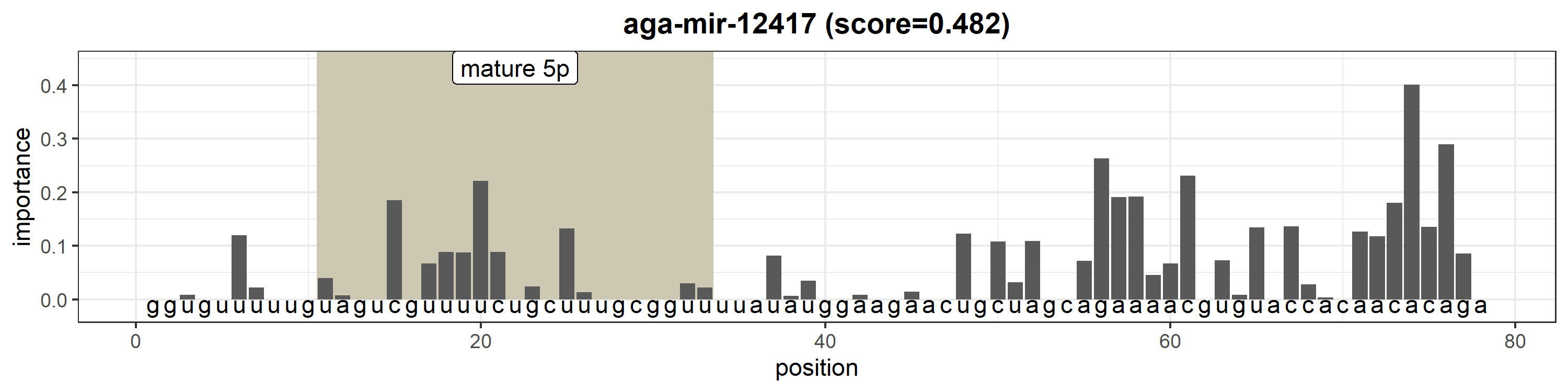

### aga-mir-12418.png

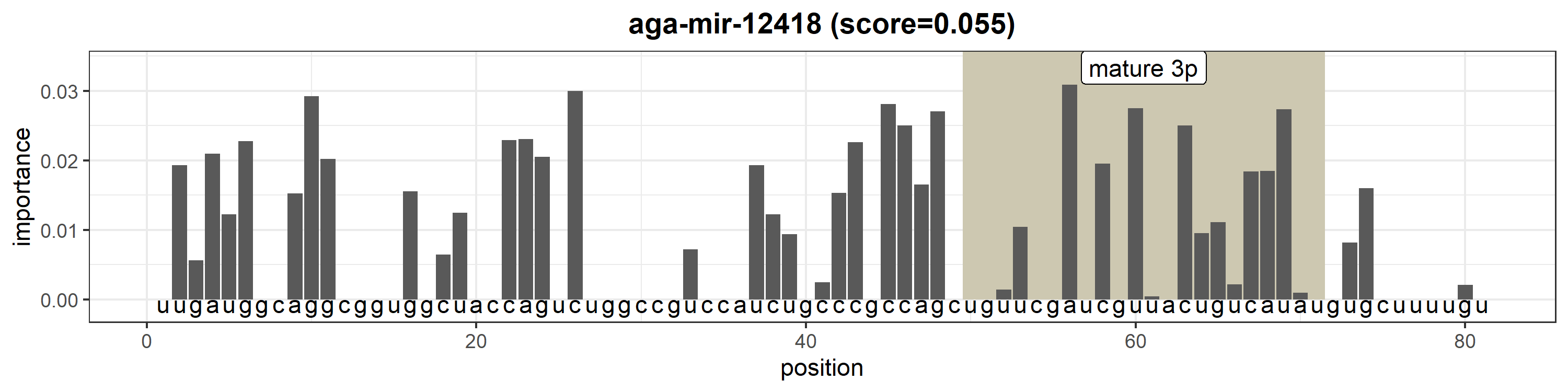

### aga-mir-12419.png

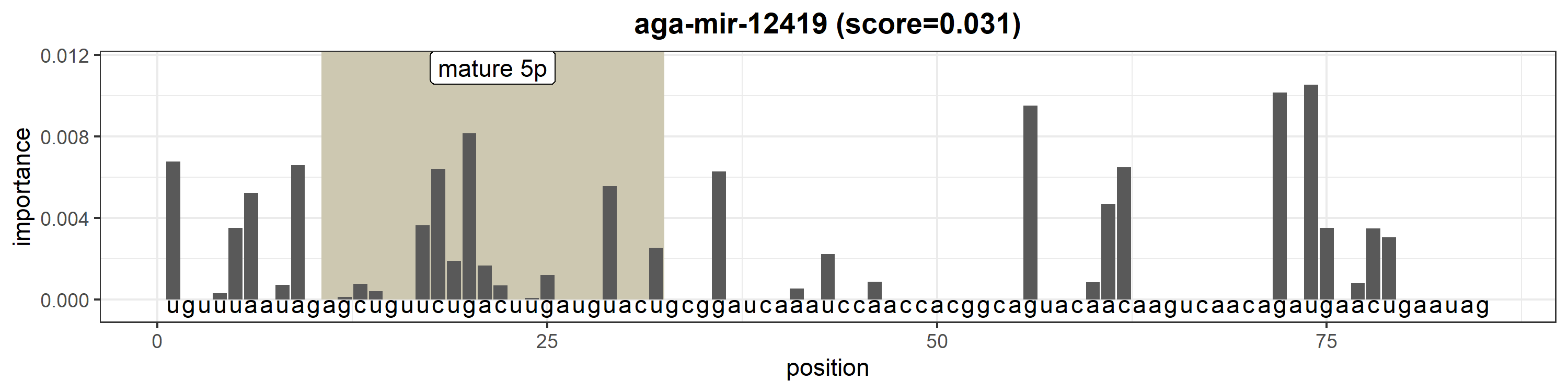
